# Functional genomic mechanisms of opioid action and opioid use disorder: a systematic review of animal models and human studies

**DOI:** 10.1101/2022.12.22.521548

**Authors:** Camille Falconnier, Alba Caparros-Roissard, Charles Decraene, Pierre-Eric Lutz

**Author notes:** Corresponding author: Pierre-Eric Lutz, INCI UPR 3212, 8 allée du général Rouvillois, 67000 Strasbourg, France.

## Abstract

In the past two decades, over-prescription of opioids for pain management has driven a steep increase in opioid use disorder (OUD) and death by overdose, exerting a dramatic toll on western countries. OUD is a chronic relapsing disease associated with a lifetime struggle to control drug consumption, suggesting that opioids trigger long-lasting brain adaptations, notably through functional genomic and epigenomic mechanisms. Current understanding of these processes, however, remain scarce, and have not been previously reviewed systematically. To do so, the goal of the present work was to synthesize current knowledge on genome-wide transcriptomic and epigenetic mechanisms of opioid action, in primate and rodent species. Using a prospectively registered methodology, comprehensive literature searches were completed in PubMed, Embase, and Web of Science. Of the 2709 articles identified, 73 met our inclusion criteria and were considered for qualitative analysis. Focusing on the 5 most studied nervous system structures (nucleus accumbens, frontal cortex, whole striatum, dorsal striatum, spinal cord; 44 articles), we also conducted a quantitative analysis of differentially expressed genes, in an effort to identify a putative core transcriptional signature of opioids. Only one gene, Cdkn1a, was consistently identified in eleven studies, and globally, our results unveil surprisingly low consistency across published work, even when considering most recent single-cell approaches. Analysis of putative sources of variability detected significant contributions from species, brain structure, duration of opioid exposure, strain, time-point of analysis, and batch effects, but not type of opioid. To go beyond those limitations, we leveraged threshold-free methods to illustrate how genome-wide comparisons may generate new findings and hypotheses. Finally, we discuss current methodological development in the field, and their implication for future research and, ultimately, better care.

## Introduction

Opioids have been used for millennia for their analgesic and euphoric properties[1]. While they remain reference pain treatments, their chronic use also associates with tolerance, physical dependence and, in some individuals, the emergence of an opioid use disorder (OUD). OUD is defined as a problematic use leading to significant impairment. This severe and chronic disorder associates with a life expectancy decreased by more than 10 years due to somatic and psychiatric comorbidities. Over the last 2 decades, western countries have faced an increase in death by opioid overdose, due to more frequent prescription for pain management, increasing use of illegal compounds, or misuse of substitution therapies. OUD imposes a major socio-economic burden, with an estimated annual cost in the trillion dollar range in the US[2], and increasing use and harm in Europe[3]. This results, overall, in a worsening public health crisis.

OUD results from interacting biological, psychological and socio-economic factors[4]. At a biological level, it originates from pharmacological effects of opioid drugs, which trigger chemical and molecular brain adaptations, under modulation by genetic vulnerability and epigenetic regulation. In turn, these effects mediate behavioral and cognitive dysfunctions, including the inability to control drug use despite harmful consequences, or persistent and intense desire for the drug, even after years of abstinence.

Since the 1990s, high-throughput approaches have been harnessed to characterize the molecular underpinnings of opioid effects in brain tissue, throughout the full genome[5], across various experimental models and species. In this context, the goal of the present systematic review was to synthesize current knowledge on functional genomic mechanisms of opioid action and OUD, defined as changes in gene expression or epigenetic regulation. To do so, we used a preregistered methodology, and performed an unbiased survey of bibliographic repositories, focusing on high-throughput studies, most notably microarrays and next-generation sequencing. While we acknowledge the contribution of candidate gene studies (already reviewed in the past[6, 7]), this focus on genome-wide analyses reflects the conviction that understanding heterogeneous phenotypes such as OUD requires analyzing the full genome.

Based on studies identified through this screening, a qualitative synthesis was first performed to summarize experimental designs and findings. Second, a focused quantitative analysis of bulk tissue transcriptomic studies was conducted, with the goal of defining a core signature of opioids. To our knowledge, while numerous reviews are regularly published in the opioid field[8–12], such a systematic synthesis across brain structures and technologies had never been performed. Our results indicate that, although more than 40 genome-wide studies have been published, available evidence for convergent findings is surprisingly limited. Third, we reviewed most recent work that leveraged cell-type specific and single-cell approaches, or that interrogated epigenetic regulatory mechanisms contributing to opioid plasticity. Finally, we discuss current challenges, as well as avenues and recommendations for future work.

## Methods

Full methodology and code are available as Supplementary Material and File.

## Results

The identification and selection of eligible studies is presented in Fig.1. For qualitative synthesis, 73 papers were selected, covering both transcriptomic and epigenomic approaches. Among these, the 44 articles that performed transcriptomic analyses in the 5 most frequently investigated regions were selected for more detailed quantitative synthesis: of these, 34 investigated a single brain region, 9 covered 2 regions, and only 1 characterized 5 regions[13]. This resulted in 52 differential expression analyses: 18, 11, 11, 6 and 6 in the nucleus accumbens (NAc), frontal cortex, whole striatum, dorsal striatum and spinal cord, respectively.

**Figure 1.**
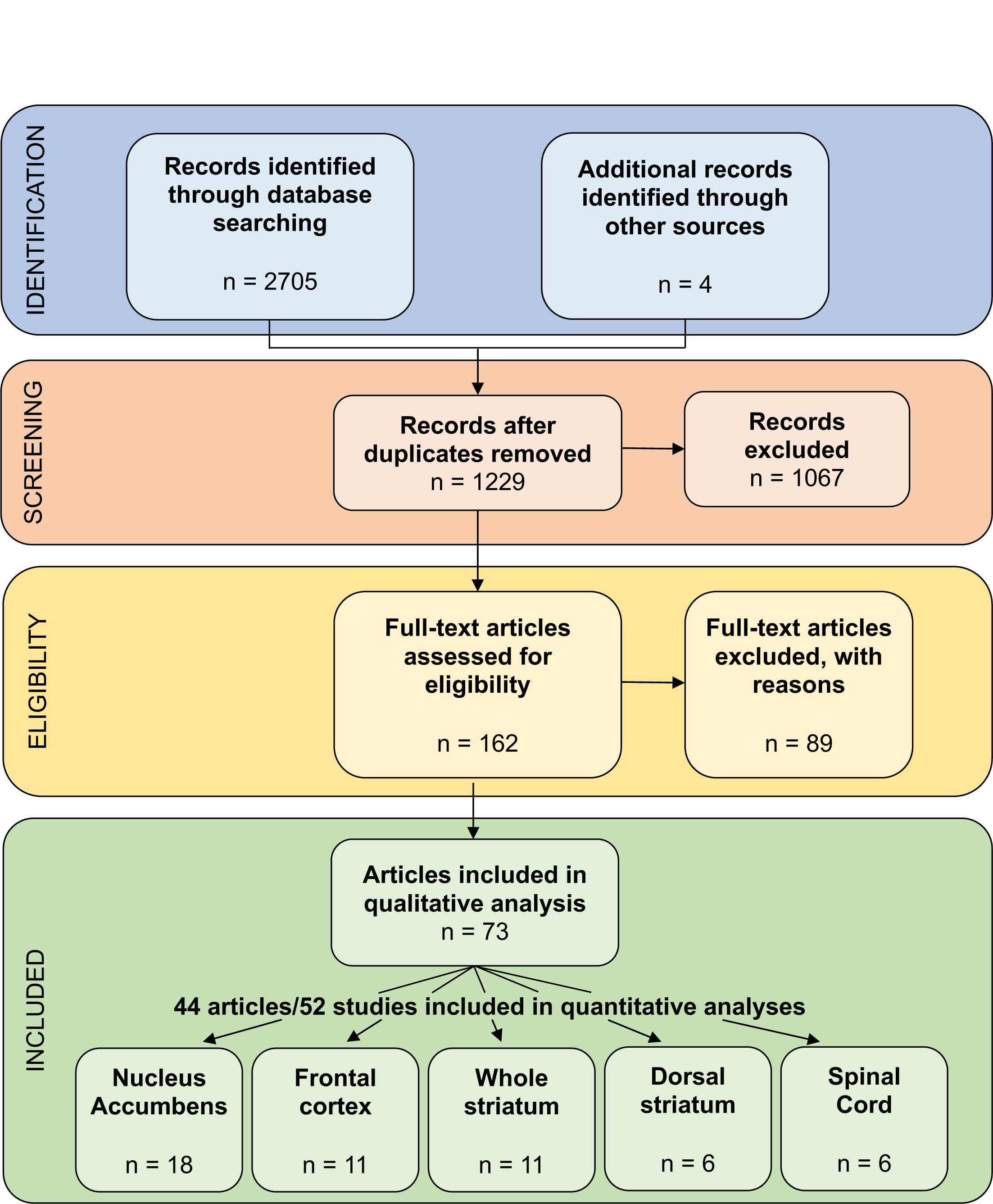
PRISMA diagram. The review protocol was registered prospectively in Prospero (record ID#CRD42022270113), and followed the Preferred Reporting Items for Systematic Review and Meta-Analysis protocol (PRISMA[190]). ***Identification of studies***. Three databases were used for systematic screening: MEDLINE, Embase and Web of Science. The literature search was performed using the following keywords: “microarray” OR “RNA sequencing” OR “bisulfite sequencing” OR “chromatin immunopurification sequencing” OR “single cell RNA sequencing” AND each of the following terms: “opiates”, “opioid”, “morphine”, “fentanyl”, “oxycodone”, “heroin”, “methadone” or “buprenorphine”. The initial search provided 2709 potentially eligible studies. ***Screening***. Articles were screened based on their title and abstract, and included if they used a high-throughput, genome-wide methodology to assess modifications of gene expression or epigenetic mechanisms as a function of opioid exposure, in the nervous system of primates or rodents (rat, mouse), and were published in English in peer-reviewed journals. Exclusion criteria included tissue other than the nervous system and candidate genes approaches. Once duplicates were removed, a total of 1229 articles were screened, among which 1067 were excluded because their title or abstract didn’t meet eligibility criteria. ***Eligibility***. After a more thorough examination of articles’ full-text, an additional 89 articles were excluded. As a result, 73 papers were selected for qualitative synthesis. Among these, 44 articles reporting on transcriptomic analyses in the 5 most frequently investigated regions were selected for a more detailed quantitative synthesis. Of these, 34 articles investigated a single brain region, 9 covered 2 regions, and only 1 characterized 5 regions [13]. Overall, this resulted in 52 differential expression analyses: 18, 11, 11, 6 and 6 in the nucleus accumbens (NAc), frontal cortex, whole striatum, dorsal striatum and spinal cord, respectively.

### 1. Qualitative analysis (n=73 studies)

#### Species and sex ratio

Most studies were conducted in rodents (Fig.2A): 30 (41%) in rats, 29 (40%) in mice, and 1 in both species. Only a small number investigated humans (n=11, 15%) and macaques (n=2, 3%), reflecting well-known practical limitations. As consequence, current knowledge on species-specific opioid effects is limited.

**Figure 2.**
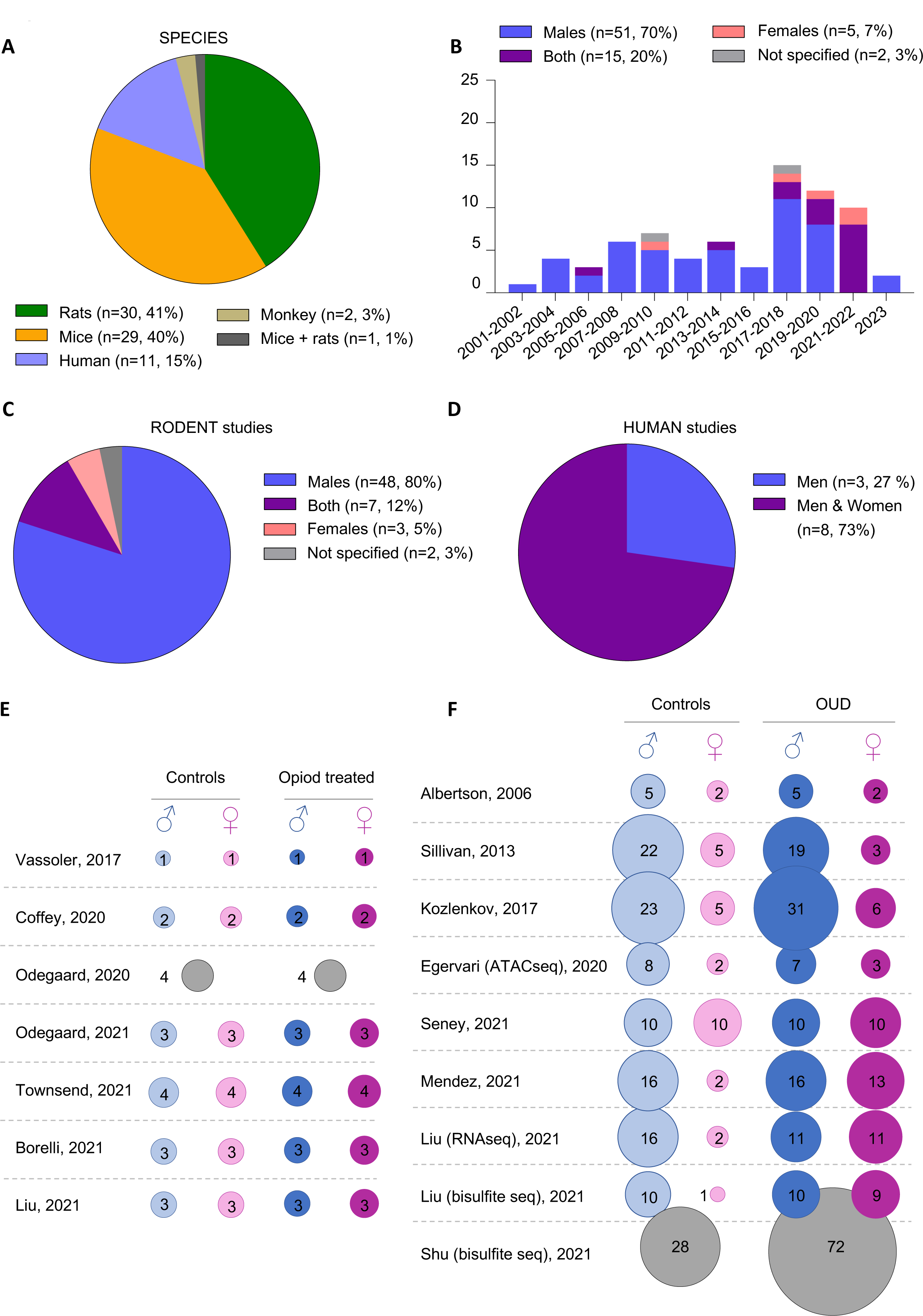
Characteristics of studies included: Species & Sex. **(A)** Species repartition in studies identified during systematic search. **(B)** Distribution of sex and year of publication across studies. **(C-D)** Sex distribution specifically in rodent and human studies. **(E-F)** Sample sizes in rodent and human studies that investigated both sexes and compared opioid-exposed animals to controls (rodents, panel E), or individuals with a diagnosis of opioid use disorder (OUD) to healthy controls (humans, panel F).

Regarding sex, epidemiological surveys indicate that OUD prevalence is higher in males than in females[3, 14] (although smaller differences have been reported with DSM-5 criteria[15]), with a number of sex-specific risk factors (e.g., ethnicity, income). This clinical situation has long been modeled in rodents, as sex differences in behavioral effects of opioids[16] have been described for analgesia[17], tolerance[18], withdrawal symptoms[19], reward[20] or motivation[21]. In comparison, molecular mechanisms underlying these differences are poorly characterized. The majority of OUD studies reviewed here investigated males only (n=51, 70%; Fig.2B-D), with few conducted in females only (n=5, 7%), and 15 (20%) that included both sexes (7 in rodents, 8 in humans). These numbers are consistent with the current landscape of neuroscience research, in which the ratio of articles reporting on males only, as opposed to females only, is around 5 to 1[22]. We nevertheless note an encouraging trend, as the majority of studies published since 2021 (8/12) investigated both sexes. Several factors likely underlie this evolution: policies from funding agencies (such as the NIH; https://grants.nih.gov/grants/guide/notice-files/not-od-15-102.html); increased recognition that females do not display more experimental variability than males[23]; and evidence that pathophysiological mechanisms may differ across sexes, as shown for depression[24], schizophrenia[25] or alcohol use disorder[26].

Specifically in rodents (Fig.2C), 80% of studies investigated males only (n=48). Seven articles investigated both sexes (n=7, 12%), using surprisingly low sample sizes (n=1-4 in male/female, and opioid-treated/control, groups; Fig.2D). Among these, 2 studies treated sex as a covariate[27, 28], and 3 pooled both sexes for the analysis of opioid effects[29–31], therefore providing no direct description of male/female differences. This is concerning, considering that the 2 remaining studies found little overlap, from 5 to 35% (see[32] and[33]), among morphine-induced changes occurring in each sex. Therefore, available evidence points toward substantial sex-specificity in opioid-induced transcriptomic plasticity, which will have to be confirmed in future work. In human, even less is known (Fig.2E). Among 11 articles, 8 investigated males and females, while 3 investigated males only. However, these 8 studies showed a strong bias towards examination of men (Fig.2F), with only one using similar sample sizes in both sexes[34]. In the latter, sex was treated as a covariate, with no specific description of its impact. Altogether, this significant gap in the literature calls for additional work, with the hope that better understanding of sex-specific pathophysiological routes may unravel distinct and more efficient therapeutic strategies.

#### Route and duration of opioid administration

Among studies included in qualitative analysis (n=73), the majority used chronic administration (n=59/73, 81%; see Fig.3A), mostly via self-administration (SA, n=22/59, 37%, including 11 human studies) and intraperitoneal injection (ip; n=19, 32%), followed by subcutaneous injection (n=8, 14%). Other procedures were less frequently used: continuous administration (with pellets or mini-pumps; n=5, 8%), intrathecal administration (n=3, 5%), or gavage (n=2, 3%).

**Figure 3.**
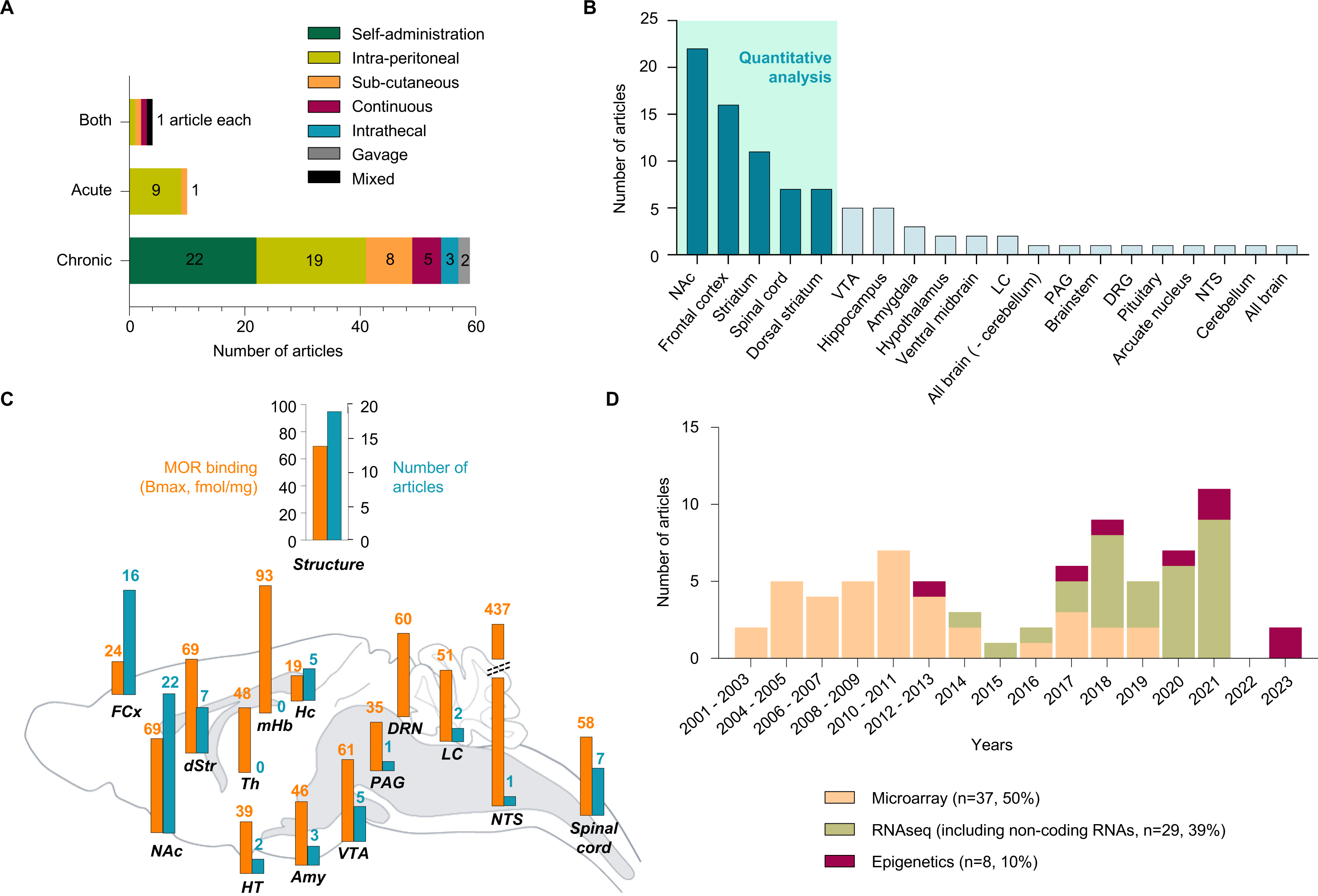
Characteristics of studies included: Injection parameters, Brain structure & Techniques. Among the 73 studies considered during qualitative analysis, a large panel of injection parameters, nervous system structures and techniques were represented. **(A)** Injection parameters used for opioid administration. NB: “mixed” refers to a single study that used a combination of chronic oral administration (through bottles) and acute ip injections of morphine[79]. **(B)** Number of articles investigating each nervous system structure. **(C)** Side-by-side comparison of the number of articles investigating each structure with the protein expression profile of the molecular target of opioids, the mu opioid receptor (MOR; as assessed using [3H]-DAMGO autoradiography, data in fmol/mg of tissue, adapted from[115, 118–120]). **(D)** Distribution of high-throughput methodologies and year of publication across studies. *Abbreviations:* FCx, Frontal cortex; NAc, Nucleus accumbens; dStr, Dorsal Striatum; HT, Hypothalamus; Th, Thalamus; mHb, Medial Habenula; Hc, Hippocampus; Amy, Amygdala; VTA, Ventral Tegmental Area; PAG, periaqueductal gray; DRN, Dorsal Raphe Nucleus; LC, Locus Coeruleus; NTS, Nucleus Tractus Solitarius.

There is limited information regarding differences among administration procedures. Interestingly, Lefevre et al[35] recently compared 3 conditions: continuous and systemic morphine infusion using osmotic mini-pumps, with (“Interrupted”) or without (“Continuous”) the induction, twice daily, of a precipitated withdrawal episode (using naloxone, a non-specific opioid antagonist), compared to Controls (saline mini-pumps). Results indicated 687 and 407 differentially expressed genes (DEG) when comparing Interrupted and Control groups in the NAc and dorsal striatum, respectively, with only 1 DEG between Continuous and Control groups. Therefore, rather than opioid exposure per se, it is the repetition of intermittent withdrawal episodes that may drive the intensity of transcriptomic adaptations affecting the mesolimbic system (whether precipitated, as in this study, or spontaneous, in-between bolus injections). This is consistent with the clinical notion that the alternation of intoxication and withdrawal phases is critical for the development of OUD.

Opioid effects have also been investigated following acute administration (10/73 studies, 14%), corresponding mostly to ip injection (9 articles, with only 1 using subcutaneous injection). While these studies were interested in the use of opioids as analgesics or anesthetics[36, 37], or compared acute effects of various psychoactive drugs[38], their results may be relevant for the understanding of OUD. Indeed, comparisons of inbred mouse strains found striking variability in the intensity of opioid physical dependence, with the same strains showing severe or mild physical signs when withdrawal was precipitated after acute or chronic opioid injections[39, 40]. In other words, there may be shared genetic determinants, within each strain, for brain adaptations triggered by acute or chronic opioid exposure. A recent single-cell RNA-sequencing study (scRNA-Seq) provides an interesting illustration of how such strain-specific genetic traits may translate into at least partially similar transcriptional plasticity during acute or chronic morphine exposure ([41] and *3. High cellular resolution approaches* below).

Another important consideration is that most studies (51/73, 70%) used passive administration, which do not model volitional or compulsive drug-taking. This reflects the technical difficulty of intravenous SA experiments (IVSA), particularly in the mouse. Few studies directly compared passive and active exposure. Among these, Tapocik et al[42] analyzed “yoker” mice and groups of “yoked-morphine” or “yoked-saline” animals which, each time the yoker self-administered morphine, were passively exposed to morphine or saline IV injections. In C57BL/6J, much more DEG were identified between the yoker and yoked-saline groups (n=1176) than between yoked-morphine and yoked-saline (n=244), with their overlap likely reflecting passive morphine effects (n=103). In DBA/2J mice, which did not develop morphine self-administration (as shown by the absence of differences in operant responding among the 3 groups), transcriptional effects were milder, as expected (yoked-morphine *vs* yoked-saline, n=262 DEG; yoker *vs* yoked-saline, n=107). This suggests that distinct transcriptional regulations may mediate voluntary opioid consumption and reflect their pharmacological effects, with both aspects ultimately contributing to OUD.

New paradigms have been developed to facilitate such studies, which notably include oral SA[43–45]), or devices for opioid vapor SA[46]. In addition, optogenetic tools now enable light-induced self-stimulation (optogenetic intracranial self-stimulation, oICSS)[47]. The latter models are based on selective manipulation, using opsins, of the activity of anatomically- or genetically-defined neuronal populations. While these approaches do not recapitulate the systemic effects of opioid IVSA, they are nevertheless able to trigger similar behavioral deficits, including uncontrolled consumption, resistance to punishment, or excessive motivation[48, 49]. It is therefore possible that they may allow dissecting the neuronal pathways that are necessary or sufficient for each of these behavioral dimensions, as well as underlying molecular mechanisms. Another avenue for improvement relates to the mode of action of genetic tools used for oICSS. Upon light-stimulation, opsins open ion channels that modulate neuronal excitability. Their high temporal resolution does not necessarily represent the best model for the longer time-frame of pharmacological action of drugs of abuse. To more faithfully mimic such kinetics and associated intracellular signaling, chemogenetics or chimeric opsins (for light-induced metabotropic signaling, with a proof-of-concept available for MOR[50]) might represent better tools.

#### Nervous system structures

Among articles eligible for qualitative analysis, 18 regions of the nervous system were explored (Fig.2B). Unsurprisingly, a large majority focused on the mesocorticolimbic dopaminergic pathway: the NAc (22 articles[13, 27, 30, 33–35, 37, 51–65]), frontal cortex (16; [13, 29, 32, 34, 62, 66–76]), whole/unspecified striatal complex (11; [31, 38, 77–85]), dorsal striatum (7; [13, 35, 59–61, 86, 87]) and ventral tegmental area (VTA, 5; [33, 62, 88–90]). Other regions included the spinal cord (7; [77, 91–96]); hippocampus (5; [13, 78,97–99]); amygdala (3; [13, 100, 101]); locus coeruleus[89, 102], ventral midbrain[42, 103], hypothalamus[104, 105] (2 each), whole brain with[106] or without[107] cerebellum, periaqueductal gray matter[108], pituitary gland[105], arcuate nucleus[109], nucleus of the tractus solitarius[36], brainstem[28], cerebellum[99] or dorsal root ganglia[110] (DRG, 1 each). Below, we briefly describe the rationale for studying such diverse structures.

The course of OUD involves 3 stages: intoxication, withdrawal/negative affect, and anticipation of next intoxication. These stages reflect gradual adaptation of the brain to drug exposure, and rely on distinct mechanisms. The intoxication stage, mostly driven by acute reward, notably associates with dopamine release, in the striatum and frontal cortex, by neurons located in the VTA. The second stage corresponds to negative reinforcement, whereby drug consumption alleviates physical and negative affective states of withdrawal, involving the locus coeruleus[111], NAc[112], and amygdala[113], among others. Finally, anticipation results from memories of drug-associated cues and contexts (hippocampus), as well as impaired goal-directed behaviors, implicating higher-order structures (e.g. prefrontal and orbital cortices[114]). These relationships are not exclusive, with some structures implicated at multiple stages (e.g., the striatum, also involved in habit formation). Other regions (spinal cord, DRG, brainstem, periaqueductal gray matter, PAG) were investigated in relation to opioid analgesia, tolerance and hyperalgesia. Finally, few articles were interested in how opioids modulate neuroendocrine systems (hypothalamus, arcuate nucleus, pituitary gland).

In addition to these associations, it is also important to consider the distribution of the endogenous opioid system. This system is composed of 3 opioid receptors (mu, delta and kappa; MOR, DOR and KOR), among which the MOR is necessary for opioid-induced analgesia, reward, and physical dependence[115]. Neurons expressing this receptor therefore represent pharmacological entry points for brain adaptations to opioids. Comparison of studies reviewed here with MOR brain distribution (assessed using ligand autoradiography[116–121]; Fig.3C) indicates that: i) MOR is highly expressed among structures most frequently studied, as expected; ii) surprisingly, other areas with similarly high expression have received little to no attention (NTS, locus coeruleus, dorsal raphe nucleus, medial habenula, thalamus), calling for more work.

#### Techniques

Thirty-seven and 29 studies used microarrays and RNA-seq, respectively, to analyze the transcriptome (Fig.3D). While the first RNA-seq study was published in 2014[37], this technique has since progressively replaced microarrays, reflecting well-recognized advantages (higher sensitivity and throughput, gene discovery). For epigenomic profiling, 3 studies used DNA methylation arrays[73, 75, 76], 2 bisulfite sequencing[72, 74], and 1 ATACseq (Assay for Transposase-Accessible Chromatin with sequencing)[87].

### 2. Quantitative analysis of transcriptomics studies

To identify reproducible transcriptomic opioid signatures, we ran quantitative analyses of bulk-tissue results available for the 5 most studied structures (44 articles; Fig.4A). To do so, gene-level results were retrieved when available. This yielded 24 lists of DEG comparing opioid-treated animals and control groups, originating from 17 articles. Among these, 6 articles did not report any p-values, while the remaining 11 used very diverse thresholds on p-values and fold changes (FC; Supplementary Table 1). Only a small minority (3/44, 6.8%[31, 33, 34]) provided p-values and FC for all genes. In addition, raw data were available in the public repository GEO for less than half of the studies that reported DEG (8/17, 47%). Overall, this underscores strong heterogeneity in data analysis and reporting, which limits re-use[122]. To counter such issues, efforts are being made to harmonize practices[123, 124], with guidelines enforced by funding agencies[125]. For the present review, a genome-wide meta-analysis combining effect and sample sizes across studies was not possible. We nevertheless: i) Systematically assessed concordance among available lists of DEG; ii) Conducted a threshold-free comparison of the 2 NAc studies for which genome-wide results were available.

**Figure 4.**
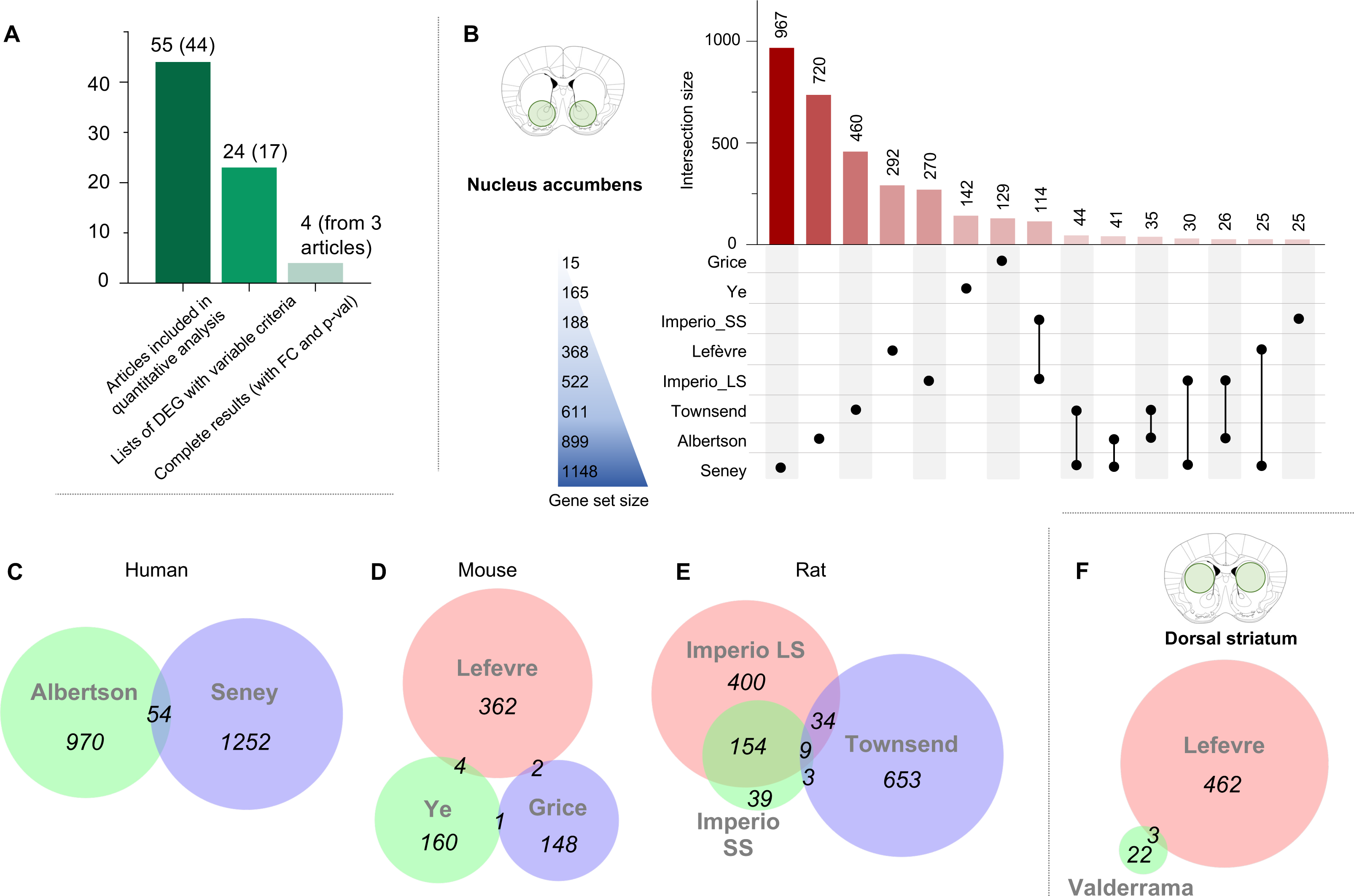
Data availability for eligible studies and quantitative analysis in the nucleus accumbens. **(A)** The graph depicts the number of: i) differential expression analysis results considered for quantitative analysis (with the corresponding number of articles between brackets); ii) cases when lists of differentially expressed genes (DEG) were available; iii) cases when full genome-wide results were available (with both fold changes, FC, and p-values). **(B)** Number of DEG identified in studies that investigated the nucleus accumbens. **(C-E)** Venn diagram of DEG identified in each species (human, mice and rat). **(F)** Venn diagram of DEG in the 2 studies that investigated the dorsal striatum.

#### 2.1 Nucleus accumbens

Among 18 articles, only 8 provided lists of DEGs: 4 in mice (males only[35, 37, 51, 54]), 2 in rats (1 in males[56], 1 in both sexes[33]), 2 in humans (both sexes[34, 58]). The 8 articles provided a total of 3502 DEG (Fig.4B-E). Among these, the vast majority were identified by 1 study only (n=3017/3502, 86%), while 424, 57 and 4 DEG were identified across 2, 3 or 4 studies, respectively. No single genes were common to 5 or more articles. In addition, the best overlap was found among the 2 DEG lists reported in a single study by Imperio et al[56] (Fig.3E). Overall, this indicates low concordance among NAc studies.

To define factors contributing to this heterogeneity, we intersected DEG lists in each species separately (Fig.3C-E). This, however, did not improve concordance: in humans, only 54 genes were common to both studies (Jaccard Index, JI=2.4%, Fig.4C, Supplementary Table 2 for the matrix of pair-wise JI across all studies); in mice, 7 were identified by at least 2 studies (JI=1%, Fig. 4D); in rats, the JI increased to 15.5% (Fig.4E), mostly driven by high overlap among the 2 aforementioned DEG lists from Imperio et al. In the latter work, the authors identified large-(LS) and small-suppressors (SS): in LS rats, greater avoidance of a natural reward (saccharine) that predicted heroin availability was associated, compared to SS rats, with worse outcomes during later heroin IVSA (increased consumption, higher motivation and relapse). Gene expression changes occurring in the LS group (compared to control rats given access to saccharine but not heroin; n=597 genes), and in the SS group (same controls, n=205) showed a JI of 25.5% (with striking concordance in up-and down-regulation). This was much higher than overlap between Imperio et al and the other rat study, by Townsend et al (Imperio-SS/Townsend, JI=1.35%; Imperio-LS/Townsend, 3.43%), suggesting that differences across laboratories and animal facilities may significantly contribute to poor concordance. Importantly, while the relatively small number of studies in the NAc precluded definitive conclusions, a more powerful analysis of sources of variability was possible across our 5 regions of interest (see section *2.6*).

#### 2.2 Dorsal striatum

Only 2 dorsal striatum studies provided DEG (Fig.3F). One study used chronic morphine administration interrupted by naloxone injections in male mice[35], as described above, while the other used acute morphine injection in male rats[86]. Reflecting these differences, a very small overlap was found between the 2 (0.62%). Therefore, while the dorsal striatum is thought to gradually gain control over drug-taking in OUD[126], underlying transcriptomic plasticity is poorly characterized.

#### 2.3 Whole striatum

Six of the 11 whole striatum studies provided DEG[31, 38, 79–82] (Fig.5B): 2 investigated chronic heroin[82] or morphine[31], while 2 compared acute effects of distinct opioids (morphine and heroin)[38, 80], and 2 compared acute and chronic morphine[79, 81]. Most DEG were identified by 1 study only (3308/3736, 88.5%), with only 112 genes common to 3 or more studies. Consistent with NAc results, strongest overlaps were observed for lists of DEG coming from the same article, with for example 54 (JI=17%, Korostynski 2013[38]) and 16 (21%, Piechota 2010[80]) DEG common to acute morphine and heroin injections, respectively, and 139 DEG (31%, Korostynski 2007 [81]; Fig.5A) common to acute and chronic morphine administration. In sharp contrast, comparisons across, rather than within, studies yielded poor overlaps, even when considering those that used similar drug administration: among chronic morphine studies (Fig.S1A), the JI varied from 0.1 to 2.6% (Korostynski 2007/Skupio 2017), and, among acute morphine studies, from 2.7% to 7.8% (Fig.S1B), with only one comparison standing out among acute heroin studies (30 common DEG, JI=19%, Piechota 2010/Korostinsksy 2013). Importantly, 5 of these 6 studies in the whole striatum were conducted by the same group, suggesting that significant variability may persist even among studies conducted in the same facility, but at different periods (see also section 2.6).

**Figure 5.**
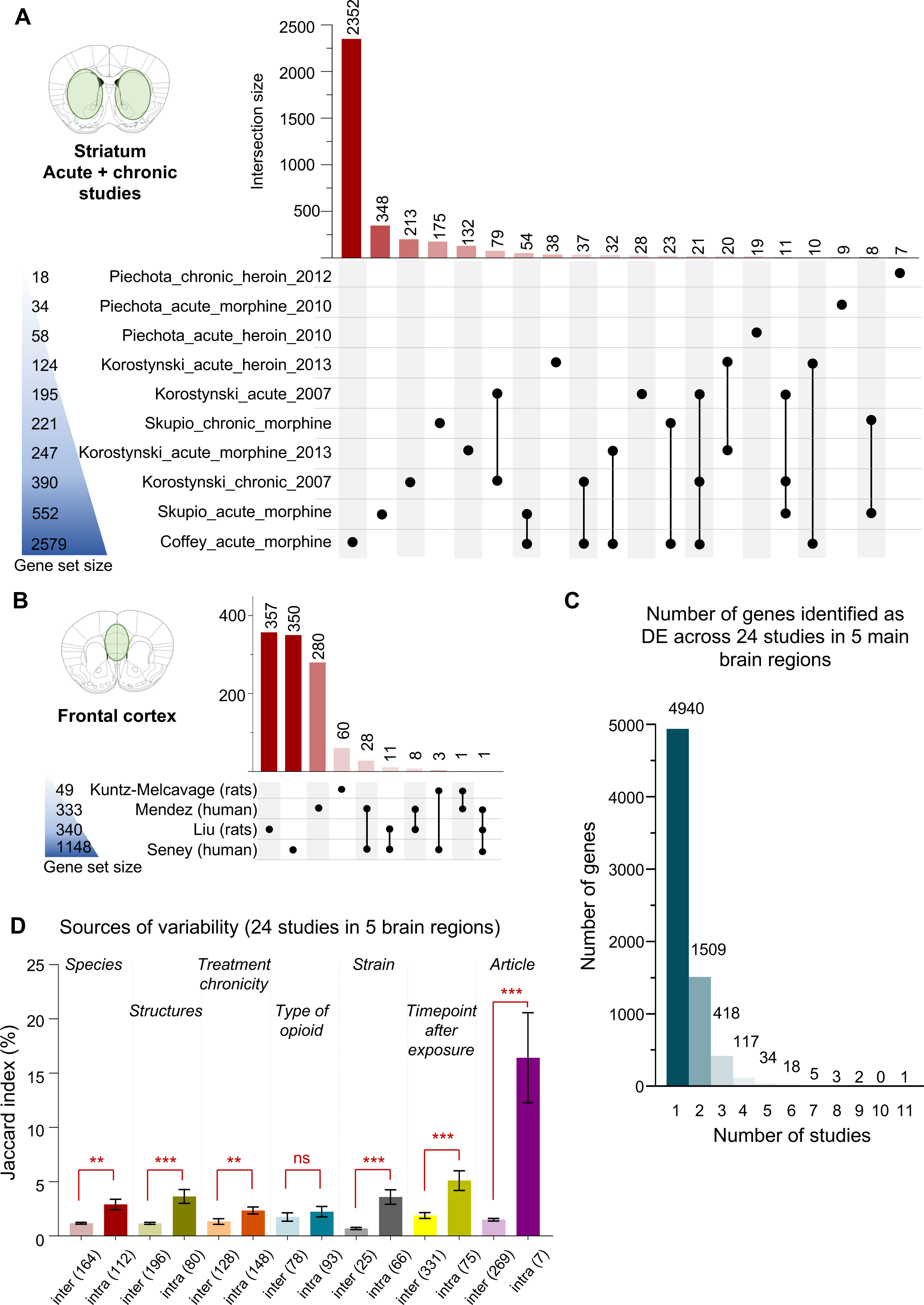
Quantitative analyses in the dorsal striatum, whole striatum and frontal cortex. **(A)** Number of differentially expressed genes (DEG) identified in studies that investigated the striatum (see Fig.S1 for separate analyses of studies that used acute or chronic opioid administration). **(B)** Number of DEG in studies of the frontal cortex. **(C)** Number of genes identified as differentially expressed (DE) across different studies. **(D)** Across the 5 brain regions considered during quantitative analysis, we assessed the impact of the following 7 factors on consistency among studies, using pairwise Jaccard indexes: species, brain structure, treatment chronicity (acute/chronic), opioid drug (morphine/heroin), strain (C57BL6J/others), time of analysis following last exposure (5 lists of DEG from tissue collected 1 hour after opioid exposure, 5 after 2 hours, 10 after 4 hours, 4 after 8 hours, 1 after 14 hours, and 3 after 24 hours) and article. For each factor (e.g., species), pair-wise JI were classified in 2 categories describing whether they belonged to the same sub-group (e.g., 2 studies conducted in humans; “intra”), or not (e.g., 1 study in humans compared to another in rats; “inter”), within that factor. For the drug factor, studies in human were not considered as they cannot be assigned to specific opioids; a single study investigated fentanyl, and was excluded[33]. For the strain factor, among the 14 mouse studies available, JI significantly increased among the 12 that used C57BL6J mice, while a similar analysis was not possible for 129P3/J, DBA/2J, SWR/J or Kunming strains, as only 1 list of DEG was available for each. One study reporting on the combined analysis of C57BL6J and DBA2J mice was excluded[54]. No similar analysis was possible in rats, as all available studies used Sprague-Dawley (except one that did not state the strain used[68]). Note that the numbers of JI values considered in each comparison are indicated at the bottom of each bar.

#### 2.4 Frontal cortex

Seven studies provided DEG (Fig.5B): 2 in humans[34, 70], 5 in rats[29, 32, 67–69]. Data from Odegaard et al[29] were discarded, as they corresponded to transgenerational effects in animals not directly exposed to opioids. For 2 additional studies [67, 69], we could not map microarray probes to genes. Among the 4 remaining articles, little overlaps were again found (Kuntz-Melcavage/Mendez, JI=0.5%; Kuntz-Melcavage/Seney, 1.1%; Liu/Mendez, 1.4%; Liu/Seney, 1.6%), with no improvement within species (Liu/Kuntz-Melcavage in rats, JI=0%; Seney/Mendez in humans, 4.1%).

#### 2.5 Spinal cord

Among the 6 spinal cord studies, 5 did not provide DEG[91–95], while the 6^th^ one used a homemade microarray[77], for which correspondence between probe IDs and genes was only partially provided. As such, no comparison was possible.

#### 2.6 Sources of variability

Considering that pharmacological recruitment and intra-cellular signaling of MOR may trigger a set of common transcriptional adaptations across the 5 regions of interest, we next pooled all lists of DEGs (Fig.5D). Again, the vast majority of DEG (70.1%, 4940/7047) were unique to a single study, while only 21.4% common to 2 studies (1509), 5.9% to 3 (418), and 1.7% to 4 or more (117; Supplementary Table 3). Interestingly, Cdkn1a (Cyclin Dependent Kinase Inhibitor 1A) was identified by 11 studies. This gene encodes for p21, a protein involved in the regulation of the cell-cycle and oligodendrocytes[127]. Avey et al[51] found Cdkn1a to be one of the most significantly upregulated genes in response to acute morphine. This upregulation occurred in oligodendrocytes, and was blocked by the opioid antagonist naltrexone, indicating a MOR-dependent mechanism. Of note, cocaine, another major drug of abuse, similarly recruits p21, as cocaine-induced behavioral responses were modified in p21 knockout mice[128]. Using this knock-out line, or viral approaches for p21 manipulation, represent appealing perspectives to further investigate this gene in OUD.

We also performed Gene Ontology (GO) analysis using DEG common to at least 3, 4 or 5 studies (Supplementary Table 4). No results were significant for the small list of 63 DEG common to 5 or more studies. When considering the 180 DEGs common to at least 4 studies, only 3 GO terms were enriched: cell periphery, plasma membrane and hormone activity. Results more directly related to the nervous system emerged for the 598 DEG common to at least 3 studies, with 46 significant terms partly related to synaptic signaling and cell-cell communication, which may reflect general opioid-induced electrophysiological and synaptic adaptations across brain structures.

Because weak DEG overlaps were identified within individual regions (even when grouping studies based on species, Fig.4C-E, or treatment chronicity, Fig.S1), we sought to more systematically assess, across the 5 regions, the impact of following factors: species, brain structure, treatment chronicity, drug, strain, time-point of assessment following last exposure, and article (Fig.5D). For each factor (e.g., species), pair-wise JI were classified in 2 categories depending on whether they belonged to the same sub-group within that factor (2 studies in humans; “intra”), or not (1 human study compared to another in rats; “inter”). As expected, grouping list of DEG by species (p=2.5E-3), brain structures (3.4E-8), treatment chronicity (p=1.4E-3), mouse genetic background (3.9E-5) or the time-point of analysis after opioid exposure (6.7E-10) all increased concordance, with higher JI. This is consistent with the notion that, although they remain poorly characterized, these 5 variables contribute to opioid-induced transcriptional plasticity. In contrast, the type of opioid used had no significant impact (p=0.059) when considering the 13 and 5 rodent DEG lists related to morphine or heroin, respectively (human studies were discarded, as the types of opioids consumed are diverse and poorly characterized). This is surprising, considering that these 2 MOR agonists exhibit strong pharmacokinetic differences and would be expected to generate distinct adaptations (heroin being a lipophilic prodrug that rapidly crosses the blood brain barrier to be metabolized in 6-monoacetylmorphine and morphine, which in turn activate MOR). Time-course and dose-response experiments may be necessary to detect such differences. Importantly, concordance increased most when DEG lists came from the same publication (p=2.5E-5), despite the fact that only 7 comparisons fell under this category. As already mentioned, this suggests that differences across laboratories may explain a substantial part of the lack of concordance across studies.

Finally, we refined our analysis by considering the directionality (up/down) of DEG, focusing on genes identified by at least 5 studies (n=63). Among results from 17 out of 24 studies that reported on directionality (Fig.S2), most genes (37/63, 59%) showed discordant results (while, for 8 others, directionality was reported by only 1 or no study). Only 18 genes showed 100% concordance across studies (29%; 17 up-regulated, 1 down), among which 11 were identified by at least 3 studies. The latter included the aforementioned Cdkn1a, as well as Slc2a1, Fkbp5, Sult1a1, Arrdc3, Ccdc117, Plin4, Wscd1, Arid5b, Pla2g3, and Tsc22d3.

Slc2a1 encodes for the glutamate transporter GLT1, which plays an important role in extracellular glutamate uptake and the regulation of excitatory transmission. A recent study showed that during prolonged morphine withdrawal, activation of KOR in the amygdala leads to increased GLT1 expression and excitatory drive in the NAc, thereby mediating withdrawal-induced depressive-like behaviors. Another interesting hit is Fkbp5, a chaperone and intra-cellular negative regulator of glucocorticoid signaling that has been largely investigated in stress-related disorders[129], including OUD[130]. The 8 remaining genes, although comparatively less studied in relation to opioids (Fig.S2), represent candidates for future work.

#### 2.7 Threshold-free genome-wide comparison

Next, we compared the 2 NAc studies for which full genome-wide results were available (Seney et al[34], Townsend et al[33]). While both included males and females, only Townsend reported sex-specific results. Therefore, we reprocessed Seney’s raw data to identify DEG in each sex separately (*Supplementary Material*). This slightly increased concordance, arguing for the importance of accounting for sex: while JI were equal to 7.5 and 4% when comparing initial Seney data (male/females pooled) with Townsend data in females or males, respectively (Fig.S3A-B), they increased to 9.5% and 4.5% when considering Seney’s sex-specific results (Fig.6A-B).

**Figure 6.**
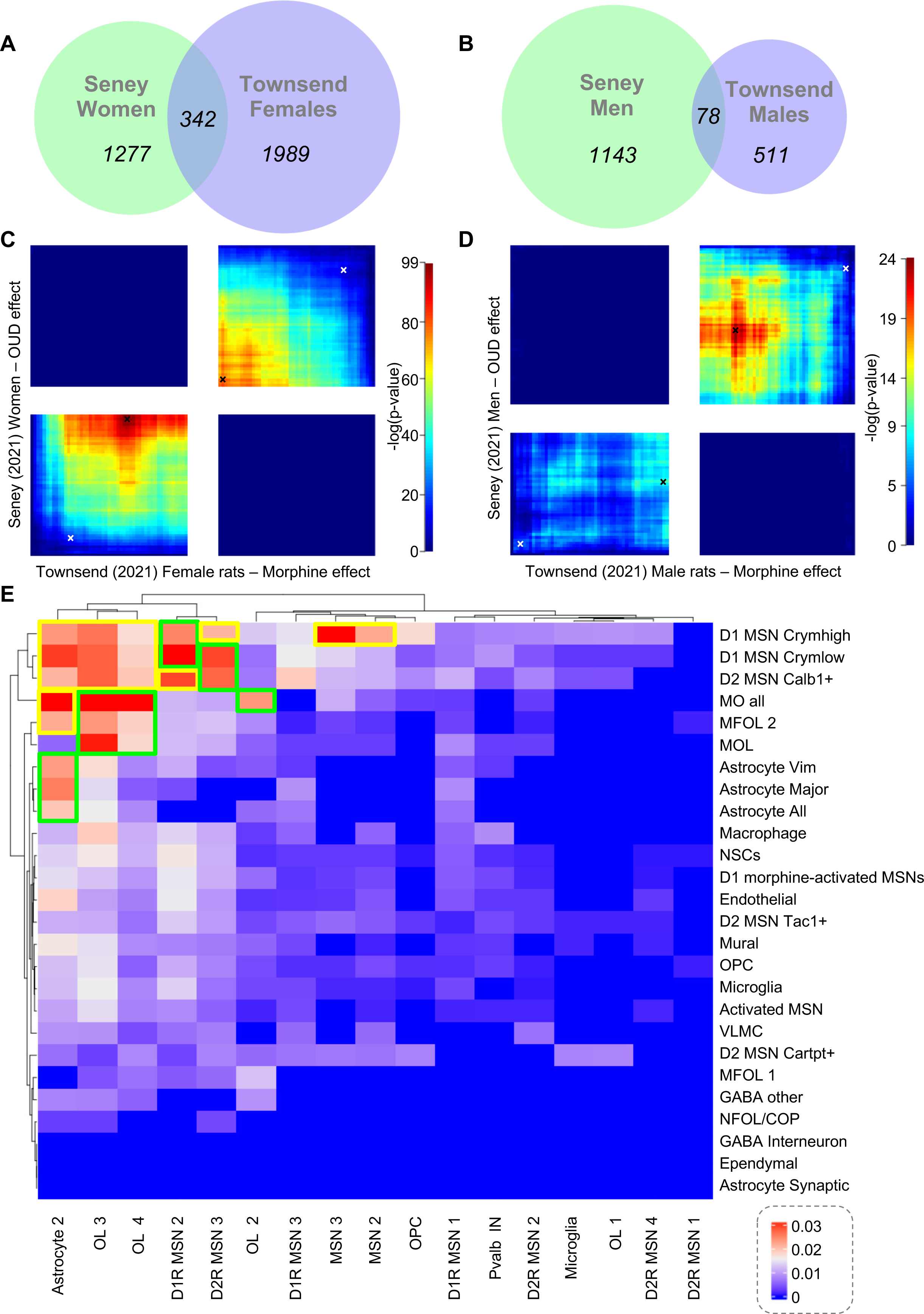
(A-D) Threshold-free comparison of 2 bulk tissue RNA-sequencing studies (Seney et al and Townsend et al), and cell-type specific comparison of 2 single-cell (Avey et al) and single-nucleus (Reiner et al) RNA-sequencing studies. **(A-B)** Venn diagram of differentially expressed genes (DEG) identified across the 2 bulk tissue RNA-Sequencing studies by Seney et al and Townsend et al (DEG defined by arbitrary significance thresholds in each study: p-value < 0.05 for both) for women/female rats (A) and men/male rats (B). **(C-D)** Genome-wide, threshold-free comparison of the same 2 studies using Rank-Rank Hypergeometric Overlap (RRHO2) for women/female (C) rats and men/male rats (D). In RRHO2, briefly, genes are ranked using p-values (signed according to directionality), from upregulated (Seney: left, Townsend: bottom) to down-regulated genes (Seney: right, fentan: top). Overlaps along the 2 distributions are then iteratively assessed using hypergeometric testing. High p-values in the bottom-left and top-right quadrants indicates concordance, whereas high p-values in the other 2 quadrants (bottom-right and top-left) indicate discordance between the 2 datasets. The white crosses indicate the location of the overlap corresponding to Venn diagrams depicted in panels A-B (split in subgroups of up- or down-regulated genes), while black crosses correspond to most significant overlaps identified using RRHO2. **(E)** Heatmap of pairwise Jaccard indexes for DEG identified across each of the 26 and 27 cell-types defined by Avey et al and Reiner et al, respectively.

We then used RRHO2 (Rank–Rank Hypergeometric Overlap[131]) to compare the 2 studies. RRHO2 iteratively performs hypergeometric testing for all combinations of thresholds applied to each dataset, generating a “threshold-free” analysis. Because this approach takes directionality into account, we also computed, for comparison, overlaps among either up- or down-regulated DEG reported by each study (white crosses, Fig.6C-D). Compared to the latter, much more significant overlaps were identified when RRHO2 considered less stringent p-value thresholds (best hypergeometric results are shown as black crosses). By definition, these increased overlaps corresponded to large gene lists which, although commonly dysregulated in similar direction, individually exhibited milder FC. Interestingly, these overlaps were detected despite the fact that Seney investigated human post mortem tissue from individuals with OUD, while Townsend focused on rat tissue collected after 24 hours of withdrawal from fentanyl IVSA. Also, different patterns of convergence were observed in each sex: in females, overlaps were highly significant among both up- and down-regulated genes while, in males, a lower concordance predominantly affected down-regulated genes. This suggests that the rat dataset may differentially capture sex-specific signatures of the human disorder. Overall, these analyses illustrate the utility of reporting full genome-wide data which, in combination with threshold-free approaches, may generate deeper understanding (see *Discussion*).

### 3. High cellular resolution approaches

Studies reviewed above indicate that opioids induce genome-wide transcriptomic changes, with striking variability across laboratories, experimental designs, and brain regions. This likely reflects a combination of: i) direct pharmacological opioid effects, which correspond to the recruitment of MOR intracellular signaling and result in decreased neuronal excitability[132]; ii) indirect effects, as the chemical identity and connectivity pattern of MOR-expressing cells determine how their inhibition leads to secondary changes affecting other neurotransmitters[117]. This raises 2 questions: how is each cell type, or even each brain cell, affected by these changes? What are underlying molecular regulatory principles?

To address the first question, several groups recently used cell-type specific and single-cell methodologies. Coffey et al[31] focused on microglial cells of the whole striatum, using translational ribosome affinity purification (TRAP, to capture ribosomes and the “translating transcriptome”[133, 134]). Chronic morphine effects were analyzed with, or without, the induction of naloxone-precipitated withdrawal. Results showed that both chronic morphine and withdrawal triggered numerous changes in gene expression, which mostly affected GO terms related to cAMP signaling, regulation of synapses, and the cytoskeleton. A striking negative correlation was observed between the 2 sets of results, providing a correlate, at the level of the microglial transcriptome, of the pharmacological antagonism between morphine and naloxone. Considering the inter-strain variability in the sensitivity to both acute and chronic opioid physical dependence described earlier (see section *1. Qualitative analysis-Route and duration of opioid administration* and[39, 40]), an interesting perspective would be to determine whether naloxone similarly antagonizes transcriptomic effects of acute opioid exposure, and how this is modulated by underlying genetic variation. Of note, surprisingly, such cell-type specific approaches have not been used to interrogate opioid effects in neuronal cells, let alone opioid-responsive MOR-expressing neurons which, arguably, may be particularly responsive. This is feasible, as knock-in lines in which the Cre recombinase is selectively active in MOR-expressing cells are now available[135–137].

Achieving higher resolution, Avey and colleagues applied single-cell RNA-seq (scRNA-Seq, using Drop-seq[138]) to analyze transcriptomic changes induced by acute morphine in more than 23,000 cells of the mouse NAc[51]. As is typical in scRNA-Seq, sequencing depth yielded data for ∼1,600 genes, corresponding to a reduced genomic representation. Changes were widespread and, interestingly, most frequent in oligodendrocytes, followed by neurons, astrocytes, and microglia, therefore uncovering a cellular hierarchy in transcriptional sensitivity to opioids. More recently, Reiner et al[41] used single-nucleus RNA-Seq (snRNA-seq, 10x Genomics 3’) to analyze a larger number of nuclei (190,030) and genes (>2000) in the rat NAc, also following passive and acute morphine injection. Compared to the former study, DEG mostly affected neurons and oligodendrocytes, followed by astrocytes and microglia. Morphine IVSA during 10 days was also investigated, and compared to acute effects. Among DEG shared across the 2 conditions (n=431), the vast majority were dysregulated in a similar direction (up/down), suggesting that they may reflect morphine exposure rather than volitional drug-taking (a “yoked” group chronically exposed to morphine would help substantiate this distinction). Importantly, a larger number of DEG were specifically dysregulated in only one condition (n=868), with more DEG associated with IVSA (reminiscent of the study by Tapocik et al described above), and distinct distributions among NAc cell-types (e.g., Dopamine receptor D1- or D2-expressing neurons). By providing the first single-cell description of morphine IVSA, these results reinforce the notion that voluntary drug-intake may recruit partly distinct molecular mechanisms from those of non-contingent administration.

We next compared results on acute morphine from these 2 studies. Both used Seurat[139] and dimensionality reduction (t-SNE[51] or UMAP[41]) to identify clusters, which were then assigned to cell-types using external data. We first compared marker genes reported to define those clusters, when available (Avey: 11/29 clusters; Reiner: 11/19 clusters). Low agreement was found (Fig.S4A), with JI ranging from 0 up to 7.1% (for Avey’s activated medium spiny neurons, MSNs, and Reiner’s D1R MSN-1), possibly because both studies used few marker genes (Avey: 39-51; Reiner: 4-12). We then confronted morphine-induced DEG for each cell-type (Fig.6E): low JI were found, with maximal overlap (6.4%) obtained when comparing Avey’s “myelin-forming and mature oligodendrocytes” (MO_all) with Reiner’s “Oligodendrocyte-3”. Surprisingly, while some clusters assigned to similar cell-types in the 2 studies showed some similarity in DEG (green squares, Fig.6E), strong concordance was also observed across clusters with different identities (e.g. Reiner’s “Astrocyte-2” and Avey’s MO_all; yellow squares). Overlaps among DEG (Fig.6E) did not increase as a function of cell-type similarity across studies (Fig.S4B), as would have been expected. While a particular emphasis has been put in sc/snRNA-Seq studies on the number of individual cells to be sequenced, these results suggest that in order to maximize their utility: i) marker genes used to assign cellular identities to clusters should be reported comprehensively; and ii) rather than short lists of DEG, differential expression results should be provided for all genes analyzed in each cluster. In addition, classical issues related to sample size, power or statistical models for group comparisons remain pregnant. The Avey and Reiner studies used similar sample sizes (4 and 5 replicates per opioid or control group, respectively), in the upper range of bulk tissue studies (Fig.2D), but applied distinct tools to identify DEG (the SCDE and MAST packages, respectively), which may partly account for discrepancies. With the rapid evolution of this field and emerging controversies (e.g. regarding dimensionality reduction[140]), additional efforts will be necessary to reach consensus on designs and analyses[141, 142].

### 4. Non-coding and epigenomic mechanisms

In parallel to efforts to deepen cellular resolution, others have sought to decipher underlying non-coding RNA and epigenomic regulations[143]. Although neurons are post-mitotic cells, they exhibit peculiar plasticity features that may contribute to the encoding of life events, including exposure to psychoactive drugs[144]. Several non-coding RNA species have been investigated. The first report used custom-made arrays to examine micro-RNA (miR) following morphine or heroin exposure, in rat primary neuronal cultures or the mouse hippocampus[99]. 7 miR exhibited significant expression changes, among which miR-339-3p was later shown to target the 3’-UTR and down-regulate MOR in response to opioid treatment[145]. A second study used more recent arrays to interrogate all mouse miR in the NAc, following chronic heroin injections[63]. 111 miR exhibited >25% expression changes (no significance reported). The study focused on miR-218, and showed that its over-expression inhibited heroin-induced reinforcement (in conditioned place preference and IVSA), possibly by targeting MeCP2 (a protein that binds methylated DNA), suggesting a potential link between opioids, miR and DNA methylation. Other studies explored long non-coding (lncRNA) or circular RNA (circRNA). One group used 2 arrays interrogating 9,000 lncRNA[93] and 15,000 circRNA[96] to characterize chronic morphine effects in the rat spinal cord: 136 lncRNA (84 up, 52 down; nominal p-value<0.05, |log2FC|>1.5) and 2038 circRNA (896 up, 1142 down; nominal p-value<0.05, FC>2) were affected. Another report used RNA-Sequencing to focus on circRNA in the NAc[64], among which 112 were differentially expressed between mice that had received chronic morphine injections with or without electroacupuncture (51 up, 61 down; nominal p-value<0.05, |log2FC|>1.5). Understanding how these non-coding RNA contribute to OUD will require further work, as numerous mechanisms have been implicated in their physiological effects (including the sequestering of miR by circRNA, or complementarity interactions among lncRNA and mRNA).

Regarding the epigenome, a handful of candidate studies had initially provided evidence for differential DNA methylation at specific loci: the MOR gene in human blood[146–148], and a few loci in the rat brain[149]. At the genome-wide scale, 6 recent studies (1 in rats[62], 5 in humans investigated DNA methylation. The rat study used Whole Genome Bisulfite Sequencing (WGBS) to analyze DNA methylation changes after heroin IVSA, in a standard or enriched environment, in 3 brain regions (VTA, NAc and medial prefrontal cortex)[62]. Because of low sequencing depth, DNA methylation was averaged across the whole genome for selected features (promoters, CG islands). As expected, neither IVSA nor environmental enrichment had any impact on these metrics, suggesting that the methylome is not massively reprogrammed. Opioids may still, however, trigger more discrete DNA methylation changes, consistent with the intuition that the large transcriptomic changes associated with opioid exposure reviewed above are unlikely to occur in the absence of any methylomic plasticity.

Two human studies conducted on bulk brain tissue strengthen this hypothesis. Focusing on Brodmann area 9 (BA9), the first applied EPIC arrays to a cohort of OUD patients (n=19) and controls (n=11)[75]. Among >800,000 CG sites (where DNA methylation mostly occurs in the mammalian brain), 11,917 located at transcription start sites (TSS) showed nominally significant differences (p-value<0.05; unfortunately, genomic localization of these probes, and full genome-wide results, were not provided). Such studies face the difficult task of controlling for confounding factors, as variation in genotype, age, sex, tissue cellular composition or socio-demographic factors have all been associated with DNA methylation changes. The second study addressed these questions[76]. Adjusting for socio-demographic characteristics, ancestry and cellular composition, the authors investigated dorsolateral prefrontal cortex from a large cohort (n=153) of 72 individuals who died of acute opioid intoxication, 53 psychiatric controls, and 28 healthy controls. Although no individual CG site survived multiple testing correction, 13 passed a relaxed significance threshold (p<1.0E-05) for an association with opioid intoxication, one of which was located in Netrin-1, a gene regulating KOR synthesis. At the genome-wide level, opioid use was also associated with enriched differential methylation in the KEGG pathway related to dopaminergic signaling.

While aforementioned studies focused on bulk tissue, 3 others used Fluorescence-Activated Nuclei Sorting (FANS) to achieve cell-type specific analysis of the neuronal methylome, in the orbitofrontal cortex[72–74]. The first used 450K arrays and, comparing 37 heroin users who died from heroin overdose with 28 controls, identified 1298 differentially methylated CG sites, which were more frequently annotated to genes expressed by glutamatergic than GABAergic neurons[73]. The 2 last studies focused on a smaller cohort of male subjects only (12 OUD patients, 26 controls), using Reduced Representation oxidative Bisulfite Sequencing (RRoxBS). Compared to arrays, this allowed for the investigation of a broader set of cytosines (3.5 million CG sites at an average 10X coverage) and, importantly, the characterization of 2 types of non-canonical modifications enriched in the brain: DNA methylation in the non-CG context (also called mCH, where H stands for A, C or T)[150, 151] and hydroxymethylation at CG sites (or hmCG, generated by oxidation of methylated CG, mCG, by *TET* enzymes[152]). Differential methylation events were found to be more frequent in the non-CG (n=2352) than in the reference CG context (n=397), with enrichments in GO terms and gene sets (from Genome-Wide Association Studies, GWAS) related to neuronal function and psychiatric diseases[72]. Similarly, hmCG levels were found more frequently modified (n=1740) than mCG, at least in genomic regions covered by RRoxBS, and enriched in GO terms and co-methylation modules relevant to neuronal physiology[74]. Together, these 2 studies suggest that OUD pathophysiology may be strongly dependent on atypical, brain-enriched forms of methylomic plasticity. Of note, these human studies identified OUD-associated differences at the level of individual cytosines. However, evidence from other research fields and animal models[153, 154] argues for a model whereby DNA methylation changes with functional impact (e.g. on transcription factor binding, or interactions with other epigenetic factors) implicate groups of cytosines in close proximity and with differential methylation in similar direction (differentially methylated regions, DMR), rather than individual sites. Future work should therefore seek to combine DMR calling with the statistical models necessary to account for human confounding factors.

Interacting with these DNA modifications, epigenomic plasticity also relies on histone proteins and chromatin structure or accessibility. Surprisingly, a single study characterized histone modifications in the mouse NAc[65]. This pioneering work showed that behavioral effects of chronic morphine (including locomotor sensitization and precipitated withdrawal) were bidirectionally modulated by down- or over-expression of the histone methyltransferase G9a, and accompanied by modulation of H3K9me2 levels across thousands of genomic sites. While these results, and later work on cocaine[152, 155], suggest widespread histone changes, no follow-up work has been published in relation to opioids, to our knowledge. Similarly, only one study has investigated chromatin accessibility. Focusing on a human cohort of 10 heroin users who died from heroin overdose, and 10 healthy individuals[87], genomic regions with “open” chromatin were identified using ATAC-Seq, in the NAc and putamen. The technique was applied to neuronal and non-neuronal cells isolated by FANS. Heroin use was associated with enriched chromatin accessibility in specific gene (CG islands, promoters, 5’ untranslated regions) or regulatory (peaks for the H3K4me1 histone mark, binding sites for the EZH2 transcription factor) features, as well as more frequent changes in neurons than in non-neurons (affecting GO terms related to the synapse, among others). At the behavioral level, FYN, a tyrosine kinase prioritized from human data, was then shown in the rat to significantly contribute to heroin IVSA. Overall, this work nicely illustrates how epigenetic analyses may uncover chromatin mechanisms contributing to OUD.

## Discussion

Here we systematically assessed functional genomic mechanisms of opioid action and OUD, and uncovered a surprisingly low convergence across studies. As a result, a lot remains to be done to move the field forward. Below, we propose recommendations for study design and data sharing, as well as perspectives to refine behavioral modelling, improve cellular resolution, and advance multiomic data integration.

### Recommendations for study design and data sharing

The low sample sizes observed in the present review (Fig.2D,F) were surprising, considering that this factor is arguably the most important one influencing statistical power[156–158]. Accordingly, a more rigorous analysis of power, and larger sample sizes[159], will lead in future work to better reproducibility, and should be enforced, especially when studying rodent outbred strains, or humans, who display higher inter-individual variability[160, 161]. In humans, similar to what has been observed for GWAS[162], functional genomic studies of OUD are expected to evolve in the next decade towards collaborative efforts involving multiple labs (e.g. the SCORCH consortium[163]), and the progressive aggregation of ever larger cohorts. Another major recommendation relates to data sharing. While public repositories have been specifically implemented for collecting and disseminating omic data (GEO[164], the IHEC data portal[165]) or bioinformatic pipelines (Zenodo, GitHub), their systematic use is not yet current practice in the OUD field, as exemplified in the present work (with similar criticisms raised in other research domains[166]). This hampers re-analysis of published data, comparisons of results across laboratories, and the use of threshold-free methods. We believe this last point is particularly meaningful. Most opioid studies report subsets of DEG identified using arbitrary significance cut-offs, which can mask subtle changes holding relevant biological information (Fig.6[131]). We therefore strongly advocate for generalized use of threshold-free algorithms for enrichment analysis of single datasets (GSEA), or comparison across pairs of differential analysis results (RRHO2). Within this line, newer packages are expected to scale-up the applicability of such tools to epigenomic data, which frequently involve larger number of observations than gene-level counts[167].

### Refine behavioral modeling

The face validity of SA or oICSS models of OUD has significantly increased over last 2 decades[168], and we now face the task of more systematically combining them with most recent molecular approaches. Another important step will be to decipher the molecular mechanisms that may differentially account for distinct behavioral dimensions, among impulsive or persistent drug-seeking, resistance to punishment, or excessive motivation (including those modelled using passive administration, such as tolerance, sensitization, or withdrawal). While the vast majority of studies reviewed here used a categorical comparison of animals exposed or not to opioids, another design, based on a dimensional approach within the exposed group, could help address the inter-individual variability in susceptibility or resilience to OUD. The latter design has been used by only one article in the present review[56] (section *2.1*), which contrasts with the amount of data available for models of other psychiatric disorders, in particular depression (e.g. chronic social defeat[169–171]. To better understand such inter-individual susceptibility, epigenetic mechanisms are obvious candidates[172], as illustrated by 2 recent studies on DNA methylation and oral SA of alcohol in macaques[173], and miRNA in a rat model of cocaine SA[174]. While similar mechanisms are likely at play in OUD, they have not been addressed by open-ended approaches yet, pointing toward an important avenue for future work.

### Improve cellular resolution

This is another major avenue for future work, considering the extreme heterogeneity of brain tissue. In addition to cell-type specific (TRAP) and sc/snRNA-Seq (already applied to OUD, see above), a newer generation of tools now enables similar molecular profiling while taking neuroanatomical context into consideration, with 2 possible factors: i) connectivity patterns, probed using antero- or retrograde viruses, in combination with TRAP (transcriptomics[175]) or FANS (epigenomics[176]); or ii) spatial context, with methodologies based on sequencing or microscopy to detect RNA “in histological space”; while spatial capture followed by transcriptome sequencing (e.g. Slide-seq[177]) enables genome-wide analysis with 10µm resolution, single-molecule fluorescence in situ hybridization (smFISH) allows for better spatial resolution (1µm), but requires probe design and target gene selection[146,147] (covering a few hundred to a few thousand genes, lower than sc/snRNA-Seq[180]). Future work using these techniques, by characterizing molecular effects of opioids in a particular neuronal pathway or cell-type, or with exquisite spatial resolution, should deepen the understanding of OUD.

### Advance multiomic data integration

The epigenomic and transcriptomic mechanisms reviewed above contribute to changes at protein level that ultimately mediate neurophysiological adaptations. Proteomic studies were initially not included in our systematic bibliography, considering the low number of studies available and because, compared to RNA- or DNA-sequencing methodologies, they only cover a few hundred proteins[181–184]). Nevertheless, to go further in the understanding of OUD, future work will need to combine transcriptomic and proteomic analyses with the characterization of upstream epigenetic regulation. The next challenge, therefore, will be to integrate those various molecular layers into multiomic models of pathophysiology.

Illustrating these concepts, we recently conducted a human study suggesting that early-life adversity and depression associate with 2 forms of DNA methylation changes in the CG and non-CG contexts (specifically focusing on CAC sites, where non-CG methylation is most abundant)[150]. By combining WGBS with ChIP-Seq analysis of 6 histone marks, the 2 forms of plasticity were found to affect distinct genomic sites and features, and to be defined by different histones modifications and chromatin states. Similar multiomic investigations of opioid effects should be conducted and, importantly, will require dedicated bioinformatic frameworks, which are only emerging in psychiatry[185]. Only 2 studies reviewed here followed that path: Mendez et al performed transcriptomic and proteomic analyses, while Liu et al analyzed the transcriptome and methylome (see *4.Non-coding and epigenomic mechanisms of opioid plasticity*). In both studies, multiomic integration consisted: first, in the identification of features (single mRNA, DNA methylation site, or protein) passing a significance threshold, and their overlap; second, in the construction of co-expression or co-methylation networks, and the prioritization of groups of features, defined as modules, most significantly associated with OUD, and their biological interpretation (enrichment in functional GO terms, significantly affected features, GWAS results, etc). While such systems-biology approaches will undoubtedly contribute to the understanding of OUD, they remain based on step-by-step aggregation of results generated individually for each layer. In the coming years, it is expected that advanced multiomic integration approaches (e.g. similarity network fusion, or multiblock methods[186–188]), may more globally define, throughout the genome, how multiple regulatory layers interact, and the chain of events linking genetic, developmental, and environmental factors to abnormal behavior and compulsive drug use[189].

## Conclusion

Although numerous functional genomic studies of opioid action and OUD have been conducted across animal models and patient cohorts, replicable findings appear limited. In the future, combining refined behavioral models with multiomic integration, at the single-cell scale, will hopefully pave the way towards the identification of innovative therapeutic targets and, ultimately, contribute to better care.

## Acknowledgements

This research was supported by the Centre National de la Recherche Scientifique (CNRS, France), ‘Université de Strasbourg’ (Idex Recherche Exploratoire 2022; PEL), French National Research Agency (ANR-19-CE37-0010; PEL), ‘Fondation Fyssen’ (Subvention recherche 2021), ‘Fondation Avenir’ (AAP 2022 Recherche médicale Appliquée), ‘Fondation pour la Recherche Médicale’ (FDT202204015236; CF), and ‘Institut pour la Recherche en Santé Publique – Institut National du Cancer’ (IReSP-INCa doctoral fellowship 2022-166; ACR).

## Code availability

All code used in the present review is available as Supplementary File 1.

## References

1. Rosenblum A, Marsch LA, Joseph H, Portenoy RK. Opioids and the treatment of chronic pain: controversies, current status, and future directions. Exp Clin Psychopharmacol. 2008;16:405– 416.

2. Florence C, Luo F, Rice K. The economic burden of opioid use disorder and fatal opioid overdose in the United States, 2017. Drug Alcohol Depend. 2021;218:108350.

3. European Monitoring Centre for Drugs and Drug Addiction. European drug report 2017:trends and developments. LU: Publications Office; 2017.

4. Heilig M, MacKillop J, Martinez D, Rehm J, Leggio L, Vanderschuren LJMJ. Addiction as a brain disease revised: why it still matters, and the need for consilience. Neuropsychopharmacology. 2021;46:1715–1723.

5. Maze I, Shen L, Zhang B, Garcia BA, Shao N, Mitchell A, et al. Analytical tools and current challenges in the modern era of neuroepigenomics. Nat Neurosci. 2014;17:1476–1490.

6. Przewlocki R. Opioid abuse and brain gene expression. Eur J Pharmacol. 2004;500:331–349.

7. McClung CA. The molecular mechanisms of morphine addiction. Rev Neurosci. 2006;17:393– 402.

8. Koob GF, Volkow ND. Neurobiology of addiction: a neurocircuitry analysis. Lancet Psychiatry. 2016;3:760–773.

9. Reed B, Kreek MJ. Genetic Vulnerability to Opioid Addiction. Cold Spring Harb Perspect Med. 2021;11:a039735.

10. Bodnar RJ. Endogenous opiates and behavior: 2020. Peptides. 2022;151:170752.

11. Przewlocki R. Opioid abuse and brain gene expression. European Journal of Pharmacology. 2004;500:331–349.

12. McClung CA. The molecular mechanisms of morphine addiction. Reviews in the Neurosciences. 2006;17:393–402.

13. Yamada K, Nagai T, Nabeshima T. Drug dependence, synaptic plasticity, and tissue plasminogen activator. J Pharmacol Sci. 2005;97:157–161.

14. UNODC, World Drug Report 2022 (United Nations publication, 2022).

15. Kerridge BT, Saha TD, Chou SP, Zhang H, Jung J, Ruan WJ, et al. Gender and nonmedical prescription opioid use and DSM-5 nonmedical prescription opioid use disorder: Results from the National Epidemiologic Survey on Alcohol and Related Conditions - III. Drug Alcohol Depend. 2015;156:47–56.

16. Bravo IM, Luster BR, Flanigan ME, Perez PJ, Cogan ES, Schmidt KT, et al. Divergent behavioral responses in protracted opioid withdrawal in male and female C57BL/6J mice. Eur J Neurosci. 2020;51:742–754.

17. Craft RM. Sex differences in opioid analgesia: ‘from mouse to man’. Clin J Pain. 2003;19:175– 186.

18. Craft RM, Stratmann JA, Bartok RE, Walpole TI, King SJ. Sex differences in development of morphine tolerance and dependence in the rat. Psychopharmacology (Berl). 1999;143:1–7.

19. Cicero TJ, Nock B, Meyer ER. Gender-linked differences in the expression of physical dependence in the rat. Pharmacol Biochem Behav. 2002;72:691–697.

20. Karami M, Zarrindast MR. Morphine sex-dependently induced place conditioning in adult Wistar rats. Eur J Pharmacol. 2008;582:78–87.

21. Cicero TJ, Aylward SC, Meyer ER. Gender differences in the intravenous self-administration of mu opiate agonists. Pharmacol Biochem Behav. 2003;74:541–549.

22. Beery AK, Zucker I. Sex bias in neuroscience and biomedical research. Neurosci Biobehav Rev. 2011;35:565–572.

23. Prendergast BJ, Onishi KG, Zucker I. Female mice liberated for inclusion in neuroscience and biomedical research. Neurosci Biobehav Rev. 2014;40:1–5.

24. Labonté B, Engmann O, Purushothaman I, Menard C, Wang J, Tan C, et al. Sex-specific transcriptional signatures in human depression. Nat Med. 2017;23:1102–1111.

25. Hoffman GE, Ma Y, Montgomery KS, Bendl J, Jaiswal MK, Kozlenkov A, et al. Sex Differences in the Human Brain Transcriptome of Cases With Schizophrenia. Biol Psychiatry. 2022;91:92–101.

26. Hitzemann R, Bergeson SE, Berman AE, Bubier JA, Chesler EJ, Finn DA, et al. Sex Differences in the Brain Transcriptome Related to Alcohol Effects and Alcohol Use Disorder. Biol Psychiatry. 2022;91:43–52.

27. Vassoler FM, Oliver DJ, Wyse C, Blau A, Shtutman M, Turner JR, et al. Transgenerational attenuation of opioid self-administration as a consequence of adolescent morphine exposure. Neuropharmacology. 2017;113:271–280.

28. Borrelli KN, Yao EJ, Yen WW, Phadke RA, Ruan QT, Chen MM, et al. Sex Differences in Behavioral and Brainstem Transcriptomic Neuroadaptations following Neonatal Opioid Exposure in Outbred Mice. ENeuro. 2021;8:ENEURO.0143-21.2021.

29. Odegaard KE, Schaal VL, Clark AR, Koul S, Sankarasubramanian J, Xia Z, et al. A Holistic Systems Approach to Characterize the Impact of Pre- and Post-natal Oxycodone Exposure on Neurodevelopment and Behavior. Front Cell Dev Biol. 2020;8:619199.

30. Odegaard KE, Schaal VL, Clark AR, Koul S, Gowen A, Sankarasubramani J, et al. Characterization of the intergenerational impact of in utero and postnatal oxycodone exposure. Transl Psychiatry. 2020;10:329.

31. Coffey KR, Lesiak AJ, Marx RG, Vo EK, Garden GA, Neumaier JF. A cAMP-Related Gene Network in Microglia Is Inversely Regulated by Morphine Tolerance and Withdrawal. Biol Psychiatry Glob Open Sci. 2022;2:180–189.

32. Liu SX, Gades MS, Swain Y, Ramakrishnan A, Harris AC, Tran PV, et al. Repeated morphine exposure activates synaptogenesis and other neuroplasticity-related gene networks in the dorsomedial prefrontal cortex of male and female rats. Drug Alcohol Depend. 2021;221:108598.

33. Townsend EA, Kim RK, Robinson HL, Marsh SA, Banks ML, Hamilton PJ. Opioid withdrawal produces sex-specific effects on fentanyl-vs.-food choice and mesolimbic transcription. Biol Psychiatry Glob Open Sci. 2021;1:112–122.

34. Seney ML, Kim S-M, Glausier JR, Hildebrand MA, Xue X, Zong W, et al. Transcriptional Alterations in Dorsolateral Prefrontal Cortex and Nucleus Accumbens Implicate Neuroinflammation and Synaptic Remodeling in Opioid Use Disorder. Biol Psychiatry. 2021;90:550–562.

35. Lefevre EM, Pisansky MT, Toddes C, Baruffaldi F, Pravetoni M, Tian L, et al. Interruption of continuous opioid exposure exacerbates drug-evoked adaptations in the mesolimbic dopamine system. Neuropsychopharmacology: Official Publication of the American College of Neuropsychopharmacology. 2020;45:1781–1792.

36. Sugino S, Konno D, Abe J, Imamura-Kawasawa Y, Kido K, Suzuki J, et al. Crucial involvement of catecholamine neurotransmission in postoperative nausea and vomiting: Whole-transcriptome profiling in the rat nucleus of the solitary tract. Genes Brain Behav. 2021:e12759.

37. Ye J, Yang Z, Li C, Cai M, Zhou D, Zhang Q, et al. NF-κB signaling and vesicle transport are correlated with the reactivation of the memory trace of morphine dependence. Diagn Pathol. 2014;9:142.

38. Korostynski M, Piechota M, Dzbek J, Mlynarski W, Szklarczyk K, Ziolkowska B, et al. Novel drug-regulated transcriptional networks in brain reveal pharmacological properties of psychotropic drugs. BMC Genomics. 2013;14:606.

39. Klein G, Juni A, Waxman AR, Arout CA, Inturrisi CE, Kest B. A survey of acute and chronic heroin dependence in ten inbred mouse strains: evidence of genetic correlation with morphine dependence. Pharmacol Biochem Behav. 2008;90:447–452.

40. Kest B, Palmese CA, Hopkins E, Adler M, Juni A, Mogil JS. Naloxone-precipitated withdrawal jumping in 11 inbred mouse strains: evidence for common genetic mechanisms in acute and chronic morphine physical dependence. Neuroscience. 2002;115:463–469.

41. Reiner BC, Zhang Y, Stein LM, Perea ED, Arauco-Shapiro G, Ben Nathan J, et al. Single nucleus transcriptomic analysis of rat nucleus accumbens reveals cell type-specific patterns of gene expression associated with volitional morphine intake. Transl Psychiatry. 2022;12:374.

42. Tapocik JD, Luu TV, Mayo CL, Wang B-D, Doyle E, Lee AD, et al. Neuroplasticity, axonal guidance and micro-RNA genes are associated with morphine self-administration behavior. Addiction Biology. 2013;18:480–495.

43. Berger AC, Whistler JL. Morphine-induced mu opioid receptor trafficking enhances reward yet prevents compulsive drug use. EMBO Mol Med. 2011;3:385–397.

44. Slivicki RA, Earnest T, Chang Y-H, Pareta R, Casey E, Li J-N, et al. Oral oxycodone self-administration leads to features of opioid misuse in male and female mice. Addict Biol. 2023;28:e13253.

45. Phillips AG, McGovern DJ, Lee S, Ro K, Huynh DT, Elvig SK, et al. Oral prescription opioid-seeking behavior in male and female mice. Addict Biol. 2020;25:e12828.

46. Moussawi K, Ortiz MM, Gantz SC, Tunstall BJ, Marchette RCN, Bonci A, et al. Fentanyl vapor self-administration model in mice to study opioid addiction. Sci Adv. 2020;6:eabc0413.

47. Corre J, van Zessen R, Loureiro M, Patriarchi T, Tian L, Pascoli V, et al. Dopamine neurons projecting to medial shell of the nucleus accumbens drive heroin reinforcement. Elife. 2018;7:e39945.

48. Corre J, van Zessen R, Loureiro M, Patriarchi T, Tian L, Pascoli V, et al. Dopamine neurons projecting to medial shell of the nucleus accumbens drive heroin reinforcement. ELife. 2018;7:e39945.

49. Pascoli V, Terrier J, Hiver A, Lüscher C. Sufficiency of Mesolimbic Dopamine Neuron Stimulation for the Progression to Addiction. Neuron. 2015;88:1054–1066.

50. Siuda ER, Copits BA, Schmidt MJ, Baird MA, Al-Hasani R, Planer WJ, et al. Spatiotemporal control of opioid signaling and behavior. Neuron. 2015;86:923–935.

51. Avey D, Sankararaman S, Yim AKY, Barve R, Milbrandt J, Mitra RD. Single-Cell RNA-Seq Uncovers a Robust Transcriptional Response to Morphine by Glia. Cell Rep. 2018;24:3619–3629.e4.

52. Liang J, Chen J-H, Chen X-H, Peng Y-H, Zheng X-G. Gene expression of conditioned locomotion and context-specific locomotor sensitization controlled by morphine-associated environment. Behav Brain Res. 2011;216:321–331.

53. Tapocik JD, Luu TV, Mayo CL, Wang B-D, Doyle E, Lee AD, et al. Neuroplasticity, axonal guidance and micro-RNA genes are associated with morphine self-administration behavior. Addict Biol. 2013;18:480–495.

54. Grice DE, Reenilä I, Männistö PT, Brooks AI, Smith GG, Golden GT, et al. Transcriptional profiling of C57 and DBA strains of mice in the absence and presence of morphine. BMC Genomics. 2007;8:76.

55. Ikeda H, Miyatake M, Koshikawa N, Ochiai K, Yamada K, Kiss A, et al. Morphine modulation of thrombospondin levels in astrocytes and its implications for neurite outgrowth and synapse formation. J Biol Chem. 2010;285:38415–38427.

56. Imperio CG, McFalls AJ, Colechio EM, Masser DR, Vrana KE, Grigson PS, et al. Assessment of individual differences in the rat nucleus accumbens transcriptome following taste-heroin extended access. Brain Res Bull. 2016;123:71–80.

57. Sillivan SE, Whittard JD, Jacobs MM, Ren Y, Mazloom AR, Caputi FF, et al. ELK1 transcription factor linked to dysregulated striatal mu opioid receptor signaling network and OPRM1 polymorphism in human heroin abusers. Biol Psychiatry. 2013;74:511–519.

58. Albertson DN, Schmidt CJ, Kapatos G, Bannon MJ. Distinctive profiles of gene expression in the human nucleus accumbens associated with cocaine and heroin abuse. Neuropsychopharmacology. 2006;31:2304–2312.

59. Zhang Y, Liang Y, Randesi M, Yuferov V, Zhao C, Kreek MJ. Chronic Oxycodone Self-administration Altered Reward-related Genes in the Ventral and Dorsal Striatum of C57BL/6J Mice: An RNA-seq Analysis. Neuroscience. 2018;393:333–349.

60. Zhang Y, Liang Y, Levran O, Randesi M, Yuferov V, Zhao C, et al. Alterations of expression of inflammation/immune-related genes in the dorsal and ventral striatum of adult C57BL/6J mice following chronic oxycodone self-administration: a RNA sequencing study. Psychopharmacology (Berl). 2017;234:2259–2275.

61. Yuferov V, Zhang Y, Liang Y, Zhao C, Randesi M, Kreek MJ. Oxycodone Self-Administration Induces Alterations in Expression of Integrin, Semaphorin and Ephrin Genes in the Mouse Striatum. Front Psychiatry. 2018;9:257.

62. Imperio CG, McFalls AJ, Hadad N, Blanco-Berdugo L, Masser DR, Colechio EM, et al. Exposure to environmental enrichment attenuates addiction-like behavior and alters molecular effects of heroin self-administration in rats. Neuropharmacology. 2018;139:26–40.

63. Yan B, Hu Z, Yao W, Le Q, Xu B, Liu X, et al. MiR-218 targets MeCP2 and inhibits heroin seeking behavior. Sci Rep. 2017;7:40413.

64. Zhang H, Wang Q, Wang Q, Liu A, Qin F, Sun Q, et al. Circular RNA expression profiling in the nucleus accumbens: Effects of electroacupuncture treatment on morphine-induced conditioned place preference. Addict Biol. 2020;25:e12794.

65. Sun H, Maze I, Dietz DM, Scobie KN, Kennedy PJ, Damez-Werno D, et al. Morphine epigenomically regulates behavior through alterations in histone H3 lysine 9 dimethylation in the nucleus accumbens. J Neurosci. 2012;32:17454–17464.

66. Martin JA, Caccamise A, Werner CT, Viswanathan R, Polanco JJ, Stewart AF, et al. A Novel Role for Oligodendrocyte Precursor Cells (OPCs) and Sox10 in Mediating Cellular and Behavioral Responses to Heroin. Neuropsychopharmacology. 2018;43:1385–1394.

67. Ammon-Treiber S, Tischmeyer H, Riechert U, Höllt V. Gene expression of transcription factors in the rat brain after morphine withdrawal. Neurochem Res. 2004;29:1267–1273.

68. Kuntz-Melcavage KL, Brucklacher RM, Grigson PS, Freeman WM, Vrana KE. Gene expression changes following extinction testing in a heroin behavioral incubation model. BMC Neurosci. 2009;10:95.

69. Ammon S, Mayer P, Riechert U, Tischmeyer H, Höllt V. Microarray analysis of genes expressed in the frontal cortex of rats chronically treated with morphine and after naloxone precipitated withdrawal. Brain Res Mol Brain Res. 2003;112:113–125.

70. Mendez EF, Wei H, Hu R, Stertz L, Fries GR, Wu X, et al. Angiogenic gene networks are dysregulated in opioid use disorder: evidence from multi-omics and imaging of postmortem human brain. Mol Psychiatry. 2021;26:7803–7812.

71. Jiang C, Wang X, Le Q, Liu P, Liu C, Wang Z, et al. Morphine coordinates SST and PV interneurons in the prelimbic cortex to disinhibit pyramidal neurons and enhance reward. Mol Psychiatry. 2021;26:1178–1193.

72. Nagamatsu ST, Rompala G, Hurd YL, Núñez-Rios DL, Montalvo-Ortiz JL, Traumatic Stress Brain Research Group. CpH methylome analysis in human cortical neurons identifies novel gene pathways and drug targets for opioid use disorder. Front Psychiatry. 2022;13:1078894.

73. Kozlenkov A, Jaffe AE, Timashpolsky A, Apontes P, Rudchenko S, Barbu M, et al. DNA Methylation Profiling of Human Prefrontal Cortex Neurons in Heroin Users Shows Significant Difference between Genomic Contexts of Hyper- and Hypomethylation and a Younger Epigenetic Age. Genes (Basel). 2017;8:152.

74. Rompala G, Nagamatsu ST, Martínez-Magaña JJ, Nuñez-Ríos DL, Wang J, Girgenti MJ, et al. Profiling neuronal methylome and hydroxymethylome of opioid use disorder in the human orbitofrontal cortex. Nat Commun. 2023;14:4544.

75. Liu A, Dai Y, Mendez EF, Hu R, Fries GR, Najera KE, et al. Genome-Wide Correlation of DNA Methylation and Gene Expression in Postmortem Brain Tissues of Opioid Use Disorder Patients. The International Journal of Neuropsychopharmacology. 2021;24:879–891.

76. Shu C, Sosnowski DW, Tao R, Deep-Soboslay A, Kleinman JE, Hyde TM, et al. Epigenome-wide study of brain DNA methylation following acute opioid intoxication. Drug and Alcohol Dependence. 2021;221:108658.

77. Loguinov AV, Anderson LM, Crosby GJ, Yukhananov RY. Gene expression following acute morphine administration. Physiol Genomics. 2001;6:169–181.

78. Choi MR, Jin Y-B, Bang SH, Im C-N, Lee Y, Kim H-N, et al. Age-related Effects of Heroin on Gene Expression in the Hippocampus and Striatum of Cynomolgus Monkeys. Clin Psychopharmacol Neurosci. 2020;18:93–108.

79. Skupio U, Sikora M, Korostynski M, Wawrzczak-Bargiela A, Piechota M, Ficek J, et al. Behavioral and transcriptional patterns of protracted opioid self-administration in mice. Addict Biol. 2017;22:1802–1816.

80. Piechota M, Korostynski M, Solecki W, Gieryk A, Slezak M, Bilecki W, et al. The dissection of transcriptional modules regulated by various drugs of abuse in the mouse striatum. Genome Biol. 2010;11:R48.

81. Korostynski M, Piechota M, Kaminska D, Solecki W, Przewlocki R. Morphine effects on striatal transcriptome in mice. Genome Biol. 2007;8:R128.

82. Piechota M, Korostynski M, Sikora M, Golda S, Dzbek J, Przewlocki R. Common transcriptional effects in the mouse striatum following chronic treatment with heroin and methamphetamine. Genes Brain Behav. 2012;11:404–414.

83. Korostynski M, Piechota M, Golda S, Przewlocki R. High-throughput gene expression profiling of opioid-induced alterations in discrete brain areas. Methods Mol Biol. 2015;1230:65–76.

84. Choi MR, Jin Y-B, Kim H-N, Chai YG, Im C-N, Lee S-R, et al. Gene expression in the striatum of cynomolgus monkeys after chronic administration of cocaine and heroin. Basic Clin Pharmacol Toxicol. 2021;128:686–698.

85. Cha HJ, Choi M-J, Ahn J-I, Jeon S-H, Kang H, Kim EJ, et al. Comparison of gene expression profiles in drug-withdrawn rats. Mol Cell Toxicol. 2016;12:197–207.

86. Valderrama-Carvajal A, Irizar H, Gago B, Jiménez-Urbieta H, Fuxe K, Rodríguez-Oroz MC, et al. Transcriptomic integration of D4R and MOR signaling in the rat caudate putamen. Sci Rep. 2018;8:7337.

87. Egervari G, Akpoyibo D, Rahman T, Fullard JF, Callens JE, Landry JA, et al. Chromatin accessibility mapping of the striatum identifies tyrosine kinase FYN as a therapeutic target for heroin use disorder. Nat Commun. 2020;11:4634.

88. Heller EA, Kaska S, Fallon B, Ferguson D, Kennedy PJ, Neve RL, et al. Morphine and cocaine increase serum- and glucocorticoid-inducible kinase 1 activity in the ventral tegmental area. J Neurochem. 2015;132:243–253.

89. McClung CA, Nestler EJ, Zachariou V. Regulation of gene expression by chronic morphine and morphine withdrawal in the locus ceruleus and ventral tegmental area. J Neurosci. 2005;25:6005–6015.

90. Zhang H, Wang Q, Sun Q, Qin F, Nie D, Li Q, et al. Effects of Compound 511 on BDNF-TrkB Signaling in the Mice Ventral Tegmental Area in Morphine-Induced Conditioned Place Preference. Cell Mol Neurobiol. 2021;41:961–975.

91. Jokinen V, Sidorova Y, Viisanen H, Suleymanova I, Tiilikainen H, Li Z, et al. Differential Spinal and Supraspinal Activation of Glia in a Rat Model of Morphine Tolerance. Neuroscience. 2018;375:10–24.

92. Cheng Y-C, Tsai R-Y, Sung Y-T, Chen I-J, Tu T-Y, Mao Y-Y, et al. Melatonin regulation of transcription in the reversal of morphine tolerance: Microarray analysis of differential gene expression. Int J Mol Med. 2019;43:791–806.

93. Shao J, Wang J, Huang J, Liu C, Pan Y, Guo Q, et al. Identification of lncRNA expression profiles and ceRNA analysis in the spinal cord of morphine-tolerant rats. Mol Brain. 2018;11:21.

94. Peng Y, Guo G, Shu B, Liu D, Su P, Zhang X, et al. Spinal CX3CL1/CX3CR1 May Not Directly Participate in the Development of Morphine Tolerance in Rats. Neurochem Res. 2017;42:3254– 3267.

95. Santiago JM, Rosas O, Torrado AI, González MM, Kalyan-Masih PO, Miranda JD. Molecular, anatomical, physiological, and behavioral studies of rats treated with buprenorphine after spinal cord injury. J Neurotrauma. 2009;26:1783–1793.

96. Weng Y, Wu J, Li L, Shao J, Li Z, Deng M, et al. Circular RNA expression profile in the spinal cord of morphine tolerated rats and screen of putative key circRNAs. Mol Brain. 2019;12:79.

97. Juul SE, Beyer RP, Bammler TK, Farin FM, Gleason CA. Effects of neonatal stress and morphine on murine hippocampal gene expression. Pediatr Res. 2011;69:285–292.

98. Marie-Claire C, Courtin C, Robert A, Gidrol X, Roques BP, Noble F. Sensitization to the conditioned rewarding effects of morphine modulates gene expression in rat hippocampus. Neuropharmacology. 2007;52:430–435.

99. Zheng H, Zeng Y, Zhang X, Chu J, Loh HH, Law P-Y. mu-Opioid receptor agonists differentially regulate the expression of miR-190 and NeuroD. Mol Pharmacol. 2010;77:102–109.

100. Befort K, Filliol D, Ghate A, Darcq E, Matifas A, Muller J, et al. Mu-opioid receptor activation induces transcriptional plasticity in the central extended amygdala. Eur J Neurosci. 2008;27:2973–2984.

101. Rodriguez Parkitna JM, Bilecki W, Mierzejewski P, Stefanski R, Ligeza A, Bargiela A, et al. Effects of morphine on gene expression in the rat amygdala. J Neurochem. 2004;91:38–48.

102. Zachariou V, Liu R, LaPlant Q, Xiao G, Renthal W, Chan GC, et al. Distinct roles of adenylyl cyclases 1 and 8 in opiate dependence: behavioral, electrophysiological, and molecular studies. Biol Psychiatry. 2008;63:1013–1021.

103. Saad MH, Rumschlag M, Guerra MH, Savonen CL, Jaster AM, Olson PD, et al. Differentially expressed gene networks, biomarkers, long noncoding RNAs, and shared responses with cocaine identified in the midbrains of human opioid abusers. Scientific Reports. 2019;9:1534.

104. Befort K, Filliol D, Darcq E, Ghate A, Matifas A, Lardenois A, et al. Gene expression is altered in the lateral hypothalamus upon activation of the mu opioid receptor. Ann N Y Acad Sci. 2008;1129:175–184.

105. Anghel A, Jamieson C a. M, Ren X, Young J, Porche R, Ozigbo E, et al. Gene expression profiling following short-term and long-term morphine exposure in mice uncovers genes involved in food intake. Neuroscience. 2010;167:554–566.

106. Hayase T, Yamamoto Y, Yamamoto K, Muso E, Shiota K. Microarray profile analysis of toxic cocaine-induced alterations in the expression of mouse brain gene sequences: a possible ‘protective’ effect of buprenorphine. J Appl Toxicol. 2004;24:15–20.

107. Hassan HE, Myers AL, Lee IJ, Chen H, Coop A, Eddington ND. Regulation of gene expression in brain tissues of rats repeatedly treated by the highly abused opioid agonist, oxycodone: microarray profiling and gene mapping analysis. Drug Metab Dispos. 2010;38:157–167.

108. Gaspari S, Purushothaman I, Cogliani V, Sakloth F, Neve RL, Howland D, et al. Suppression of RGSz1 function optimizes the actions of opioid analgesics by mechanisms that involve the Wnt/β-catenin pathway. Proc Natl Acad Sci U S A. 2018;115:E2085–E2094.

109. Toorie AM, Vassoler FM, Qu F, Schonhoff CM, Bradburn S, Murgatroyd CA, et al. A history of opioid exposure in females increases the risk of metabolic disorders in their future male offspring. Addict Biol. 2021;26:e12856.

110. Thibault K, Calvino B, Rivals I, Marchand F, Dubacq S, McMahon SB, et al. Molecular mechanisms underlying the enhanced analgesic effect of oxycodone compared to morphine in chemotherapy-induced neuropathic pain. PLoS One. 2014;9:e91297.

111. Delfs JM, Zhu Y, Druhan JP, Aston-Jones G. Noradrenaline in the ventral forebrain is critical for opiate withdrawal-induced aversion. Nature. 2000;403:430–434.

112. Zhu Y, Wienecke CF, Nachtrab G, Chen X. A thalamic input to the nucleus accumbens mediates opiate dependence. Nature. 2016;530:219–222.

113. Koob GF. The dark side of emotion: the addiction perspective. Eur J Pharmacol. 2015;753:73– 87.

114. Hearing M. Prefrontal-accumbens opioid plasticity: Implications for relapse and dependence. Pharmacol Res. 2019;139:158–165.

115. Lutz P-E, Kieffer BL. The multiple facets of opioid receptor function: implications for addiction. Curr Opin Neurobiol. 2013;23:473–479.

116. Lutz P-E, Almeida D, Filliol D, Jollant F, Kieffer BL, Turecki G. Increased functional coupling of the mu opioid receptor in the anterior insula of depressed individuals. Neuropsychopharmacology. 2021;46:920–927.

117. Lutz P-E, Kieffer BL. Opioid receptors: distinct roles in mood disorders. Trends Neurosci. 2013;36:195–206.

118. Kitchen I, Slowe SJ, Matthes HW, Kieffer B. Quantitative autoradiographic mapping of mu-, delta- and kappa-opioid receptors in knockout mice lacking the mu-opioid receptor gene. Brain Res. 1997;778:73–88.

119. Goody RJ, Oakley SM, Filliol D, Kieffer BL, Kitchen I. Quantitative autoradiographic mapping of opioid receptors in the brain of delta-opioid receptor gene knockout mice. Brain Res. 2002;945:9–19.

120. Slowe SJ, Simonin F, Kieffer B, Kitchen I. Quantitative autoradiography of mu-,delta- and kappa1 opioid receptors in kappa-opioid receptor knockout mice. Brain Res. 1999;818:335– 345.

121. Xia Y, Haddad GG. Ontogeny and distribution of opioid receptors in the rat brainstem. Brain Res. 1991;549:181–193.

122. Miyakawa T. No raw data, no science: another possible source of the reproducibility crisis. Mol Brain. 2020;13:24.

123. Field D, Sansone S-A, Collis A, Booth T, Dukes P, Gregurick SK, et al. Megascience. ‘Omics data sharing. Science. 2009;326:234–236.

124. FAIR Principles. GO FAIR. https://www.go-fair.org/fair-principles/. Accessed 14 February 2023.

125. NIH Genomic Data Sharing.

126. Murray JE, Belin-Rauscent A, Simon M, Giuliano C, Benoit-Marand M, Everitt BJ, et al. Basolateral and central amygdala differentially recruit and maintain dorsolateral striatum-dependent cocaine-seeking habits. Nat Commun. 2015;6:10088.

127. Zezula J, Casaccia-Bonnefil P, Ezhevsky SA, Osterhout DJ, Levine JM, Dowdy SF, et al. p21cip1 is required for the differentiation of oligodendrocytes independently of cell cycle withdrawal. EMBO Rep. 2001;2:27–34.

128. Scholpa NE, Briggs SB, Wagner JJ, Cummings BS. Cyclin-Dependent Kinase Inhibitor 1a (p21) Modulates Response to Cocaine and Motivated Behaviors. J Pharmacol Exp Ther. 2016;357:56– 65.

129. Häusl AS, Brix LM, Hartmann J, Pöhlmann ML, Lopez J-P, Menegaz D, et al. The co-chaperone Fkbp5 shapes the acute stress response in the paraventricular nucleus of the hypothalamus of male mice. Mol Psychiatry. 2021;26:3060–3076.

130. Levran O, Peles E, Randesi M, Li Y, Rotrosen J, Ott J, et al. Stress-related genes and heroin addiction: a role for a functional FKBP5 haplotype. Psychoneuroendocrinology. 2014;45:67–76.

131. Cahill KM, Huo Z, Tseng GC, Logan RW, Seney ML. Improved identification of concordant and discordant gene expression signatures using an updated rank-rank hypergeometric overlap approach. Scientific Reports. 2018;8:9588.

132. Al-Hasani R, Bruchas MR. Molecular mechanisms of opioid receptor-dependent signaling and behavior. Anesthesiology. 2011;115:1363–1381.

133. Nectow AR, Moya MV, Ekstrand MI, Mousa A, McGuire KL, Sferrazza CE, et al. Rapid Molecular Profiling of Defined Cell Types Using Viral TRAP. Cell Rep. 2017;19:655–667.

134. Heiman M, Schaefer A, Gong S, Peterson JD, Day M, Ramsey KE, et al. A translational profiling approach for the molecular characterization of CNS cell types. Cell. 2008;135:738–748.

135. Bailly J, Del Rossi N, Runtz L, Li J-J, Park D, Scherrer G, et al. Targeting morphine-responsive neurons: generation of a knock-in mouse line expressing Cre recombinase from the mu opioid receptor gene locus. ENeuro. 2020. 7 May 2020. https://doi.org/10.1523/ENEURO.0433-19.2020.

136. Okunomiya T, Hioki H, Nishimura C, Yawata S, Imayoshi I, Kageyama R, et al. Generation of a MOR-CreER knock-in mouse line to study cells and neural circuits involved in mu opioid receptor signaling. Genesis. 2020;58:e23341.

137. Märtin A, Calvigioni D, Tzortzi O, Fuzik J, Wärnberg E, Meletis K. A Spatiomolecular Map of the Striatum. Cell Rep. 2019;29:4320–4333.e5.

138. Macosko EZ, Basu A, Satija R, Nemesh J, Shekhar K, Goldman M, et al. Highly Parallel Genome-wide Expression Profiling of Individual Cells Using Nanoliter Droplets. Cell. 2015;161:1202– 1214.

139. Stuart T, Butler A, Hoffman P, Hafemeister C, Papalexi E, Mauck WM, et al. Comprehensive Integration of Single-Cell Data. Cell. 2019;177:1888–1902.e21.

140. Chari T, Pachter L. The Specious Art of Single-Cell Genomics. Genomics; 2021.

141. Gibson G. Perspectives on rigor and reproducibility in single cell genomics. PLoS Genet. 2022;18:e1010210.

142. Schmid KT, Höllbacher B, Cruceanu C, Böttcher A, Lickert H, Binder EB, et al. scPower accelerates and optimizes the design of multi-sample single cell transcriptomic studies. Nat Commun. 2021;12:6625.

143. Browne CJ, Godino A, Salery M, Nestler EJ. Epigenetic Mechanisms of Opioid Addiction. Biol Psychiatry. 2020;87:22–33.

144. Belzeaux R, Lalanne L, Kieffer BL, Lutz PE. Focusing on the Opioid System for Addiction Biomarker Discovery. Trends Mol Med. 2018;24:206–220.

145. Wu Q, Hwang CK, Zheng H, Wagley Y, Lin H-Y, Kim DK, et al. MicroRNA 339 down-regulates μ- opioid receptor at the post-transcriptional level in response to opioid treatment. FASEB J. 2013;27:522–535.

146. Ebrahimi G, Asadikaram G, Akbari H, Nematollahi MH, Abolhassani M, Shahabinejad G, et al. Elevated levels of DNA methylation at the OPRM1 promoter region in men with opioid use disorder. Am J Drug Alcohol Abuse. 2017:1–7.

147. Nielsen DA, Yuferov V, Hamon S, Jackson C, Ho A, Ott J, et al. Increased OPRM1 DNA methylation in lymphocytes of methadone-maintained former heroin addicts. Neuropsychopharmacology. 2009;34:867–873.

148. Doehring A, Oertel BG, Sittl R, Lotsch J. Chronic opioid use is associated with increased DNA methylation correlating with increased clinical pain. Pain. 2013;154:15–23.

149. Barrow TM, Byun HM, Li X, Smart C, Wang YX, Zhang Y, et al. The effect of morphine upon DNA methylation in ten regions of the rat brain. Epigenetics. 2017;12:1038–1047.

150. Lutz P-E, Chay M-A, Pacis A, Chen GG, Aouabed Z, Maffioletti E, et al. Non-CG methylation and multiple histone profiles associate child abuse with immune and small GTPase dysregulation. Nat Commun. 2021;12:1132.

151. Lutz P-E, Mechawar N, Turecki G. Neuropathology of suicide: recent findings and future directions. Mol Psychiatry. 2017;22:1395–1412.

152. Feng J, Shao N, Szulwach KE, Vialou V, Huynh J, Zhong C, et al. Role of Tet1 and 5-hydroxymethylcytosine in cocaine action. Nat Neurosci. 2015;18:536–544.

153. Benelli M, Franceschini GM, Magi A, Romagnoli D, Biagioni C, Migliaccio I, et al. Charting differentially methylated regions in cancer with Rocker-meth. Commun Biol. 2021;4:1249.

154. Kaluscha S, Domcke S, Wirbelauer C, Stadler MB, Durdu S, Burger L, et al. Evidence that direct inhibition of transcription factor binding is the prevailing mode of gene and repeat repression by DNA methylation. Nat Genet. 2022;54:1895–1906.

155. Feng J, Wilkinson M, Liu X, Purushothaman I, Ferguson D, Vialou V, et al. Chronic cocaine-regulated epigenomic changes in mouse nucleus accumbens. Genome Biol. 2014;15:R65.

156. Ching T, Huang S, Garmire LX. Power analysis and sample size estimation for RNA-Seq differential expression. RNA. 2014;20:1684–1696.

157. Baccarella A, Williams CR, Parrish JZ, Kim CC. Empirical assessment of the impact of sample number and read depth on RNA-Seq analysis workflow performance. BMC Bioinformatics. 2018;19:423.

158. Hoffman GE, Jaffe AE, Gandal MJ, Collado-Torres L, Sieberts SK, Devlin B, et al. Comment on: What genes are differentially expressed in individuals with schizophrenia? A systematic review. Mol Psychiatry. 2023;28:523–525.

159. Khan AH, Bagley JR, LaPierre N, Gonzalez-Figueroa C, Spencer TC, Choudhury M, et al. Genetic pathways regulating the longitudinal acquisition of cocaine self-administration in a panel of inbred and recombinant inbred mice. Cell Rep. 2023;42:112856.

160. Hekman JP, Johnson JL, Kukekova AV. Transcriptome Analysis in Domesticated Species: Challenges and Strategies. Bioinform Biol Insights. 2015;9:21–31.

161. Williams AG, Thomas S, Wyman SK, Holloway AK. RNA-seq Data: Challenges in and Recommendations for Experimental Design and Analysis. Curr Protoc Hum Genet. 2014;83:11.13.1-20.

162. Crist RC, Reiner BC, Berrettini WH. A review of opioid addiction genetics. Curr Opin Psychol. 2019;27:31–35.

163. scorch - Home. https://scorch.igs.umaryland.edu/. Accessed 19 May 2023.

164. Barrett T, Wilhite SE, Ledoux P, Evangelista C, Kim IF, Tomashevsky M, et al. NCBI GEO: archive for functional genomics data sets--update. Nucleic Acids Res. 2013;41:D991-995.

165. Data Portal | IHEC. https://epigenomesportal.ca/ihec/. Accessed 19 May 2023.

166. Savage CJ, Vickers AJ. Empirical study of data sharing by authors publishing in PLoS journals. PLoS One. 2009;4:e7078.

167. Piron A, Szymczak F, Alvelos MI, Defrance M, Lenaerts T, Eizirik DL, et al. RedRibbon: A new rank-rank hypergeometric overlap pipeline to compare gene and transcript expression signatures. Bioinformatics; 2022.

168. Deroche-Gamonet V, Belin D, Piazza PV. Evidence for addiction-like behavior in the rat. Science. 2004;305:1014–1017.

169. Domingo-Rodriguez L, Ruiz de Azua I, Dominguez E, Senabre E, Serra I, Kummer S, et al. A specific prelimbic-nucleus accumbens pathway controls resilience versus vulnerability to food addiction. Nat Commun. 2020;11:782.

170. Laine MA, Trontti K, Misiewicz Z, Sokolowska E, Kulesskaya N, Heikkinen A, et al. Genetic Control of Myelin Plasticity after Chronic Psychosocial Stress. ENeuro. 2018;5:ENEURO.0166-18.2018.

171. Bagot RC, Cates HM, Purushothaman I, Lorsch ZS, Walker DM, Wang J, et al. Circuit-wide Transcriptional Profiling Reveals Brain Region-Specific Gene Networks Regulating Depression Susceptibility. Neuron. 2016;90:969–983.

172. Roy B, Wang Q, Dwivedi Y. Long Noncoding RNA-Associated Transcriptomic Changes in Resiliency or Susceptibility to Depression and Response to Antidepressant Treatment. Int J Neuropsychopharmacol. 2018;21:461–472.

173. Cervera-Juanes R, Wilhelm LJ, Park B, Grant KA, Ferguson B. Alcohol-dose-dependent DNA methylation and expression in the nucleus accumbens identifies coordinated regulation of synaptic genes. Transl Psychiatry. 2017;7:e994.

174. Domingo-Rodriguez L, Cabana-Domínguez J, Fernàndez-Castillo N, Cormand B, Martín-García E, Maldonado R. Differential expression of miR-1249-3p and miR-34b-5p between vulnerable and resilient phenotypes of cocaine addiction. Addict Biol. 2022;27:e13201.

175. Ekstrand MI, Nectow AR, Knight ZA, Latcha KN, Pomeranz LE, Friedman JM. Molecular profiling of neurons based on connectivity. Cell. 2014;157:1230–1242.

176. Zhang Z, Zhou J, Tan P, Pang Y, Rivkin AC, Kirchgessner MA, et al. Epigenomic diversity of cortical projection neurons in the mouse brain. Nature. 2021;598:167–173.

177. Rodriques SG, Stickels RR, Goeva A, Martin CA, Murray E, Vanderburg CR, et al. Slide-seq: A scalable technology for measuring genome-wide expression at high spatial resolution. Science. 2019;363:1463–1467.

178. Eng C-HL, Lawson M, Zhu Q, Dries R, Koulena N, Takei Y, et al. Transcriptome-scale super-resolved imaging in tissues by RNA seqFISH. Nature. 2019;568:235–239.

179. Borm LE, Mossi Albiach A, Mannens CCA, Janusauskas J, Özgün C, Fernández-García D, et al. Scalable in situ single-cell profiling by electrophoretic capture of mRNA using EEL FISH. Nat Biotechnol. 2022. 22 September 2022. https://doi.org/10.1038/s41587-022-01455-3.

180. Armand EJ, Li J, Xie F, Luo C, Mukamel EA. Single-Cell Sequencing of Brain Cell Transcriptomes and Epigenomes. Neuron. 2021;109:11–26.

181. Mendez EF, Wei H, Hu R, Stertz L, Fries GR, Wu X, et al. Angiogenic gene networks are dysregulated in opioid use disorder: evidence from multi-omics and imaging of postmortem human brain. Molecular Psychiatry. 2021;26:7803–7812.

182. Li KW, Jimenez CR, van der Schors RC, Hornshaw MP, Schoffelmeer ANM, Smit AB. Intermittent administration of morphine alters protein expression in rat nucleus accumbens. Proteomics. 2006;6:2003–2008.

183. Van den Oever MC, Lubbers BR, Goriounova NA, Li KW, Van der Schors RC, Loos M, et al. Extracellular matrix plasticity and GABAergic inhibition of prefrontal cortex pyramidal cells facilitates relapse to heroin seeking. Neuropsychopharmacology. 2010;35:2120–2133.

184. Bu Q, Yang Y, Yan G, Hu Z, Hu C, Duan J, et al. Proteomic analysis of the nucleus accumbens in rhesus monkeys of morphine dependence and withdrawal intervention. J Proteomics. 2012;75:1330–1342.

185. Mokhtari A, Porte B, Belzeaux R, Etain B, Ibrahim EC, Marie-Claire C, et al. The molecular pathophysiology of mood disorders: From the analysis of single molecular layers to multi-omic integration. Prog Neuropsychopharmacol Biol Psychiatry. 2022;116:110520.

186. Li CX, Wheelock CE, Sköld CM, Wheelock ÅM. Integration of multi-omics datasets enables molecular classification of COPD. Eur Respir J. 2018;51:1701930.

187. Yang M, Matan-Lithwick S, Wang Y, De Jager PL, Bennett DA, Felsky D. Multi-omic integration via similarity network fusion to detect molecular subtypes of ageing. Brain Commun. 2023;5:fcad110.

188. Revilla L, Mayorgas A, Corraliza AM, Masamunt MC, Metwaly A, Haller D, et al. Multi-omic modelling of inflammatory bowel disease with regularized canonical correlation analysis. PLoS One. 2021;16:e0246367.

189. Hasin Y, Seldin M, Lusis A. Multi-omics approaches to disease. Genome Biol. 2017;18:83.

190. Liberati A, Altman DG, Tetzlaff J, Mulrow C, Gotzsche PC, Ioannidis JPA, et al. The PRISMA statement for reporting systematic reviews and meta-analyses of studies that evaluate healthcare interventions: explanation and elaboration. BMJ. 2009;339:b2700–b2700.

