## Supplementary Material (Methods, Supplementary Figures) for "Functional genomic mechanisms of opioid action and opioid use disorder: a systematic review of animal models and human studies"

**Corresponding author**:

**Supplementary Material**

**1. Material and methods**

**2. Legends of supplementary Figure and Tables**

#### 1. Material and methods

***Identification of studies***. Relevant studies were identified based on the PRISMA methodology (Preferred Reporting Items for Systematic Reviews and Meta-Analyses[1]). The review protocol was registered prospectively in Prospero (registration number: CRD42022270113), and 3 databases were used for systematic screening (MEDLINE, Embase and Web of Science). The literature search was performed until September 2021, using the following keywords: “microarray” OR “RNA sequencing” OR “bisulfite sequencing” OR “chromatin immunopurification sequencing” OR “single cell RNAsequencing” AND each of the following terms: “opiates”, “opioid”, “morphine”, “fentanyl”, “oxycodone”, “heroin”, “methadone” or “buprenorphine”. References were imported to Zotero, then titles and abstract were screened to remove duplicates.

***Eligibility***. Articles were deemed eligible if they used a high-throughput methodology to assess genome-wide modifications of gene expression or epigenetic mechanisms as a function of opioid exposure, in the nervous system of primates or rodents. They had to be published in English in peer-reviewed journals. Exclusion criteria included tissue other than the nervous system, and candidate genes approaches. Additional studies that could not be identified using aforementioned keywords, but nonetheless were known to the authors (preprint servers, curation of references from eligible studies, etc) and met eligibility criteria, were also included. Of note, human genome-wide association studies of genetic variation associated with OUD were not included, as recent reviews are available on this topic[2, 3], and our primary focus was on functional genomic mechanisms.

***Quantitative analysis***. In order to run a quantitative analysis focusing on the 5 most frequently investigated structures, all DEG reported in each study were used, regardless of additional selection performed. The full code to reproduce all results is available as Supplementary File 1.

***RRHO2.*** We retrieved the whole-genome transcriptomic data (p-values and fold-changes) for the Seney et al[4] and Townsend et al[5] studies. The Seney dataset was re-analyzed to identify separately transcriptomic adaptations associated with OUD in males or females. Both datasets were filtered to keep only orthologous genes that had p-values and fold-changes reported in both studies. RRHO2 (Rank-Rank Hypergeometric Overlap[6]) was then conducted using the "RRHO2" R package (v.1.0; see Supplementary File for full code). Genes are ranked by p-value and by direction of fold-change for both studies, which defines 4 quadrants: in the bottom-left one genes are up-regulated in both studies, in the top-right one genes are down-regulated in both studies (concordance between datasets), the two other quadrants describing genes that are up-regulated in one study and down-regulated in the other (discordance between datasets).

***Comparison of single-cell studies.*** From the 2 single cell studies (Avey et al[7] and Reiner et al[8]), we gathered information regarding differentially expressed genes (DEG) by cluster, as well as genes used to define cluster identity in terms of cell-types. DEG by clusters were extracted from Supplementary Table 3 (sheet "Cluster-specific SCDE") and from Supplementary Table 4 in Avey et al and Reiner et al, respectively. Marker genes used to define cell-type identity were extracted from Supplementary Table 2 (sheet entitled "Neuronal Cluster-enriched Genes") and from Supplementary Table 10 in Avey et al and Reiner et al, respectively. We then compared the lists of DEG and of marker genes for both studies, cluster by cluster, using pairwise Jaccard indices. Those were then used to compute a correlation between DEG and marker genes Jaccard indices (Fig.S4B).

***Gene orthologs.*** To compare lists of genes across studies conducted in different species, orthologous genes were identified using BioMart (see Supplementary File 1 for code).

***Statistical analyses.*** Statistical analyses were performed in R. Data are expressed as mean±SEM, with statistical significance set as *p<0.05, **p<0.01, ***p<0.001. A Mann-Whitney test was used to compare Jaccard indices within (“intra”) or across (“inter”) factor levels, for each potential source of variability among studies.

***Code availability.*** All code used in the present review is available as Supplementary File 1.

**2. Legends of supplementary Figure and Tables**

**Supplementary Figure 1**. **Quantitative analyses in the whole striatum.** Number of differentially expressed genes identified in the whole striatum by individual studies, or across several ones, among those that used chronic (A) or acute (B) opioid administration. Among chronic morphine studies, the JI varied from 0.1% (Coffey 2020 versus Skupio 2017), 1.2% (Korostynski 2007/Skupio 2017) to 2.6% (Korostynski 2007/Skupio 2017). Among acute morphine studies, from 2.7% to 7.8%: Piechota 2010/Skupio 2017, JI=2.7%; Piechota 2010/Korostynski 2007, 3.2%; Piechota 2010/Korostynski 2013, 6.4%; Korostynski 2013/Skupio 2017, 6.6%; Korostynski 2013/Korostynski 2007, 7.3%; Korostynski 2007/Skupio 2017, 7.8%. Only one comparison stood out among acute heroin studies (30 common DEG, JI=19%, Piechota 2010/Korostinsksy 2013). For clarity, only intersection sizes higher than or equal to 4 are represented in (B).

**Supplementary figure 2. Comparison of directionality for differentially expressed genes (DEG) identified by at least 5 studies.** Here we considered the directionality (up/down) of DEG associated with opioid exposure, focusing on genes identified across at least 5 studies. Among the 17/24 studies that reported on directionality, most genes showed discordant results (37/63, 59%): for example, Cables1 was upregulated in Imperio et al (nucleus accumbens) and Coffey et al (whole striatum), but downregulated in Seney et al in both the frontal cortex and NAc. Studies are arranged by brain structure, while genes are arranged by percentage of concordance (for example, a gene which was found as upregulated in two studies and as downregulated in one will be marked as 66%). 17 genes were upregulated: Cdkn1a (6 studies showing upregulation), Slc2a1 (5), Sult1a1 (5), Arrdc3 (4), Ccdc117 (4), Fkpb5 (4), Plin4 (4), Arid5b (3), Pla2g3 (3), Tsc22d3 (3), Dusp1 (2), Errfi1 (2), Map3k6 (2), Pdk4 (2), Sgk1 (2), Spsb1 (2), Xdh (2). One gene was downregulated: Wscd1 (4). Among these, 11 genes were identified by at least 3 studies: i) 3 of these, Cdkn1a, Fkbp5 and Slc2a1, have been previously investigated at functional or mechanistic level in relation to opioid use disorder (see main text); for the other 8 (Arid5b, Arrdc3, Ccdc117, Pla2g3, Plin4, Sult1a1, Tsc22d3 and Wscd1), comparatively less data is currently available in relation to those specific drugs of abuse, as summarized briefly below. Arid5b encodes a DNA binding protein that forms a histone H3K9Me2 demethylase complex. It has been implicated in B-lymphocytes differentiation and acute lymphoblastic leukemia, with one genetic study reporting an association with alcohol and other substance use disorders[9]. Arrdc3 encodes an α-arrestin regulating GPCR signalling. It has been associated with body mass index in humans, and its absence in mice is protective against obesity[10], suggesting a role in metabolism regulation. A few studies have started to investigate its role in cancer[11, 12]. Ccdc117 encodes a coiled-coil domain-containing protein whose function is only starting to be characterized, in relation to cell cycle progression and DNA synthesis and repair[13], anxiety disorders[14] or, more recently, stress resilience[15]. Pla2g3 is a phospholipase, involved in lipid metabolism, ciliogenesis and cancer, that has been identified in a microarray analysis of midbrain dopamine neurons upon gestational nicotine exposure[16], as well as a RNA-Seq comparison of selectively bred high and low alcohol-preferring mice[17]. Plin4 encodes a perilipin protein that regulates lipid homeostasis by coating intracellular lipid droplets. It is a glucocorticoid responsive gene[18, 19] that might participate in the pathophysiology of neurodegenerative diseases such as the amyotrophic lateral sclerosis[20] and Parkinson disease[21]. Sult1a1 encodes an enzyme of the sulfotransferase family that catalyses the sulfation of multiple endogenous compounds and drugs[22]. It is also a glucocorticoid-responsive gene[23, 24] that has been linked to opioid metabolism[25], alcohol consumption[26], nicotine addiction[27] and lifetime cannabis consumption[28]. Tsc22d3 is an anti-inflammatory effector of glucocorticoid signaling, highlighted in a recent re-analysis[29] of 3 of the mouse studies included in the present systematic review[30–32], and also associated with ketamine action. Finally, Wscd1 is highly expressed in the brain, predicted to enable sulfotransferase activity and located in membranes, and has recently been identified in a longitudinal Epigenome-Wide Association Study of sensation-seeking, a behavioral trait considered as an endophenotype for substance use disorders[33]. The other genes didn't show concordant dysregulations across studies, or direction was provided by one study, or none. Studies considered : 1 : Lefevre et al (dorsal striatum), 2 : Valderrama-Carjaval et al, 3 : Kuntz-Melcavage et al, 4 : Liu et al, 5 : Mendez et al, 6 : Seney et al (frontal cortex), 7 : Albertson et al, 8 : Grice et al, 9 : Imperio et al (Large Suppressor), 10 : Imperio et al (Small Suppressor), 11 : Lefevre et al (nucleus accumbens), 12 : Seney et al (nucleus accumbens), 13 : Townsend et al, 14 : Ye et al, 15 : Coffey et al, 16 : Korostynski et al 2007 (acute), 17 : Korostynski et al 2007 (chronic), 18 : Korostynski et al 2013 (heroin), 19 : Korostynski et al 2013 (morphine), 20 : Piechota et al 2012, 21 : Piechota et al 2010 (heroin), 22 : Piechota et al 2010 (morphine), 23 : Skupio et al (acute), 24 : Skupio et al (chronic). Gene Names : Cdkn1a : Cyclin dependent kinase inhibitor 1A, Slc2a1 : Solute carrier family 2 member 1, Sult1a1 : Sulfotransferase family 1A member 1, Arrdc3 : Arrestin domain containing 3, Ccdc117 : Coiled-coil domain containing 117, Fkbp5 : FKBP prolyl isomerase 5, Plin4 : Perilipin 4, Wscd1 : WSC domain containing 1, Arid5b : AT-rich interaction domain 5B, Pla2g3 : Phospholipase A2 group III, Tsc22d3 : TSC22 domain family member 3, Dusp1 : Dual specificity phosphatase 1, Errfi1 : ERBB receptor feedback inhibitor 1, Map3k6 : Mitogen-activated protein kinase kinase kinase 6, Pdk4 : Pyruvate dehydrogenase kinase 4, Sgk1 : Serum/glucocorticoid regulated kinase 1, Spsb1 : SplA/ryanodine receptor domain and SOCS box containing 1, Xdh : Xanthine dehydrogenase, Avil : Advillin, Emx2 : Empty spiracles homeobox 2, Etnppl : Ethanolamine-phosphate phospho-lyase, Fam107a : Family with sequence similarity 107 member A, Fam163a : Family with sequence similarity 163 member A, Gjb6 : Gap junction protein beta 6, Hmgcs1 : 3-hydroxy-3-methylglutaryl-CoA synthase 1, Kcng1 : Potassium voltage-gated channel modifier subfamily G member 1, Cx3cr1 : C-X3-C motif chemokine receptor 1, Dusp6 : Dual specificity phosphatase 6, Fn1 : Fibronectin 1, Heph : Hephaestin, Net1 : Neuroepithelial cell transforming 1, Opalin : Oligodendrocytic myelin paranodal and inner loop protein, Oscp1 : Organic solute carrier partner 1, Phyhd1 : Phytanol-CoA dioxygenase domain containing 1, Pnpla2 : Patatin like phospholipase domain containing 2, Angptl4 : Angiopoietin like 4, Mat2a : Methionine adenosyltransferase 2A, Mcam : Melanoma cell adhesion molecule, Rasl11a : RAS like family 11 member A, Tfcp2l1 : Transcription factor CP2 like 1, Txnip : Thioredoxin interacting protein, Camk1g : Calcium/calmodulin dependent protein kinase IG, Cryab : Crystallin alpha B, Dbp : D-box binding PAR BZIP transcription factor, Enho : Energy homeostasis associated, Irf2bpl : Interferon regulatory factor 2 binding protein like, Irs1 : Insulin receptor substrate 1, Npy : Neuropeptide Y, Pdzd2 : PDZ domain containing 2, Rprml : Reprimo like, Slc6a11 : Solute carrier family 6 member 11, Cables1 : Cdk5 and Abl enzyme substrate 1, Cyp2d22 : Cytochrome P450 family 2 subfamily D member 22, Homer1 : Homer scaffold protein 1, Tnfrsf25 : TNF receptor superfamily member 25, Cldn5 : Claudin 5, Ddit4 : DNA damage inducible transcript 4, Hif3a : Hypoxia inducible factor 3 subunit alpha, Plekhf1 : Pleckstrin homology and FYVE domain containing 1, Rasd1 : Ras related dexamethasone induced 1, Arrdc2 : Arrestin domain containing 2, Itgad : Integrin subunit alpha D, Polr3e : RNA polymerase III subunit E.

**Supplementary Figure 3. Comparison of opioid-induced gene expression changes (Seney et al and Townsend et al) (A)** Venn diagram of differentially expressed genes in Townsend et al, 2021 and in Seney et al, 2021, for pooled women and men against female rats. **(B)** Venn diagram of differentially expressed genes in Townsend et al, 2021 and in Seney et al, 2021, for pooled women and men against male rats.

**Supplementary Figure 4. Cross-comparison of 2 single-cell RNA-sequencing studies (Avey et al and Reiner et al) (A)** Heatmap of pairwise Jaccard indexes for marker genes (genes used to assign clusters to cell types) across cell types in Avey et al and Reiner et al. **(B)** Correlation of Jaccard indexes for marker genes and Jaccard indexes for differentially expressed genes. Note: only n=8 and n=9 cell types had both information for marker genes and differentially expressed genes, 3 and 2 cell types only had information for marker genes, and 18 and 8 cell types only had information for differentially expressed genes (in Avey et al and Reiner et al, respectively).

**Suppl Table 1**. Transcriptomic studies and their characteristics. **(A)** Transcriptomic studies that were included in quantitative analysis and their various thresholds for listing a gene as differentially expressed. **(B)** Transcriptomic studies that were excluded from the quantitative analysis, with reason for exclusion.

**Suppl Table 2.** Matrix of pairwise Jaccard indexes (%) of differentially expressed genes for all studies included in quantitative analysis.

**Suppl Table 3.** Lists of DEGs identified by 1 or more studies included in quantitative analysis.

**Suppl Table 4.** Results of Gene Ontology enrichment analysis for DEG identified by at least 5 studies included in quantitative analysis.

**Suppl Table 5.** Results of Gene Ontology enrichment analysis for DEG identified by at least 4 studies included in quantitative analysis.

**Suppl Table 6.** Results of Gene Ontology enrichment analysis for DEG identified by at least 3 studies included in quantitative analysis.

#### References


1. Liberati A, Altman DG, Tetzlaff J, Mulrow C, Gotzsche PC, Ioannidis JPA, et al. The PRISMA statement for reporting systematic reviews and meta-analyses of studies that evaluate healthcare interventions: explanation and elaboration. BMJ. 2009;339:b2700–b2700.
2. Reed B, Kreek MJ. Genetic Vulnerability to Opioid Addiction. Cold Spring Harb Perspect Med. 2021;11:a039735.
3. Crist RC, Reiner BC, Berrettini WH. A review of opioid addiction genetics. Curr Opin Psychol. 2019;27:31–35.
4. Seney ML, Kim S-M, Glausier JR, Hildebrand MA, Xue X, Zong W, et al. Transcriptional Alterations in Dorsolateral Prefrontal Cortex and Nucleus Accumbens Implicate Neuroinflammation and Synaptic Remodeling in Opioid Use Disorder. Biological Psychiatry. 2021:S000632232101369X.
5. Townsend EA, Kim RK, Robinson HL, Marsh SA, Banks ML, Hamilton PJ. Opioid withdrawal produces sex-specific effects on fentanyl-vs.-food choice and mesolimbic transcription. Biol Psychiatry Glob Open Sci. 2021;1:112–122.
6. Cahill KM, Huo Z, Tseng GC, Logan RW, Seney ML. Improved identification of concordant and discordant gene expression signatures using an updated rank-rank hypergeometric overlap approach. Sci Rep. 2018;8:9588.
7. Avey D, Sankararaman S, Yim AKY, Barve R, Milbrandt J, Mitra RD. Single-Cell RNA-Seq Uncovers a Robust Transcriptional Response to Morphine by Glia. Cell Rep. 2018;24:3619-3629 e4.
8. Reiner BC, Zhang Y, Stein LM, Perea ED, Arauco-Shapiro G, Ben Nathan J, et al. Single nucleus transcriptomic analysis of rat nucleus accumbens reveals cell type-specific patterns of gene expression associated with volitional morphine intake. Transl Psychiatry. 2022;12:374.
9. Peng Q, Wilhelmsen KC, Ehlers CL. Common genetic substrates of alcohol and substance use disorder severity revealed by pleiotropy detection against GWAS catalog in two populations. Addict Biol. 2021;26:e12877.

10. Patwari P, Emilsson V, Schadt EE, Chutkow WA, Lee S, Marsili A, et al. The arrestin domain-containing 3 protein regulates body mass and energy expenditure. Cell Metab. 2011;14:671–683.

11. Wedegaertner H, Pan W-A, Gonzalez CC, Gonzalez DJ, Trejo J. The α-Arrestin ARRDC3 Is an Emerging Multifunctional Adaptor Protein in Cancer. Antioxid Redox Signal. 2022;36:1066–1079.

12. Arakaki AKS, Pan W-A, Lin H, Trejo J. The α-arrestin ARRDC3 suppresses breast carcinoma invasion by regulating G protein-coupled receptor lysosomal sorting and signaling. J Biol Chem. 2018;293:3350–3362.

13. Horton AJ, Brooker J, Streitfeld WS, Flessa ME, Pillai B, Simpson R, et al. Nkx2-5 Second Heart Field Target Gene Ccdc117 Regulates DNA Metabolism and Proliferation. Sci Rep. 2019;9:1738.

14. Hettema JM, Webb BT, Guo A-Y, Zhao Z, Maher BS, Chen X, et al. Prioritization and association analysis of murine-derived candidate genes in anxiety-spectrum disorders. Biol Psychiatry. 2011;70:888–896.

15. Lorsch ZS, Hamilton PJ, Ramakrishnan A, Parise EM, Salery M, Wright WJ, et al. Stress resilience is promoted by a Zfp189-driven transcriptional network in prefrontal cortex. Nat Neurosci. 2019;22:1413–1423.

16. Kanlikilicer P, Zhang D, Dragomir A, Akay YM, Akay M. Gene expression profiling of midbrain dopamine neurons upon gestational nicotine exposure. Med Biol Eng Comput. 2017;55:467–482.

17. Grecco GG, Haggerty DL, Doud EH, Fritz BM, Yin F, Hoffman H, et al. A multi-omic analysis of the dorsal striatum in an animal model of divergent genetic risk for alcohol use disorder. J Neurochem. 2021;157:1013–1031.

18. Buurstede JC, Umeoka EHL, da Silva MS, Krugers HJ, Joëls M, Meijer OC. Application of a pharmacological transcriptome filter identifies a shortlist of mouse glucocorticoid receptor target genes associated with memory consolidation. Neuropharmacology. 2022;216:109186.

19. Vanderhaeghen T, Timmermans S, Watts D, Paakinaho V, Eggermont M, Vandewalle J, et al. Reprogramming of glucocorticoid receptor function by hypoxia. EMBO Rep. 2022;23:e53083.

20. Zhu L, Hu F, Li C, Zhang C, Hang R, Xu R. Perilipin 4 Protein: an Impending Target for Amyotrophic Lateral Sclerosis. Mol Neurobiol. 2021;58:1723–1737.

21. Han X, Zhu J, Zhang X, Song Q, Ding J, Lu M, et al. Plin4-Dependent Lipid Droplets Hamper Neuronal Mitophagy in the MPTP/p-Induced Mouse Model of Parkinson's Disease. Front Neurosci. 2018;12:397.

22. Gamage N, Barnett A, Hempel N, Duggleby RG, Windmill KF, Martin JL, et al. Human sulfotransferases and their role in chemical metabolism. Toxicol Sci. 2006;90:5–22.

23. Juszczak GR, Stankiewicz AM. Glucocorticoids, genes and brain function. Prog Neuropsychopharmacol Biol Psychiatry. 2018;82:136–168.

24. Duanmu Z, Kocarek TA, Runge-Morris M. Transcriptional regulation of rat hepatic aryl sulfotransferase (SULT1A1) gene expression by glucocorticoids. Drug Metab Dispos. 2001;29:1130–1135.

25. Kurogi K, Chepak A, Hanrahan MT, Liu M-Y, Sakakibara Y, Suiko M, et al. Sulfation of opioid drugs by human cytosolic sulfotransferases: metabolic labeling study and enzymatic analysis. Eur J Pharm Sci. 2014;62:40–48.

26. Maiti S, Chen G. Ethanol up-regulates phenol sulfotransferase (SULT1A1) and hydroxysteroid sulfotransferase (SULT2A1) in rat liver and intestine. Arch Physiol Biochem. 2015;121:68–74.

27. Liu M, Fan R, Liu X, Cheng F, Wang J. Pathways and networks-based analysis of candidate genes associated with nicotine addiction. PLoS One. 2015;10:e0127438.

28. Dennen CA, Blum K, Bowirrat A, Khalsa J, Thanos PK, Baron D, et al. Neurogenetic and Epigenetic Aspects of Cannabinoids. Epigenomes. 2022;6:27.

29. Jiang Y, Wei D, Xie Y. Functional modular networks identify the pivotal genes associated with morphine addiction and potential drug therapies. BMC Anesthesiol. 2023;23:151.

30. Korostynski M, Piechota M, Kaminska D, Solecki W, Przewlocki R. Morphine effects on striatal transcriptome in mice. Genome Biology. 2007;8:R128.

31. Piechota M, Korostynski M, Sikora M, Golda S, Dzbek J, Przewlocki R. Common transcriptional effects in the mouse striatum following chronic treatment with heroin and methamphetamine. Genes, Brain, and Behavior. 2012;11:404–414.

32. Skupio U, Sikora M, Korostynski M, Wawrzczak-Bargiela A, Piechota M, Ficek J, et al. Behavioral and transcriptional patterns of protracted opioid self-administration in mice. Addiction Biology. 2017;22:1802–1816.

33. Tieskens JM, van Lier PAC, Buil JM, Barker ED. Sensation-seeking-related DNA methylation and the development of delinquency: A longitudinal epigenome-wide study. Dev Psychopathol. 2023;35:791–799.

### Supplementary Figure 1

A

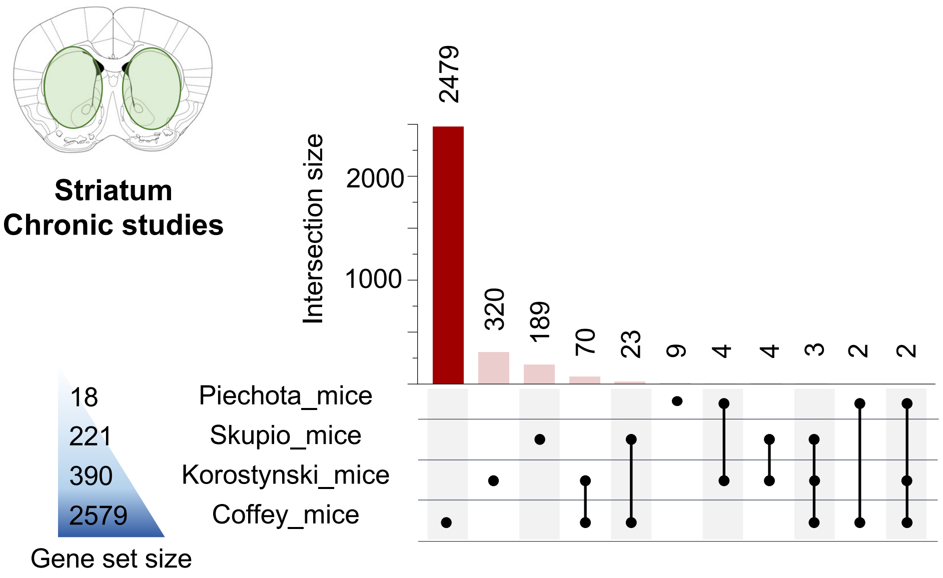


B

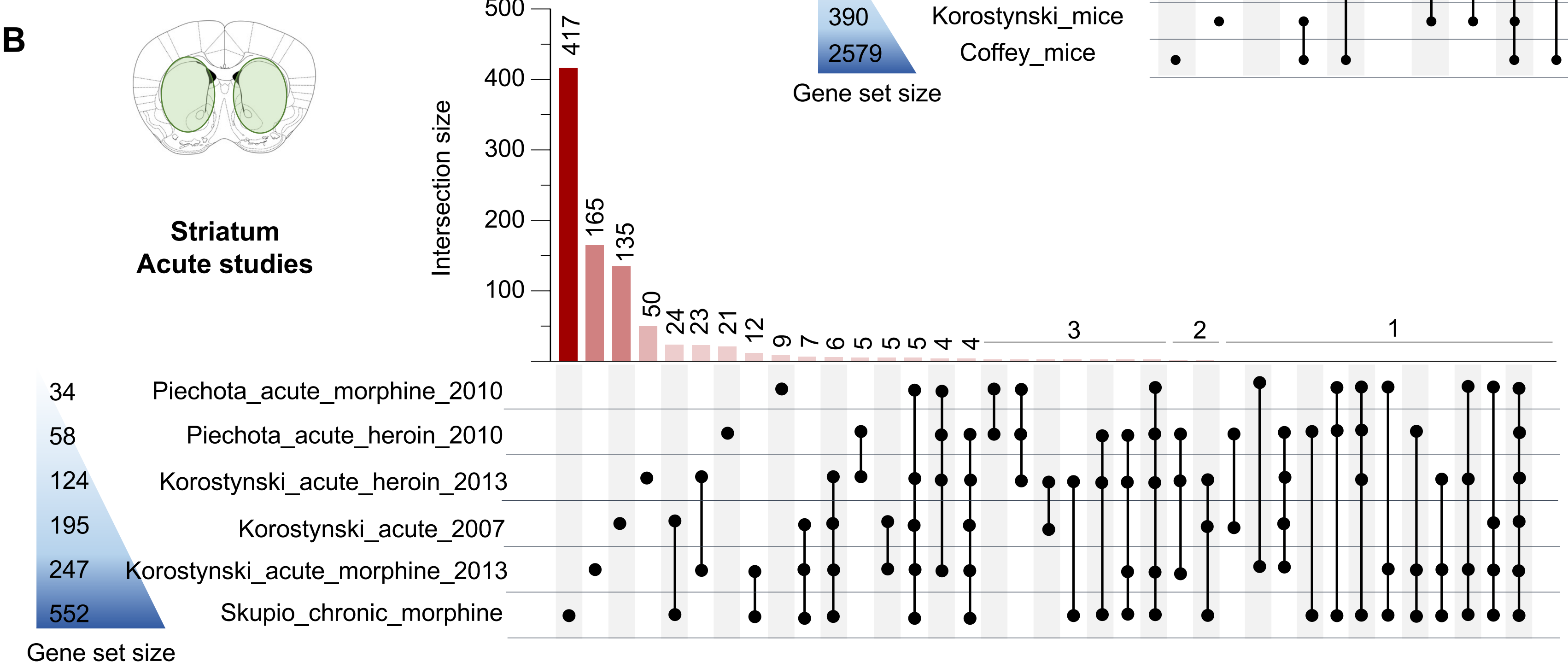

**Supplementary Figure 2**

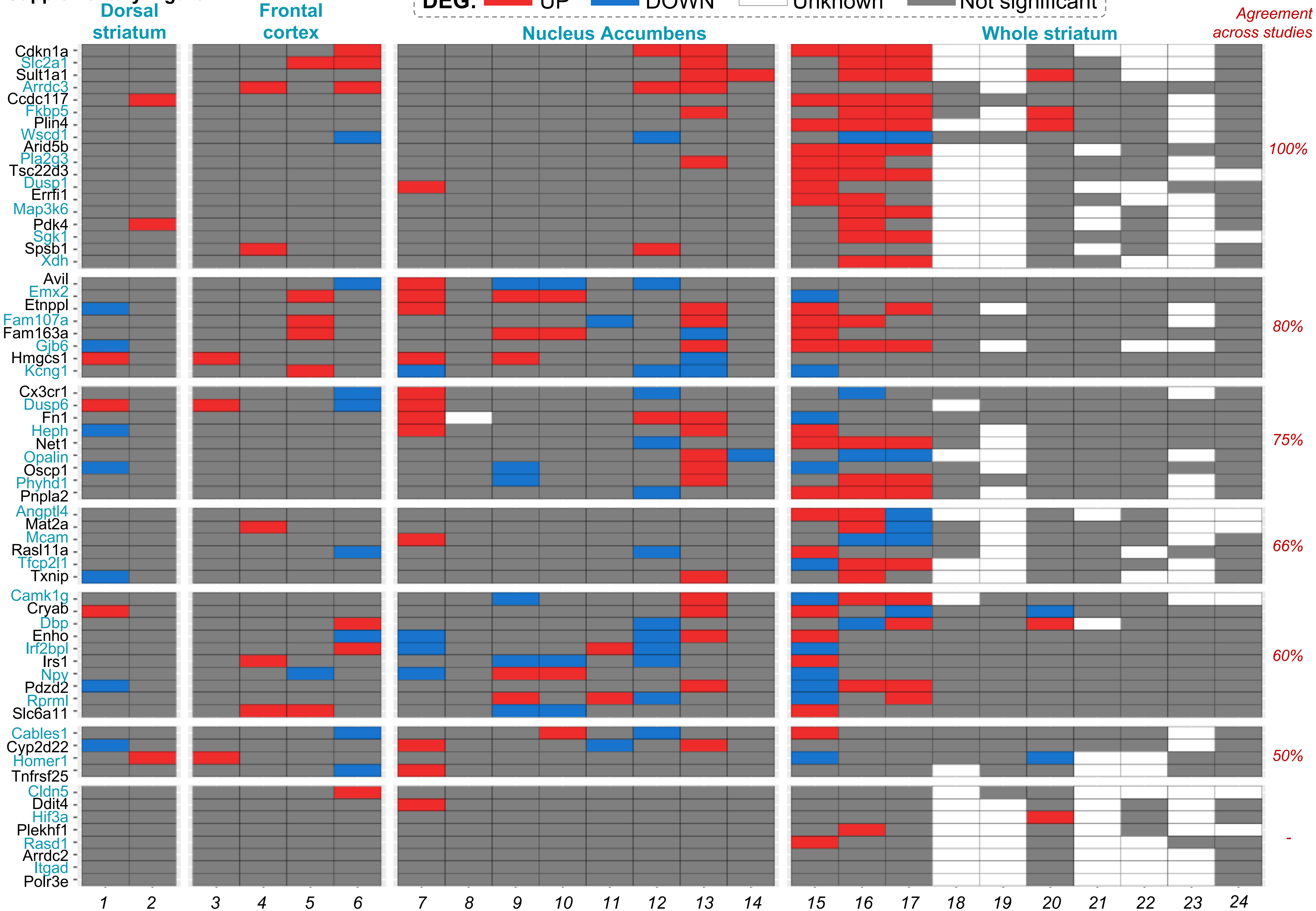

**Supplementary Figure 3**

**A**

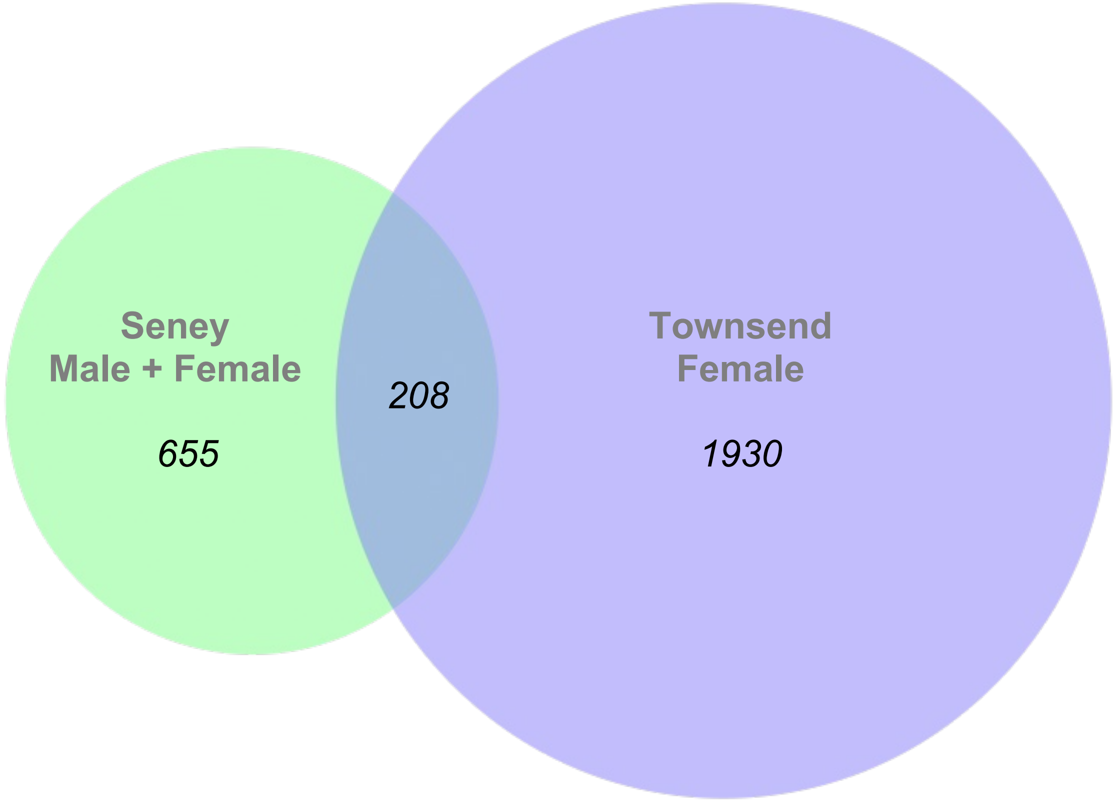


**B**

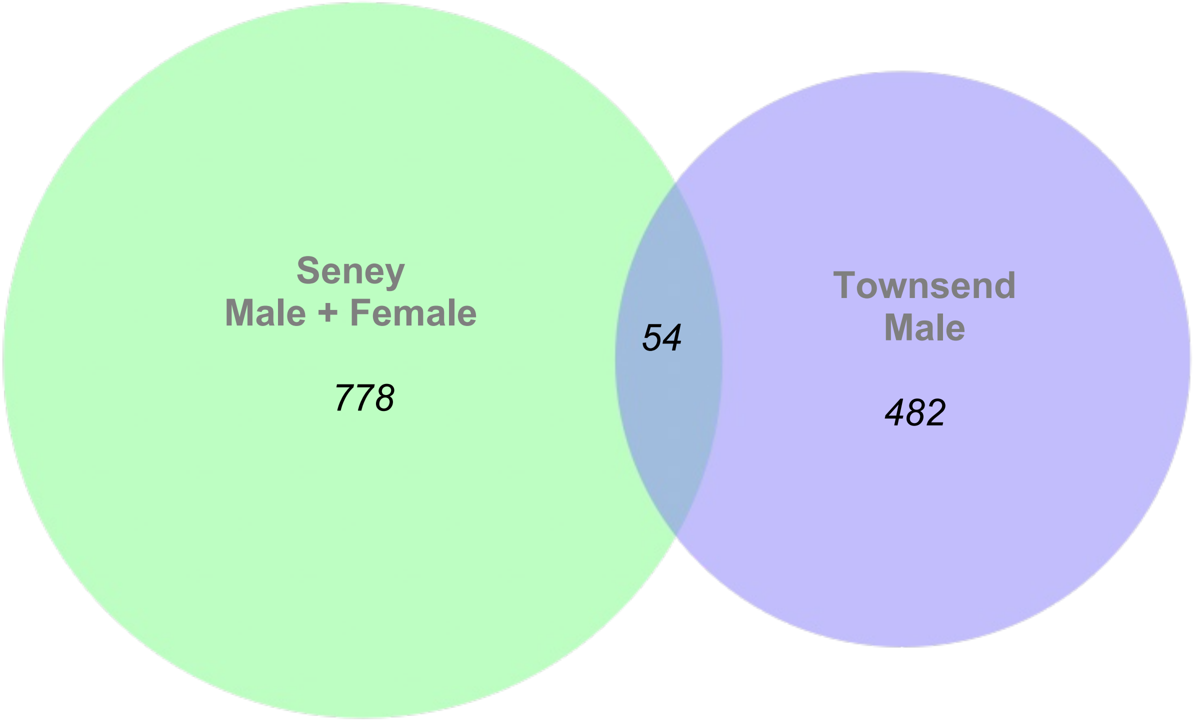

**Supplementary Figure 4**

A

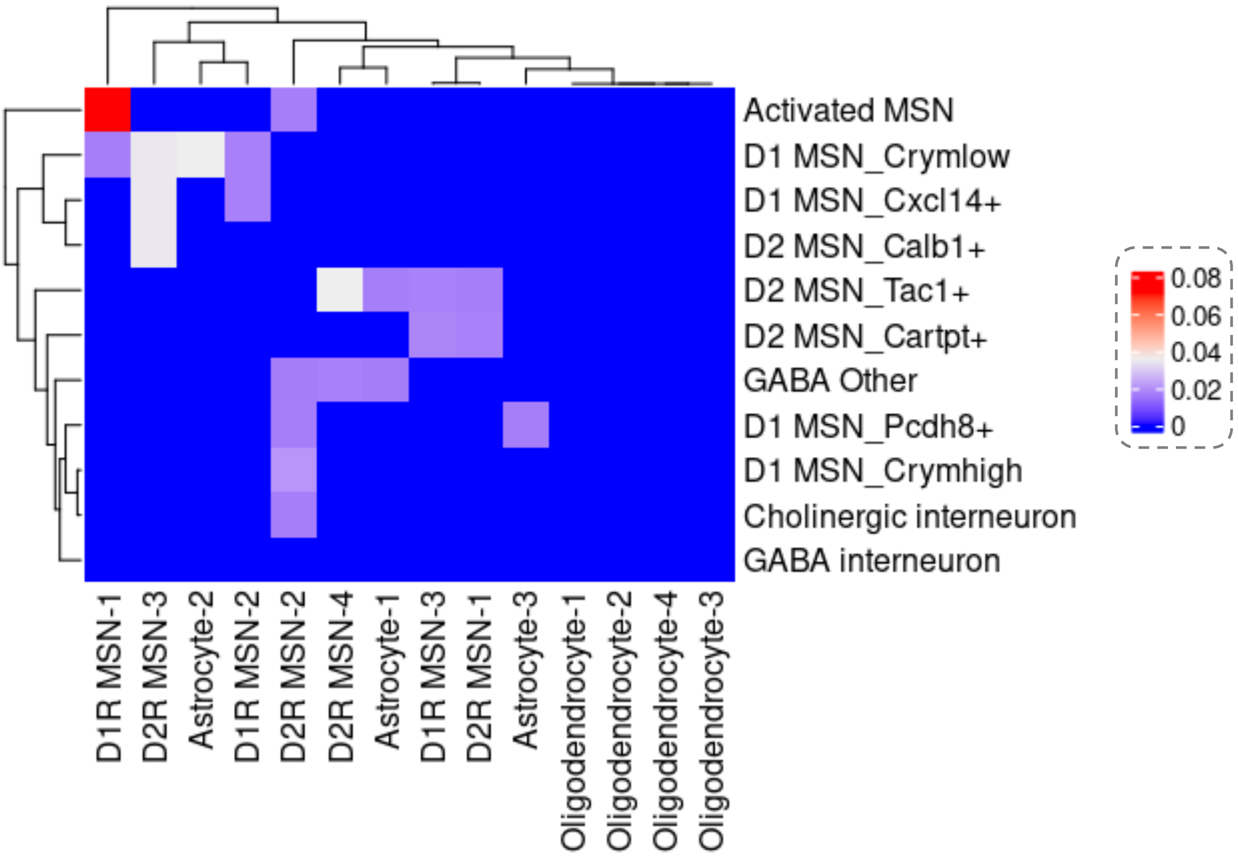


B

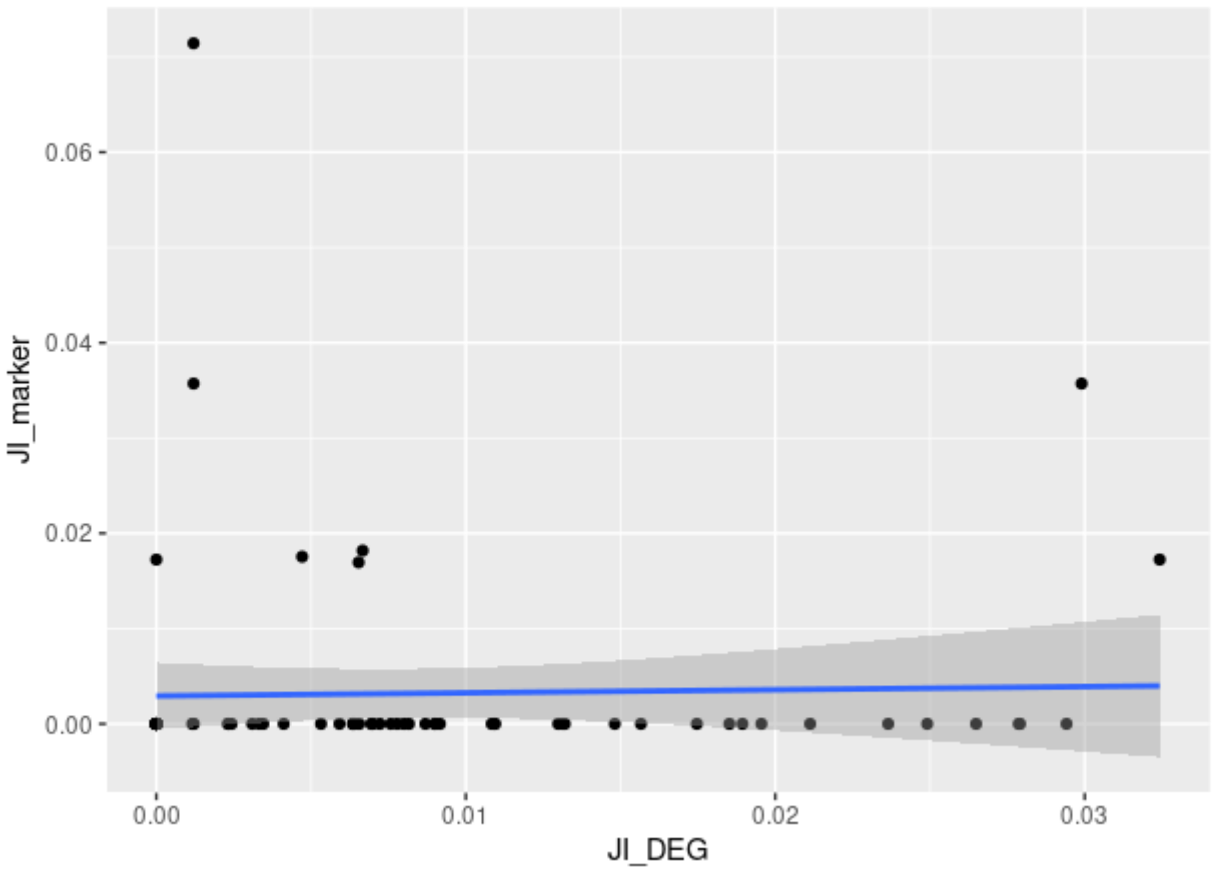
