## Supplementary material for "Functional genomic mechanisms of opioid action and opioid use disorder: a systematic review of animal models and human studies": Code and data: Fig4.R

```
# Code to reproduce Figure 4.B-F
# Alba Caparros-Roissard

library(readxl)
library(UpSetR)
library(dplyr)
library(biomaRt)

##########################
# Figure 4C - Human, Nac #
##########################

# 2 studies: Albertson et al and Seney et al

Albertson_human <- read_excel("DEG_Nac.xlsx", sheet = "Albertson_human", skip = 2)
# significant probes are already selected, chips AFFYMETRIX HumanGene U133A and AFFYMETRIX HumanGene U133B were used (need to translate probes to genes)
mart <- useMart("ENSEMBL_MART_ENSEMBL")
mart <- useDataset("hsapiens_gene_ensembl", mart)
Albertson_human_U133A <- (getBM(attributes = c("affy_hg_u133a", "external_gene_name"), filters = "affy_hg_u133a", 
                                values = Albertson_human$...1, mart = mart))$external_gene_name
Albertson_human_U133B <- (getBM(attributes = c("affy_hg_u133b", "external_gene_name"), filters = "affy_hg_u133b", 
                                values = Albertson_human$...1, mart = mart))$external_gene_name
# discard duplicates (duplicates of genes that were found by both chips, or by several probes in one chip so that each gene only appears once)
Albertson_human <- unique(c(Albertson_human_U133A, Albertson_human_U133B)); rm(Albertson_human_U133A, Albertson_human_U133B)

Seney_human <- read_excel("DEG_Nac.xlsx", sheet = "Seney_human")
# in their data file they mention their cutoffs: corrected p value < 0.01 and absolute log2 FC > 1.3 
Seney_human <- (Seney_human %>% filter(corrected.p < 0.01 & abs(coefficient) > 0.26))$gene_symbol

upset(fromList(list(Albertson_human = Albertson_human, Seney_human = Seney_human)), nsets = 2, order.by = "freq", nintersects = NA, keep.order = TRUE)
# We then used https://www.biovenn.nl/ to make the Venn diagram using both lists of genes

#########################
# Figure 4D - Mice, Nac #
#########################

# 3 studies: Lefevre et al, Grice et al and Ye et al

Lefevre_mice <- read_excel("DEG_Nac.xlsx", sheet = "Lefevre_mice", skip = 1)
# pvalue is at < 0.05 already
Lefevre_mice <- Lefevre_mice$GeneName

Grice_mice <- read_excel("DEG_Nac.xlsx", sheet = "Grice_mice", skip = 1)
# pvalue is at < 0.01 already, chip AFFYMETRIX MouseGene U74A was used (need to translate probes to genes)
mart <- useMart("ENSEMBL_MART_ENSEMBL")
mart <- useDataset("mmusculus_gene_ensembl", mart)
Grice_mice <- unique((getBM(attributes = c("affy_mg_u74a", "external_gene_name"), filters = "affy_mg_u74a", 
                            values = Grice_mice$`probe set`, mart = mart))$external_gene_name)

Ye_mice <- read_excel("DEG_Nac.xlsx", sheet = "Ye_mice", skip = 0)
# significant genes already selected
Ye_mice <- Ye_mice$gene_id

upset(fromList(list(Grice_mice = Grice_mice, Lefevre_mice = Lefevre_mice, Ye_mice = Ye_mice)), nsets = 3, order.by = "freq", nintersects = NA, keep.order = TRUE)
# We then used https://www.biovenn.nl/ to make the Venn diagram using the three lists of genes

########################
# Figure 4E - Rat, Nac #
########################

# 3 studies: Imperio et al (Small Suppressor and Large Suppressor) and Townsend et al

Imperio_LS_rat <- read_excel("DEG_Nac.xlsx", sheet = "Imperio_rat")
# discard genes where FC == NA (not significant)
Imperio_LS_rat <- (Imperio_LS_rat %>% filter(!is.na(`FC ([Large Suppressor] vs [Saline]) ( Sal vs Lrg Supp Fold change >= 1.25 )`)))$`Gene Symbol`

Imperio_SS_rat <- read_excel("DEG_Nac.xlsx", sheet = "Imperio_rat")
# discard genes where FC == NA (not significant)
Imperio_SS_rat <- (Imperio_SS_rat %>% filter(!is.na(`FC ([Small Suppressor] vs [Saline]) (Small Supp vs Saline Fold change >= 1.25 )`)))$`Gene Symbol`

Townsend_rat <- read_excel("DEG_Nac.xlsx", sheet = "Townsend_rat_males", skip = 1)
# in their figure they mention their cutoffs: raw p value < 0.05 and absolute log2 FC > 1.3
# we discarded the fold-change cutoff, as otherwise we only had 9 DEG, which would have hindered the DEG list comparisons
# length((Townsend_rat %>% filter(pvalue < 0.05 & abs(log2FoldChange) > 1.3))$Gene.name) # 9
Townsend_rat <- (Townsend_rat %>% filter(pvalue < 0.05))$Gene.name

upset(fromList(list(Imperio_LS_rat = Imperio_LS_rat, Imperio_SS_rat = Imperio_SS_rat, Townsend_rat = Townsend_rat)), nsets = 3, order.by = "freq", 
      nintersects = NA, keep.order = TRUE)
# We then used https://www.biovenn.nl/ to make the Venn diagram using the three lists of genes

################################
# Figure 4B - all species, Nac #
################################

# We are choosing mice as a "reference" (thus need to find ortholog genes for rat > mice and human > mice; no need to change mice gene lists)
mart <- useMart("ENSEMBL_MART_ENSEMBL")
mart <- useDataset("rnorvegicus_gene_ensembl", mart)
Imperio_LS_rat_tr_mice <- unique((getBM(attributes = c("mmusculus_homolog_associated_gene_name", "external_gene_name"), 
                                        filters = c("with_mmusculus_homolog", "external_gene_name"),
                                        values = list(TRUE, Imperio_LS_rat), mart = mart))$mmusculus_homolog_associated_gene_name)
Imperio_SS_rat_tr_mice <- unique((getBM(attributes = c("mmusculus_homolog_associated_gene_name", "external_gene_name"), 
                                        filters = c("with_mmusculus_homolog", "external_gene_name"),
                                        values = list(TRUE, Imperio_SS_rat), mart = mart))$mmusculus_homolog_associated_gene_name)
Townsend_rat_tr_mice <- unique((getBM(attributes = c("mmusculus_homolog_associated_gene_name", "external_gene_name"), 
                                           filters = c("with_mmusculus_homolog", "external_gene_name"),
                                           values = list(TRUE, Townsend_rat), mart = mart))$mmusculus_homolog_associated_gene_name)
mart <- useMart("ENSEMBL_MART_ENSEMBL")
mart <- useDataset("hsapiens_gene_ensembl", mart)
Albertson_human_tr_mice <- unique((getBM(attributes = c("mmusculus_homolog_associated_gene_name", "external_gene_name"), 
                                         filters = c("with_mmusculus_homolog", "external_gene_name"),
                                         values = list(TRUE, Albertson_human), mart = mart))$mmusculus_homolog_associated_gene_name)
Seney_human_tr_mice <- unique((getBM(attributes = c("mmusculus_homolog_associated_gene_name", "external_gene_name"), 
                                     filters = c("with_mmusculus_homolog", "external_gene_name"),
                                     values = list(TRUE, Seney_human), mart = mart))$mmusculus_homolog_associated_gene_name)
library(UpSetR)
upset(fromList(list(Grice_mice = Grice_mice, Lefevre_mice = Lefevre_mice, Ye_mice = Ye_mice,
                    Imperio_LS_rat = Imperio_LS_rat_tr_mice, Imperio_SS_rat = Imperio_SS_rat_tr_mice, Townsend_rat = Townsend_rat_tr_mice,
                    Albertson_human = Albertson_human_tr_mice, Seney_human = Seney_human_tr_mice)), nsets = 8, order.by = "freq", nintersects = 15, keep.order = TRUE)

save(Albertson_human_tr_mice, Grice_mice, Imperio_LS_rat_tr_mice, Imperio_SS_rat_tr_mice, Lefevre_mice, Seney_human_tr_mice, Townsend_rat_tr_mice, Ye_mice, 
     file = "genes_DE_all_NAc.RData")

###############################
# Figure 4F - Dorsal striatum #
###############################

# 2 studies: Lefevre et al and Valderrama et al

Lefevre_mice <- read_excel("DEG_dorsal_striatum.xlsx", sheet = "Lefevre_mice", skip = 1)
Valderrama_rats <- read_excel("DEG_dorsal_striatum.xlsx", sheet = "Valderrama_rat", skip = 3, 
                              col_names = c("Probeset ID", "Gene Symbol", "Morphine vs control 1h", "Morphine vs control 2h"))
mart <- useMart("ENSEMBL_MART_ENSEMBL")
mart <- useDataset("rnorvegicus_gene_ensembl", mart)
Valderrama_rats_genes <- Valderrama_rats$`Gene Symbol`[!Valderrama_rats$`Gene Symbol` == "---"]
Valderrama_rats_genes[1] <- "Hspa1b"
# we only re-annotate probes for which no corresponding genes were provided in their study and keep their other annotations
Valderrama_rats_conv <- (getBM(attributes = c("affy_rat230_2", "external_gene_name"), filters = "affy_rat230_2", 
                               values = Valderrama_rats[Valderrama_rats$`Gene Symbol` == "---",]$`Probeset ID`, mart = mart))
Valderrama_rats <- unique(c(Valderrama_rats_genes, Valderrama_rats_conv$external_gene_name))

Lefevre_mice <- Lefevre_mice$GeneName

# We are choosing mice as a "reference" (thus need to find ortholog genes for rat > mice; no need to change the mice gene list)
Valderrama_rats_tr_mice <- unique((getBM(attributes = c("mmusculus_homolog_associated_gene_name", "external_gene_name"), 
                                         filters = c("with_mmusculus_homolog", "external_gene_name"),
                                         values = list(TRUE, Valderrama_rats), mart = mart))$mmusculus_homolog_associated_gene_name)

upset(fromList(list(Lefevre_mice = Lefevre_mice, Valderrama_rats = Valderrama_rats_tr_mice)), nsets = 2, order.by = "freq", nintersects = NA, keep.order = TRUE)

save(Lefevre_mice, Valderrama_rats_tr_mice, file = "genes_DE_all_dorsal_striatum.RData")
```
