## Supplementary material for "Functional genomic mechanisms of opioid action and opioid use disorder: a systematic review of animal models and human studies": Code and data: Fig5.R

```
# Code to reproduce Figure 5.A-D and Supplementary Figure 1.A-B
# Alba Caparros-Roissard

library(readxl)
library(biomaRt)
library(UpSetR)
library(dplyr)
library(ggplot2)

######################################################
# Supplementary Figure 1A - Striatum (acute studies) #
######################################################

# 6 studies: Piechota et al 2010 (morphine and heroin), Korostynski et al 2007 (acute), Korostynski et al 2013 (morphine and heroin), Skupio et al (acute)

# We chose to pool time points for Piechota et al, 2010 and Korostynski et al, 2013
Piechota_mice_acute <- read_excel("DEG_striatum.xlsx", sheet = "Piechota_acute_2010")
Piechota_mice_acute_morphine <- Piechota_mice_acute[Piechota_mice_acute$`1h Morphine` < 0.05,]$`Gene symbol`
Piechota_mice_acute_morphine <- c(Piechota_mice_acute_morphine, Piechota_mice_acute[Piechota_mice_acute$`2h Morphine` < 0.05,]$`Gene symbol`)
Piechota_mice_acute_morphine <- c(Piechota_mice_acute_morphine, Piechota_mice_acute[Piechota_mice_acute$`4h Morphine` < 0.05,]$`Gene symbol`)
Piechota_mice_acute_morphine <- unique(c(Piechota_mice_acute_morphine, Piechota_mice_acute[Piechota_mice_acute$`8h Morphine` < 0.05,]$`Gene symbol`))

Piechota_mice_acute_heroin <- Piechota_mice_acute[Piechota_mice_acute$`1h Heroin` < 0.05,]$`Gene symbol`
Piechota_mice_acute_heroin <- c(Piechota_mice_acute_heroin, Piechota_mice_acute[Piechota_mice_acute$`2h Heroin` < 0.05,]$`Gene symbol`)
Piechota_mice_acute_heroin <- c(Piechota_mice_acute_heroin, Piechota_mice_acute[Piechota_mice_acute$`4h Heroin` < 0.05,]$`Gene symbol`)
Piechota_mice_acute_heroin <- unique(c(Piechota_mice_acute_heroin, Piechota_mice_acute[Piechota_mice_acute$`8h Heroin` < 0.05,]$`Gene symbol`))

# Korostynski et al, 2007: AFFYMETRIX GeneChip Mouse Genome 430 2.0 chip used 
Korostynski_mice_acute <- read_excel("DEG_striatum.xlsx", sheet = "Korostynski_acute_2007")
mart <- useMart("ENSEMBL_MART_ENSEMBL")
mart <- useDataset("mmusculus_gene_ensembl", mart)
Korostynski_mice_conv <- (getBM(attributes = c("affy_mouse430_2", "external_gene_name"), filters = "affy_mouse430_2", 
                                values = Korostynski_mice_acute[!is.na(Korostynski_mice_acute$`Probe set...2`),]$`Probe set...2`, mart = mart))
Korostynski_mice_conv <- rbind(Korostynski_mice_conv, (getBM(attributes = c("affy_mouse430_2", "external_gene_name"), filters = "affy_mouse430_2", 
                                                             values = Korostynski_mice_acute[!is.na(Korostynski_mice_acute$`Probe set...6`),]$`Probe set...6`, mart = mart)))
Korostynski_mice_acute <- unique(Korostynski_mice_conv$external_gene_name)

Korostynski_2013 <- read_excel("DEG_striatum.xlsx", sheet = "Korostynski_acute_2013")
Korostynski_mice_acute_morphine <- na.omit(Korostynski_2013$`1h Morphine`)
Korostynski_mice_acute_morphine <- c(Korostynski_mice_acute_morphine, na.omit(Korostynski_2013$`2h Morphine`))
Korostynski_mice_acute_morphine <- c(Korostynski_mice_acute_morphine, na.omit(Korostynski_2013$`4h Morphine`))
Korostynski_mice_acute_morphine <- unique(c(Korostynski_mice_acute_morphine, na.omit(Korostynski_2013$`8h Morphine`)))
Korostynski_mice_conv <- getBM(attributes = c("ensembl_gene_id", "external_gene_name"), filters = "ensembl_gene_id", 
                               values = Korostynski_mice_acute_morphine, mart = mart)
Korostynski_mice_acute_morphine <- unique(Korostynski_mice_conv$external_gene_name)

Korostynski_mice_acute_heroin <- na.omit(Korostynski_2013$`1h Heroin`)
Korostynski_mice_acute_heroin <- c(Korostynski_mice_acute_heroin, na.omit(Korostynski_2013$`2h Heroin`))
Korostynski_mice_acute_heroin <- c(Korostynski_mice_acute_heroin, na.omit(Korostynski_2013$`4h Heroin`))
Korostynski_mice_acute_heroin <- unique(c(Korostynski_mice_acute_heroin, na.omit(Korostynski_2013$`8h Heroin`)))
Korostynski_mice_conv <- getBM(attributes = c("ensembl_gene_id", "external_gene_name"), filters = "ensembl_gene_id", 
                               values = Korostynski_mice_acute_heroin, mart = mart)
Korostynski_mice_acute_heroin <- unique(Korostynski_mice_conv$external_gene_name); rm(Korostynski_2013)

Skupio_mice_acute <- read_excel("DEG_striatum.xlsx", sheet = "Skupio_acute")
Skupio_mice_acute <- unique(Skupio_mice_acute$Gene)

upset(fromList(list(Korostynski_mice_acute_2007 = Korostynski_mice_acute, Korostynski_mice_acute_heroin_2013 = Korostynski_mice_acute_heroin, 
                    Korostynski_mice_acute_morphine_2013 = Korostynski_mice_acute_morphine, Piechota_mice_acute_heroin = Piechota_mice_acute_heroin, 
                    Piechota_mice_acute_morphine = Piechota_mice_acute_morphine, Skupio_mice_acute = Skupio_mice_acute)), 
      nsets = 6, order.by = "freq", nintersects = 16, keep.order = TRUE)

########################################################
# Supplementary Figure 1B - Striatum (chronic studies) #
########################################################

# 4 studies: Korostynski et al 2007 (chronic), Coffey et al (chronic), Skupio et al (chronic), Piechota et al 2012

# Korostynski et al, 2007: AFFYMETRIX GeneChip Mouse Genome 430 2.0 chip used 
Korostynski_mice_chronic <- read_excel("DEG_striatum.xlsx", sheet = "Korostynski_chronic_2007")
mart <- useMart("ENSEMBL_MART_ENSEMBL")
mart <- useDataset("mmusculus_gene_ensembl", mart)
Korostynski_mice_conv <- (getBM(attributes = c("affy_mouse430_2", "external_gene_name"), filters = "affy_mouse430_2", 
                                values = Korostynski_mice_chronic[!is.na(Korostynski_mice_chronic$`Probe set...2`),]$`Probe set...2`, mart = mart))
Korostynski_mice_conv <- rbind(Korostynski_mice_conv, (getBM(attributes = c("affy_mouse430_2", "external_gene_name"), filters = "affy_mouse430_2", 
                                                             values = Korostynski_mice_chronic[!is.na(Korostynski_mice_chronic$`Probe set...6`),]$`Probe set...6`, mart = mart)))
Korostynski_mice_chronic <- unique(Korostynski_mice_conv$external_gene_name)

Coffey_mice_chronic <- read_excel("DEG_striatum.xlsx", sheet = "Coffey")
Coffey_mice_chronic <- (na.omit(Coffey_mice_chronic)[na.omit(Coffey_mice_chronic$`P-adj` < 0.1),])$GeneID

Skupio_mice_chronic <- read_excel("DEG_striatum.xlsx", sheet = "Skupio_chronic")
Skupio_mice_chronic <- unique(Skupio_mice_chronic$Gene)

Piechota_mice_chronic <- read_excel("DEG_striatum.xlsx", sheet = "Piechota_chronic_2012", col_names = FALSE)
Piechota_mice_chronic <- unique(Piechota_mice_chronic$...1)

upset(fromList(list(Korostynski_mice_chronic = Korostynski_mice_chronic,Piechota_mice_chronic = Piechota_mice_chronic, 
                    Skupio_mice_chronic = Skupio_mice_chronic, Coffey_mice_chronic = Coffey_mice_chronic)), 
      nsets = 4, order.by = "freq", nintersects = NA, keep.order = TRUE)

######################################
# Figure 5A - Striatum (all studies) #
######################################

upset(fromList(list(Korostynski_mice_acute_2007 = Korostynski_mice_acute, Korostynski_mice_acute_heroin_2013 = Korostynski_mice_acute_heroin, 
                    Korostynski_mice_acute_morphine_2013 = Korostynski_mice_acute_morphine, Korostynski_mice_chronic_2007 = Korostynski_mice_chronic,
                    Piechota_mice_acute_heroin = Piechota_mice_acute_heroin, Piechota_mice_acute_morphine = Piechota_mice_acute_morphine, 
                    Piechota_mice_chronic = Piechota_mice_chronic, Skupio_mice_chronic = Skupio_mice_chronic, Skupio_mice_acute = Skupio_mice_acute, 
                    Coffey_mice_chronic = Coffey_mice_chronic)), 
      nsets = 10, order.by = "freq", nintersects = 20, keep.order = TRUE)

save(Korostynski_mice_acute, Korostynski_mice_acute_heroin, Korostynski_mice_acute_morphine, Korostynski_mice_chronic, 
     Piechota_mice_acute_heroin, Piechota_mice_acute_morphine, Piechota_mice_chronic, 
     Skupio_mice_acute, Skupio_mice_chronic, Coffey_mice_chronic, file = "genes_DE_all_striatum.RData")

##############################
# Figure 5B - Frontal cortex #
##############################

# 6 studies: Ammon et al 2003, Ammon-Treiber et al 2004, Seney et al, Mendez et al, Kuntz-Melcavage et al and Liu et al

Seney_human <- read_excel("DEG_frontal_cortex.xlsx", sheet = "Seney_human")
# Seney: cutoff- corrected p value < 0.01 and absolute log2 FC > 1.3
Seney_human <- (Seney_human %>% filter(corrected.p < 0.01 & abs(coefficient) > 0.26))$gene_symbol

Mendez_human <- read_excel("DEG_frontal_cortex.xlsx", sheet = "Mendez_human")
Mendez_human <- Mendez_human$gene[!is.na(Mendez_human$gene)]

# Ammon_rats_2003: Rat Genome U34 A Gene Chip (Affymetrix, Santa Clara, CA, USA)
Ammon_rats_2003 <- read_excel("DEG_frontal_cortex.xlsx", sheet = "Ammon_2003", skip = 1)
mart <- useMart("ENSEMBL_MART_ENSEMBL")
mart <- useDataset("rnorvegicus_gene_ensembl", mart)
Ammon_rats_2003 <- unique((getBM(attributes = c("affy_rg_u34a", "external_gene_name"), 
                                 filters = c("affy_rg_u34a", "external_gene_name"),
                                 values = list(TRUE, Ammon_rats_2003$`Gene bank acc. Number`), mart = mart))$external_gene_name)
# no probe identifiers manage to be linked to genes thanks to the chip name provided in their article > unusable

# Ammon_rats_2004: Rat Genome U34 B and C Gene Chip (Affymetrix, Santa Clara, CA, USA)
Ammon_rats_2004 <- read_excel("DEG_frontal_cortex.xlsx", sheet = "Ammon-Treiber_2004", skip = 1)
library(biomaRt)
mart <- useMart("ENSEMBL_MART_ENSEMBL")
mart <- useDataset("rnorvegicus_gene_ensembl", mart)
Ammon_rats_2004_affy_rg_u34b <- unique((getBM(attributes = c("affy_rg_u34b", "external_gene_name"), 
                                              filters = c("affy_rg_u34b", "external_gene_name"),
                                              values = list(TRUE, Ammon_rats_2004$`Gene chip, probe set ID`), mart = mart))$external_gene_name)
Ammon_rats_2004_affy_rg_u34c <- unique((getBM(attributes = c("affy_rg_u34c", "external_gene_name"), 
                                              filters = c("affy_rg_u34c", "external_gene_name"),
                                              values = list(TRUE, Ammon_rats_2004$`Gene chip, probe set ID`), mart = mart))$external_gene_name)
# no probe identifiers manage to be linked to genes thanks to either of the chip names provided in their article > unusable

Kuntz_Melcavage_rats <- read_excel("DEG_frontal_cortex.xlsx", sheet = "Kuntz-Melcavage_rat", skip = 1)
Kuntz_Melcavage_rats <- Kuntz_Melcavage_rats$Alias
Kuntz_Melcavage_rats <- Kuntz_Melcavage_rats[!is.na(Kuntz_Melcavage_rats)]

Liu_rats <- read_excel("DEG_frontal_cortex.xlsx", sheet = "Liu_rat")
Liu_rats <- Liu_rats$`Gene Symbol`

# We are choosing rat as a "reference" (thus need to find ortholog genes for human > rat; no need to change rat gene lists)
mart <- useMart("ENSEMBL_MART_ENSEMBL")
mart <- useDataset("hsapiens_gene_ensembl", mart)
Seney_human_tr_rats <- unique((getBM(attributes = c("rnorvegicus_homolog_associated_gene_name", "external_gene_name"), 
                                     filters = c("with_rnorvegicus_homolog", "external_gene_name"),
                                     values = list(TRUE, Seney_human), mart = mart))$rnorvegicus_homolog_associated_gene_name)
Mendez_human_tr_rats <- unique((getBM(attributes = c("rnorvegicus_homolog_associated_gene_name", "external_gene_name"), 
                                      filters = c("with_rnorvegicus_homolog", "external_gene_name"),
                                      values = list(TRUE, Mendez_human), mart = mart))$rnorvegicus_homolog_associated_gene_name)

upset(fromList(list(Kuntz_Melcavage_rats = Kuntz_Melcavage_rats, Liu_rats = Liu_rats, Mendez_human = Mendez_human_tr_rats, Seney_human = Seney_human_tr_rats)), 
      nsets = 4, order.by = "freq", nintersects = NA, keep.order = TRUE)

save(Kuntz_Melcavage_rats, Mendez_human, Liu_rats, Seney_human, file = "genes_DE_all_frontal_cortex.RData")

#########################################
# Figure 5C - Histogram for all regions #
#########################################

load("genes_DE_all_dorsal_striatum.RData")
Lefevre_mice_dorsal_striatum <- Lefevre_mice; rm(Lefevre_mice)
Valderrama_rat_dorsal_striatum <- Valderrama_rats_tr_mice; rm(Valderrama_rats_tr_mice)

load("genes_DE_all_frontal_cortex.RData")
Kuntz_Melcavage_rat_frontal_cortex <- Kuntz_Melcavage_rats; rm(Kuntz_Melcavage_rats)
Liu_rat_frontal_cortex <- Liu_rats; rm(Liu_rats)
Seney_human_frontal_cortex <- Seney_human; rm(Seney_human)
Mendez_human_frontal_cortex <- Mendez_human; rm(Mendez_human)

load("genes_DE_all_NAc.RData")
Albertson_human_NAc <- Albertson_human_tr_mice; rm(Albertson_human_tr_mice)
Grice_mice_NAc <- Grice_mice; rm(Grice_mice)
Imperio_LS_rat_NAc <- Imperio_LS_rat_tr_mice; rm(Imperio_LS_rat_tr_mice)
Imperio_SS_rat_NAc <- Imperio_SS_rat_tr_mice; rm(Imperio_SS_rat_tr_mice)
Lefevre_mice_NAc <- Lefevre_mice; rm(Lefevre_mice)
Seney_human_NAc <- Seney_human_tr_mice; rm(Seney_human_tr_mice)
Townsend_rat_NAc <- Townsend_rat_tr_mice; rm(Townsend_rat_tr_mice)
Ye_mice_NAc <- Ye_mice; rm(Ye_mice)

load("genes_DE_all_striatum.RData")
Coffey_mice_chronic_striatum <- Coffey_mice_chronic; rm(Coffey_mice_chronic)
Korostynski_mice_acute_striatum <- Korostynski_mice_acute; rm(Korostynski_mice_acute)
Korostynski_mice_acute_heroin_striatum <- Korostynski_mice_acute_heroin; rm(Korostynski_mice_acute_heroin)
Korostynski_mice_acute_morphine_striatum <- Korostynski_mice_acute_morphine; rm(Korostynski_mice_acute_morphine)
Korostynski_mice_chronic_striatum <- Korostynski_mice_chronic; rm(Korostynski_mice_chronic)
Piechota_mice_acute_heroin_striatum <- Piechota_mice_acute_heroin; rm(Piechota_mice_acute_heroin)
Piechota_mice_acute_morphine_striatum <- Piechota_mice_acute_morphine; rm(Piechota_mice_acute_morphine)
Piechota_mice_chronic_striatum <- Piechota_mice_chronic; rm(Piechota_mice_chronic)
Skupio_mice_chronic_striatum <- Skupio_mice_chronic; rm(Skupio_mice_chronic)
Skupio_mice_acute_striatum <- Skupio_mice_acute; rm(Skupio_mice_acute)

mart <- useMart("ENSEMBL_MART_ENSEMBL")
mart <- useDataset("rnorvegicus_gene_ensembl", mart)
Kuntz_Melcavage_rat_frontal_cortex <- unique(getBM(attributes = c("mmusculus_homolog_associated_gene_name", "external_gene_name"), 
                                                   filters = c("with_mmusculus_homolog", "external_gene_name"),
                                                   values = list(TRUE, Kuntz_Melcavage_rat_frontal_cortex), mart = mart)$mmusculus_homolog_associated_gene_name)
Liu_rat_frontal_cortex <- unique(getBM(attributes = c("mmusculus_homolog_associated_gene_name", "external_gene_name"), 
                                       filters = c("with_mmusculus_homolog", "external_gene_name"),
                                       values = list(TRUE, Liu_rat_frontal_cortex), mart = mart)$mmusculus_homolog_associated_gene_name)
mart <- useMart("ENSEMBL_MART_ENSEMBL")
mart <- useDataset("hsapiens_gene_ensembl", mart)
Seney_human_frontal_cortex <- unique(getBM(attributes = c("mmusculus_homolog_associated_gene_name", "external_gene_name"), 
                                           filters = c("with_mmusculus_homolog", "external_gene_name"),
                                           values = list(TRUE, Seney_human_frontal_cortex), mart = mart)$mmusculus_homolog_associated_gene_name)
Mendez_human_frontal_cortex <- unique(getBM(attributes = c("mmusculus_homolog_associated_gene_name", "external_gene_name"), 
                                            filters = c("with_mmusculus_homolog", "external_gene_name"),
                                            values = list(TRUE, Mendez_human_frontal_cortex), mart = mart)$mmusculus_homolog_associated_gene_name)

comp_genes <- table(c(Albertson_human_NAc, Coffey_mice_chronic_striatum, Grice_mice_NAc, Imperio_LS_rat_NAc, Imperio_SS_rat_NAc, Korostynski_mice_acute_heroin_striatum, 
                      Korostynski_mice_acute_morphine_striatum, Korostynski_mice_acute_striatum, Korostynski_mice_chronic_striatum, Kuntz_Melcavage_rat_frontal_cortex, 
                      Lefevre_mice_dorsal_striatum, Lefevre_mice_NAc, Liu_rat_frontal_cortex, Mendez_human_frontal_cortex, Piechota_mice_acute_heroin_striatum, 
                      Piechota_mice_acute_morphine_striatum, Piechota_mice_chronic_striatum, Seney_human_frontal_cortex, Seney_human_NAc, Skupio_mice_acute_striatum, 
                      Skupio_mice_chronic_striatum, Townsend_rat_NAc, Valderrama_rat_dorsal_striatum, Ye_mice_NAc))[order(
                        table(c(Albertson_human_NAc, Coffey_mice_chronic_striatum, Grice_mice_NAc, Imperio_LS_rat_NAc, Imperio_SS_rat_NAc, 
                                Korostynski_mice_acute_heroin_striatum, Korostynski_mice_acute_morphine_striatum, Korostynski_mice_acute_striatum, 
                                Korostynski_mice_chronic_striatum, Kuntz_Melcavage_rat_frontal_cortex, Lefevre_mice_dorsal_striatum, Lefevre_mice_NAc, 
                                Liu_rat_frontal_cortex, Mendez_human_frontal_cortex, Piechota_mice_acute_heroin_striatum, Piechota_mice_acute_morphine_striatum, 
                                Piechota_mice_chronic_striatum, Seney_human_frontal_cortex, Seney_human_NAc, Skupio_mice_acute_striatum, Skupio_mice_chronic_striatum,
                                Townsend_rat_NAc, Valderrama_rat_dorsal_striatum, Ye_mice_NAc)), 
                        decreasing = TRUE)]

data_frame(n_studies = as.vector(comp_genes)) %>%
  ggplot(aes(x = n_studies)) + geom_histogram(binwidth = 1) +
  stat_bin(binwidth= 1, geom="text", aes(label=..count..), vjust = -1) + scale_x_continuous(breaks = c(1:11), limits = c(0.5,11)) +
  ylab("Number of genes") + xlab("Number of studies a gene was found as DE by")

##########################################
# Figure 5D - Jaccard indexes comparison #
##########################################

rm(mart, comp_genes)
# For this step, only the 24 lists of genes should be loaded, remove any other objects from the environment using the rm() command

# get all jaccard indexes
jaccard_indexes <- matrix(rep(NA, length(ls())*length(ls())), ncol = length(ls()), dimnames = list(ls(), ls()))
for (i in rownames(jaccard_indexes)) {
  for (j in colnames(jaccard_indexes)) {
    common <- sum(get(i) %in% get(j))
    jaccard_indexes[i,j] <- common/(length(unique(c(get(i), get(j)))))
  }
}

# characteristics of the studies used for grouping purposes in this step are described in the "study_characteristics.ods" file

# jaccard index inter vs intraspecies
mice_studies <- c("Lefevre_mice_dorsal_striatum", "Grice_mice_NAc", "Lefevre_mice_NAc", "Ye_mice_NAc", "Coffey_mice_chronic_striatum", 
                  "Korostynski_mice_acute_heroin_striatum", "Korostynski_mice_acute_morphine_striatum", "Korostynski_mice_acute_striatum", 
                  "Korostynski_mice_chronic_striatum", "Piechota_mice_acute_heroin_striatum", "Piechota_mice_acute_morphine_striatum", "Piechota_mice_chronic_striatum", 
                  "Skupio_mice_acute_striatum", "Skupio_mice_chronic_striatum")
rat_studies <- c("Valderrama_rat_dorsal_striatum", "Kuntz_Melcavage_rat_frontal_cortex", "Liu_rat_frontal_cortex", "Imperio_LS_rat_NAc", 
                 "Imperio_SS_rat_NAc", "Townsend_rat_NAc")
human_studies <- c("Seney_human_frontal_cortex", "Mendez_human_frontal_cortex", "Albertson_human_NAc", "Seney_human_NAc")

i <- rep(human_studies, seq(from = (length(human_studies)-1), to = 0, by = -1)); j <- c()
for(n in 2:length(human_studies)) {j <- c(j, human_studies[n:length(human_studies)])}
i <- c(i, rep(rat_studies, seq(from = (length(rat_studies)-1), to = 0, by = -1)))
for(n in 2:length(rat_studies)) {j <- c(j, rat_studies[n:length(rat_studies)])}
i <- c(i, rep(mice_studies, seq(from = (length(mice_studies)-1), to = 0, by = -1)))
for(n in 2:length(mice_studies)) {j <- c(j, mice_studies[n:length(mice_studies)])}
JI_intra <- c()
for(n in 1:length(i)) {JI_intra <- c(JI_intra, (jaccard_indexes[i[n], j[n]])*100)}

i <- rep(row.names(jaccard_indexes), seq(from = (length(row.names(jaccard_indexes))-1), to = 0, by = -1)); j <- c()
for(n in 2:length(row.names(jaccard_indexes))) {j <- c(j, row.names(jaccard_indexes)[n:length(row.names(jaccard_indexes))])}
JI_all <- c()
for(n in 1:length(i)) {JI_all <- c(JI_all, (jaccard_indexes[i[n], j[n]])*100)}
JI_inter <- JI_all
for (n in 1:length(JI_intra)) {JI_inter <- JI_inter[-which.max(JI_inter == JI_intra[n])]}

data.frame(jaccard_indexes = c(JI_intra, JI_inter), type = c(rep("intraspecies", length(JI_intra)), rep("interspecies", length(JI_inter)))) %>%
  ggplot(aes(x = type, y = jaccard_indexes)) + geom_boxplot()
t.test(x = JI_intra, y = JI_inter) # significant: p = 0.0005924

# jaccard index inter vs intrastructures
dorsal_striatum_studies <- c("Lefevre_mice_dorsal_striatum", "Valderrama_rat_dorsal_striatum")
frontal_cortex_studies <- c("Kuntz_Melcavage_rat_frontal_cortex", "Liu_rat_frontal_cortex", "Seney_human_frontal_cortex", 
                            "Mendez_human_frontal_cortex")
NAc_studies <- c("Albertson_human_NAc", "Grice_mice_NAc", "Imperio_LS_rat_NAc", "Imperio_SS_rat_NAc", "Lefevre_mice_NAc", 
                 "Seney_human_NAc", "Townsend_rat_NAc", "Ye_mice_NAc")
striatum_studies <- c("Coffey_mice_chronic_striatum", "Korostynski_mice_acute_heroin_striatum", "Korostynski_mice_acute_morphine_striatum", 
                      "Korostynski_mice_acute_striatum", "Korostynski_mice_chronic_striatum", "Piechota_mice_acute_heroin_striatum", 
                      "Piechota_mice_acute_morphine_striatum", "Piechota_mice_chronic_striatum", "Skupio_mice_acute_striatum", "Skupio_mice_chronic_striatum")

i <- rep(dorsal_striatum_studies, seq(from = (length(dorsal_striatum_studies)-1), to = 0, by = -1)); j <- c()
for(n in 2:length(dorsal_striatum_studies)) {j <- c(j, dorsal_striatum_studies[n:length(dorsal_striatum_studies)])}
i <- c(i, rep(frontal_cortex_studies, seq(from = (length(frontal_cortex_studies)-1), to = 0, by = -1)))
for(n in 2:length(frontal_cortex_studies)) {j <- c(j, frontal_cortex_studies[n:length(frontal_cortex_studies)])}
i <- c(i, rep(NAc_studies, seq(from = (length(NAc_studies)-1), to = 0, by = -1)))
for(n in 2:length(NAc_studies)) {j <- c(j, NAc_studies[n:length(NAc_studies)])}
i <- c(i, rep(striatum_studies, seq(from = (length(striatum_studies)-1), to = 0, by = -1)))
for(n in 2:length(striatum_studies)) {j <- c(j, striatum_studies[n:length(striatum_studies)])}
JI_intra <- c()
for(n in 1:length(i)) {JI_intra <- c(JI_intra, (jaccard_indexes[i[n], j[n]])*100)}

i <- rep(row.names(jaccard_indexes), seq(from = (length(row.names(jaccard_indexes))-1), to = 0, by = -1)); j <- c()
for(n in 2:length(row.names(jaccard_indexes))) {j <- c(j, row.names(jaccard_indexes)[n:length(row.names(jaccard_indexes))])}
JI_all <- c()
for(n in 1:length(i)) {JI_all <- c(JI_all, (jaccard_indexes[i[n], j[n]])*100)}
JI_inter <- JI_all
for (n in 1:length(JI_intra)) {JI_inter <- JI_inter[-which.max(JI_inter == JI_intra[n])]}

data.frame(jaccard_indexes = c(JI_intra, JI_inter), type = c(rep("intraregions", length(JI_intra)), rep("interregions", length(JI_inter)))) %>%
  ggplot(aes(x = type, y = jaccard_indexes)) + geom_boxplot()
t.test(x = JI_intra, y = JI_inter) # significant: p = 0.0002268

# jaccard index inter vs intrachronicity of treatment
chronic_studies <- c("Lefevre_mice_dorsal_striatum", "Kuntz_Melcavage_rat_frontal_cortex", "Liu_rat_frontal_cortex", 
                     "Seney_human_frontal_cortex", "Mendez_human_frontal_cortex", "Albertson_human_NAc", "Grice_mice_NAc", 
                     "Imperio_LS_rat_NAc", "Imperio_SS_rat_NAc", "Lefevre_mice_NAc", "Seney_human_NAc", "Townsend_rat_NAc", 
                     "Coffey_mice_chronic_striatum", "Korostynski_mice_chronic_striatum", "Piechota_mice_chronic_striatum", "Skupio_mice_chronic_striatum")
acute_studies <- c("Valderrama_rat_dorsal_striatum", "Ye_mice_NAc", "Korostynski_mice_acute_heroin_striatum", "Korostynski_mice_acute_morphine_striatum", 
                   "Korostynski_mice_acute_striatum", "Piechota_mice_acute_heroin_striatum", "Piechota_mice_acute_morphine_striatum", "Skupio_mice_acute_striatum")

i <- rep(chronic_studies, seq(from = (length(chronic_studies)-1), to = 0, by = -1)); j <- c()
for(n in 2:length(chronic_studies)) {j <- c(j, chronic_studies[n:length(chronic_studies)])}
i <- c(i, rep(acute_studies, seq(from = (length(acute_studies)-1), to = 0, by = -1)))
for(n in 2:length(acute_studies)) {j <- c(j, acute_studies[n:length(acute_studies)])}
JI_intra <- c()
for(n in 1:length(i)) {JI_intra <- c(JI_intra, (jaccard_indexes[i[n], j[n]])*100)}

i <- rep(row.names(jaccard_indexes), seq(from = (length(row.names(jaccard_indexes))-1), to = 0, by = -1)); j <- c()
for(n in 2:length(row.names(jaccard_indexes))) {j <- c(j, row.names(jaccard_indexes)[n:length(row.names(jaccard_indexes))])}
JI_all <- c()
for(n in 1:length(i)) {JI_all <- c(JI_all, (jaccard_indexes[i[n], j[n]])*100)}
JI_inter <- JI_all
for (n in 1:length(JI_intra)) {JI_inter <- JI_inter[-which.max(JI_inter == JI_intra[n])]}

data.frame(jaccard_indexes = c(JI_intra, JI_inter), type = c(rep("intratreatments", length(JI_intra)), rep("intertreatments", length(JI_inter)))) %>%
  ggplot(aes(x = type, y = jaccard_indexes)) + geom_boxplot()
t.test(x = JI_intra, y = JI_inter) # significant: p = 0.01079

# jaccard index inter vs intratype of opioid > discard human studies and Townsend
morphine_studies <- c("Lefevre_mice_dorsal_striatum", "Valderrama_rat_dorsal_striatum", "Liu_rat_frontal_cortex", "Grice_mice_NAc", "Lefevre_mice_NAc", 
                      "Ye_mice_NAc", "Coffey_mice_chronic_striatum", "Korostynski_mice_acute_morphine_striatum", "Korostynski_mice_acute_striatum", 
                      "Korostynski_mice_chronic_striatum", "Piechota_mice_acute_morphine_striatum", "Skupio_mice_acute_striatum", "Skupio_mice_chronic_striatum")
heroin_studies <- c("Kuntz_Melcavage_rat_frontal_cortex", "Imperio_LS_rat_NAc", "Imperio_SS_rat_NAc", "Korostynski_mice_acute_heroin_striatum", 
                    "Piechota_mice_acute_heroin_striatum", "Piechota_mice_chronic_striatum")

i <- rep(morphine_studies, seq(from = (length(morphine_studies)-1), to = 0, by = -1)); j <- c()
for(n in 2:length(morphine_studies)) {j <- c(j, morphine_studies[n:length(morphine_studies)])}
i <- c(i, rep(heroin_studies, seq(from = (length(heroin_studies)-1), to = 0, by = -1)))
for(n in 2:length(heroin_studies)) {j <- c(j, heroin_studies[n:length(heroin_studies)])}
JI_intra <- c()
for(n in 1:length(i)) {JI_intra <- c(JI_intra, (jaccard_indexes[i[n], j[n]])*100)}

not_discarded <- row.names(jaccard_indexes)[!(row.names(jaccard_indexes) %in% c("Seney_human_frontal_cortex", "Mendez_human_frontal_cortex", 
                                                                                "Albertson_human_NAc", "Seney_human_NAc", "Townsend_rat_NAc"))]
i <- rep(not_discarded, seq(from = (length(not_discarded)-1), to = 0, by = -1)); j <- c()
for(n in 2:length(not_discarded)) {j <- c(j, not_discarded[n:length(not_discarded)])}
JI_all <- c()
for(n in 1:length(i)) {JI_all <- c(JI_all, (jaccard_indexes[i[n], j[n]])*100)}
JI_inter <- JI_all
for (n in 1:length(JI_intra)) {JI_inter <- JI_inter[-which.max(JI_inter == JI_intra[n])]}

data.frame(jaccard_indexes = c(JI_intra, JI_inter), type = c(rep("intradrugs", length(JI_intra)), rep("interdrugs", length(JI_inter)))) %>%
  ggplot(aes(x = type, y = jaccard_indexes)) + geom_boxplot()
t.test(x = JI_intra, y = JI_inter) # NOT significant: p = 0.4357

# jaccard index inter vs intrastrain (only intragroup possible is C57BL6 mice)
C57BL6_studies <- c("Coffey_mice_chronic_striatum", "Korostynski_mice_acute_striatum", "Korostynski_mice_acute_heroin_striatum", 
                    "Korostynski_mice_acute_morphine_striatum", "Korostynski_mice_chronic_striatum", "Lefevre_mice_dorsal_striatum", 
                    "Lefevre_mice_NAc", "Piechota_mice_acute_heroin_striatum", "Piechota_mice_acute_morphine_striatum", 
                    "Piechota_mice_chronic_striatum", "Skupio_mice_acute_striatum", "Skupio_mice_chronic_striatum")

i <- rep(C57BL6_studies, seq(from = (length(C57BL6_studies)-1), to = 0, by = -1)); j <- c()
for(n in 2:length(C57BL6_studies)) {j <- c(j, C57BL6_studies[n:length(C57BL6_studies)])}
JI_intra <- c()
for(n in 1:length(i)) {JI_intra <- c(JI_intra, (jaccard_indexes[i[n], j[n]])*100)}

not_discarded <- row.names(jaccard_indexes)[!(row.names(jaccard_indexes) %in% c("Seney_human_frontal_cortex", "Mendez_human_frontal_cortex", 
                                                                                "Albertson_human_NAc", "Seney_human_NAc", "Townsend_rat_NAc", 
                                                                                "Imperio_LS_rat_NAc", "Imperio_SS_rat_NAc", 
                                                                                "Kuntz_Melcavage_rat_frontal_cortex", "Liu_rat_frontal_cortex",
                                                                                "Valderrama_rat_dorsal_striatum"))]
i <- rep(not_discarded, seq(from = (length(not_discarded)-1), to = 0, by = -1)); j <- c()
for(n in 2:length(not_discarded)) {j <- c(j, not_discarded[n:length(not_discarded)])}
JI_all <- c()
for(n in 1:length(i)) {JI_all <- c(JI_all, (jaccard_indexes[i[n], j[n]])*100)}
JI_inter <- JI_all
for (n in 1:length(JI_intra)) {JI_inter <- JI_inter[-which.max(JI_inter == JI_intra[n])]}

data.frame(jaccard_indexes = c(JI_intra, JI_inter), type = c(rep("intrastrains", length(JI_intra)), rep("interstrains", length(JI_inter)))) %>%
  ggplot(aes(x = type, y = jaccard_indexes)) + geom_boxplot()
t.test(x = JI_intra, y = JI_inter) # significant: p = 5.53e-05

# jaccard index inter vs intraarticle
i <- c("Lefevre_mice_dorsal_striatum", "Seney_human_frontal_cortex", "Imperio_LS_rat_NAc", "Korostynski_mice_acute_heroin_striatum", 
       "Korostynski_mice_acute_striatum", "Piechota_mice_acute_heroin_striatum", "Skupio_mice_acute_striatum")
j <- c("Lefevre_mice_NAc", "Seney_human_NAc", "Imperio_SS_rat_NAc", "Korostynski_mice_acute_morphine_striatum", 
       "Korostynski_mice_chronic_striatum", "Piechota_mice_acute_morphine_striatum", "Skupio_mice_chronic_striatum")
JI_intra <- c()
for(n in 1:length(i)) {JI_intra <- c(JI_intra, (jaccard_indexes[i[n], j[n]])*100)}

i <- rep(row.names(jaccard_indexes), seq(from = (length(row.names(jaccard_indexes))-1), to = 0, by = -1)); j <- c()
for(n in 2:length(row.names(jaccard_indexes))) {j <- c(j, row.names(jaccard_indexes)[n:length(row.names(jaccard_indexes))])}
JI_all <- c()
for(n in 1:length(i)) {JI_all <- c(JI_all, (jaccard_indexes[i[n], j[n]])*100)}
JI_inter <- JI_all
for (n in 1:length(JI_intra)) {JI_inter <- JI_inter[-which.max(JI_inter == JI_intra[n])]}

data.frame(jaccard_indexes = c(JI_intra, JI_inter), type = c(rep("intrastudy", length(JI_intra)), rep("interstudy", length(JI_inter)))) %>%
  ggplot(aes(x = type, y = jaccard_indexes)) + geom_boxplot()
t.test(x = JI_intra, y = JI_inter) # significant: p = 0.01104
```
