## Supplementary material for "Functional genomic mechanisms of opioid action and opioid use disorder: a systematic review of animal models and human studies": Code and data: Fig5E_timepoint.R

```
library(readxl)
library(biomaRt)
library(UpSetR)
library(dplyr)
library(ggplot2)

##########################################
# Figure 5D - Jaccard indexes comparison #
##########################################

# immediately: 2 (Townsend, Grice) > 1 JI
# 1h: 5 (Valderrama-Carjaval, Korostynski heroin, Korostynski morphine, Piechota heroin, Piechota morphine) > 10 JI
# 2h: 5 (Valderrama-Carjaval, Korostynski heroin, Korostynski morphine, Piechota heroin, Piechota morphine) > 10 JI
# 4h: 10 (Coffey, Skupio chronic, Skupio acute, Ye, Korostynski heroin, Korostynski morphine, Piechota morphine, Piechota heroin, Korostynski acute,     
#         Korostynski chronic) > 45 JI
# 8h: 4 (Korostynski heroin, Korostynski morphine, Piechota heroin, Piechota morphine) > 6 JI
# 24h: 3 (Liu, Imperio SS, Imperio LS) > 3 JI
# > 75 JI intra
# 30 studies > 406 JI all
# > 331 JI inter

Townsend_rat <- read_excel("DEG_Nac.xlsx", sheet = "Townsend_rat_males", skip = 1)
Townsend_rat <- (Townsend_rat %>% filter(pvalue < 0.05))$Gene.name
Grice_mice <- read_excel("DEG_Nac.xlsx", sheet = "Grice_mice", skip = 1)
# pvalue is at < 0.01 already, chip AFFYMETRIX MouseGene U74A was used (need to translate probes to genes)
mart <- useMart("ENSEMBL_MART_ENSEMBL")
mart <- useDataset("mmusculus_gene_ensembl", mart)
Grice <- unique((getBM(attributes = c("affy_mg_u74a", "external_gene_name"), filters = "affy_mg_u74a", 
                       values = Grice_mice$`probe set`, mart = mart))$external_gene_name); rm(Grice_mice)
Korostynski_2013 <- read_excel("DEG_striatum.xlsx", sheet = "Korostynski_acute_2013")
Korostynski_1h_morphine <- unique(getBM(attributes = c("ensembl_gene_id", "external_gene_name"), filters = "ensembl_gene_id", 
                                        values = na.omit(Korostynski_2013$`1h Morphine`), mart = mart)$external_gene_name)
Korostynski_2h_morphine <- unique(getBM(attributes = c("ensembl_gene_id", "external_gene_name"), filters = "ensembl_gene_id", 
                                        values = na.omit(Korostynski_2013$`2h Morphine`), mart = mart)$external_gene_name)
Korostynski_4h_morphine <- unique(getBM(attributes = c("ensembl_gene_id", "external_gene_name"), filters = "ensembl_gene_id", 
                                        values = na.omit(Korostynski_2013$`4h Morphine`), mart = mart)$external_gene_name)
Korostynski_8h_morphine <- unique(getBM(attributes = c("ensembl_gene_id", "external_gene_name"), filters = "ensembl_gene_id", 
                                        values = na.omit(Korostynski_2013$`8h Morphine`), mart = mart)$external_gene_name)
Korostynski_1h_heroin <- unique(getBM(attributes = c("ensembl_gene_id", "external_gene_name"), filters = "ensembl_gene_id", 
                                      values = na.omit(Korostynski_2013$`1h Heroin`), mart = mart)$external_gene_name)
Korostynski_2h_heroin <- unique(getBM(attributes = c("ensembl_gene_id", "external_gene_name"), filters = "ensembl_gene_id", 
                                      values = na.omit(Korostynski_2013$`2h Heroin`), mart = mart)$external_gene_name)
Korostynski_4h_heroin <- unique(getBM(attributes = c("ensembl_gene_id", "external_gene_name"), filters = "ensembl_gene_id", 
                                      values = na.omit(Korostynski_2013$`4h Heroin`), mart = mart)$external_gene_name)
Korostynski_8h_heroin <- unique(getBM(attributes = c("ensembl_gene_id", "external_gene_name"), filters = "ensembl_gene_id", 
                                      values = na.omit(Korostynski_2013$`8h Heroin`), mart = mart)$external_gene_name); rm(Korostynski_2013)
Piechota_mice_acute <- read_excel("DEG_striatum.xlsx", sheet = "Piechota_acute_2010")
Piechota_1h_morphine <- Piechota_mice_acute[Piechota_mice_acute$`1h Morphine` < 0.05,]$`Gene symbol`
Piechota_2h_morphine <- Piechota_mice_acute[Piechota_mice_acute$`2h Morphine` < 0.05,]$`Gene symbol`
Piechota_4h_morphine <- Piechota_mice_acute[Piechota_mice_acute$`4h Morphine` < 0.05,]$`Gene symbol`
Piechota_8h_morphine <- Piechota_mice_acute[Piechota_mice_acute$`8h Morphine` < 0.05,]$`Gene symbol`
Piechota_1h_heroin <- Piechota_mice_acute[Piechota_mice_acute$`1h Heroin` < 0.05,]$`Gene symbol`
Piechota_2h_heroin <- Piechota_mice_acute[Piechota_mice_acute$`2h Heroin` < 0.05,]$`Gene symbol`
Piechota_4h_heroin <- Piechota_mice_acute[Piechota_mice_acute$`4h Heroin` < 0.05,]$`Gene symbol`
Piechota_8h_heroin <- Piechota_mice_acute[Piechota_mice_acute$`8h Heroin` < 0.05,]$`Gene symbol`; rm(Piechota_mice_acute)
Coffey <- read_excel("DEG_striatum.xlsx", sheet = "Coffey")
Coffey <- (na.omit(Coffey)[na.omit(Coffey$`P-adj` < 0.1),])$GeneID
Skupio_chronic <- read_excel("DEG_striatum.xlsx", sheet = "Skupio_chronic")
Skupio_chronic <- unique(Skupio_chronic$Gene)
Skupio_acute <- read_excel("DEG_striatum.xlsx", sheet = "Skupio_acute")
Skupio_acute <- unique(Skupio_acute$Gene)
Ye <- read_excel("DEG_Nac.xlsx", sheet = "Ye_mice", skip = 0)
Ye <- Ye$gene_id
Korostynski_mice_chronic <- read_excel("DEG_striatum.xlsx", sheet = "Korostynski_chronic_2007")
Korostynski_mice_conv <- (getBM(attributes = c("affy_mouse430_2", "external_gene_name"), filters = "affy_mouse430_2", 
                                values = Korostynski_mice_chronic[!is.na(Korostynski_mice_chronic$`Probe set...2`),]$`Probe set...2`, mart = mart))
Korostynski_mice_conv <- rbind(Korostynski_mice_conv, (getBM(attributes = c("affy_mouse430_2", "external_gene_name"), filters = "affy_mouse430_2", 
                                                             values = Korostynski_mice_chronic[!is.na(Korostynski_mice_chronic$`Probe set...6`),]$`Probe set...6`, mart = mart)))
Korostynski_4h_chronic <- unique(Korostynski_mice_conv$external_gene_name)
Korostynski_mice_acute <- read_excel("DEG_striatum.xlsx", sheet = "Korostynski_acute_2007")
Korostynski_mice_conv <- (getBM(attributes = c("affy_mouse430_2", "external_gene_name"), filters = "affy_mouse430_2", 
                                values = Korostynski_mice_acute[!is.na(Korostynski_mice_acute$`Probe set...2`),]$`Probe set...2`, mart = mart))
Korostynski_mice_conv <- rbind(Korostynski_mice_conv, (getBM(attributes = c("affy_mouse430_2", "external_gene_name"), filters = "affy_mouse430_2", 
                                                             values = Korostynski_mice_acute[!is.na(Korostynski_mice_acute$`Probe set...6`),]$`Probe set...6`, mart = mart)))
Korostynski_4h_acute <- unique(Korostynski_mice_conv$external_gene_name); rm(Korostynski_mice_chronic, Korostynski_mice_acute, Korostynski_mice_conv)
# columns were only colored and we can't retrieve that in R so gene IDs were entered by hand
Valderrama_rats_1h_annot <- c("Kcnj6", "Mfsd4", "Neurod6", "Gpr165", "Sstr1", "Cpne4", "Trpc4", "Nr4a3", "Slc17a6", "Unc5d", "Unc5d", "Nrp2", "Cbln2", "Vsnl1", 
                              "Tmem178a", "Pnck", "Vsnl1", "Gfra1", "Gng2", "Atp2b4")
mart <- useMart("ENSEMBL_MART_ENSEMBL")
mart <- useDataset("rnorvegicus_gene_ensembl", mart)
# we only re-annotate probes for which no corresponding genes were provided in their study and keep their other annotations
Valderrama_rats_1h_conv <- (getBM(attributes = c("affy_rat230_2", "external_gene_name"), filters = "affy_rat230_2", 
                               values = c("1379445_at", "1391598_at", "1375043_at", "1390645_at"), mart = mart))
Valderrama_1h <- unique(c(Valderrama_rats_1h_annot, Valderrama_rats_1h_conv$external_gene_name)); rm(Valderrama_rats_1h_annot, Valderrama_rats_1h_conv)
Valderrama_rats_2h_annot <- c("Hspa1b", "Hspa1a", "Rsrp1", "Pdk4", "Hspb1", "Sc5d", "Ccdc117", "Dnajb1", "Bag3", "Jun", "Homer1")
Valderrama_rats_2h_conv <- (getBM(attributes = c("affy_rat230_2", "external_gene_name"), filters = "affy_rat230_2", 
                                  values = c("1378960_at", "1375388_at", "1391654_at", "1380306_at", "1375043_at"), mart = mart))
Valderrama_2h <- unique(c(Valderrama_rats_2h_annot, Valderrama_rats_2h_conv$external_gene_name)); rm(Valderrama_rats_2h_annot, Valderrama_rats_2h_conv)
Liu_rats <- read_excel("DEG_frontal_cortex.xlsx", sheet = "Liu_rat")
Liu_rats <- Liu_rats$`Gene Symbol`
Imperio_LS_rat <- read_excel("DEG_Nac.xlsx", sheet = "Imperio_rat")
Imperio_LS_rat <- (Imperio_LS_rat %>% filter(!is.na(`FC ([Large Suppressor] vs [Saline]) ( Sal vs Lrg Supp Fold change >= 1.25 )`)))$`Gene Symbol`
Imperio_SS_rat <- read_excel("DEG_Nac.xlsx", sheet = "Imperio_rat")
Imperio_SS_rat <- (Imperio_SS_rat %>% filter(!is.na(`FC ([Small Suppressor] vs [Saline]) (Small Supp vs Saline Fold change >= 1.25 )`)))$`Gene Symbol`

mart <- useMart("ENSEMBL_MART_ENSEMBL")
mart <- useDataset("rnorvegicus_gene_ensembl", mart)
Townsend <- unique((getBM(attributes = c("mmusculus_homolog_associated_gene_name", "external_gene_name"), 
                    filters = c("with_mmusculus_homolog", "external_gene_name"),
                    values = list(TRUE, Townsend_rat), mart = mart))$mmusculus_homolog_associated_gene_name); rm(Townsend_rat)
Valderrama_1h <- unique((getBM(attributes = c("mmusculus_homolog_associated_gene_name", "external_gene_name"), 
                          filters = c("with_mmusculus_homolog", "external_gene_name"),
                          values = list(TRUE, Valderrama_1h), mart = mart))$mmusculus_homolog_associated_gene_name)
Valderrama_2h <- unique((getBM(attributes = c("mmusculus_homolog_associated_gene_name", "external_gene_name"), 
                               filters = c("with_mmusculus_homolog", "external_gene_name"),
                               values = list(TRUE, Valderrama_2h), mart = mart))$mmusculus_homolog_associated_gene_name)
Liu <- unique((getBM(attributes = c("mmusculus_homolog_associated_gene_name", "external_gene_name"), 
                               filters = c("with_mmusculus_homolog", "external_gene_name"),
                               values = list(TRUE, Liu_rats), mart = mart))$mmusculus_homolog_associated_gene_name); rm(Liu_rats)
Imperio_LS <- unique((getBM(attributes = c("mmusculus_homolog_associated_gene_name", "external_gene_name"), 
                     filters = c("with_mmusculus_homolog", "external_gene_name"),
                     values = list(TRUE, Imperio_LS_rat), mart = mart))$mmusculus_homolog_associated_gene_name); rm(Imperio_LS_rat)
Imperio_SS <- unique((getBM(attributes = c("mmusculus_homolog_associated_gene_name", "external_gene_name"), 
                            filters = c("with_mmusculus_homolog", "external_gene_name"),
                            values = list(TRUE, Imperio_SS_rat), mart = mart))$mmusculus_homolog_associated_gene_name); rm(Imperio_SS_rat)


rm(mart)
# For this step, only the 29 lists of genes should be loaded, remove any other objects from the environment using the rm() command

# get all jaccard indexes
jaccard_indexes <- matrix(rep(NA, length(ls())*length(ls())), ncol = length(ls()), dimnames = list(ls(), ls()))
for (i in rownames(jaccard_indexes)) {
  for (j in colnames(jaccard_indexes)) {
    common <- sum(get(i) %in% get(j))
    jaccard_indexes[i,j] <- common/(length(unique(c(get(i), get(j)))))
  }
}

# characteristics of the studies used for grouping purposes in this step are described in the "study_characteristics.ods" file

# jaccard index inter vs intrastrains
studies_t0 <- c("Townsend", "Grice")
studies_t1 <- c("Valderrama_1h", "Korostynski_1h_morphine", "Korostynski_1h_heroin", "Piechota_1h_morphine", "Piechota_1h_heroin")
studies_t2 <- c("Valderrama_2h", "Korostynski_2h_morphine", "Korostynski_2h_heroin", "Piechota_2h_morphine", "Piechota_2h_heroin")
studies_t4 <- c("Coffey", "Skupio_acute", "Skupio_chronic", "Ye", "Korostynski_4h_morphine", "Korostynski_4h_heroin", "Piechota_4h_morphine", "Piechota_4h_heroin",
                "Korostynski_4h_chronic", "Korostynski_4h_acute")
studies_t8 <- c("Korostynski_8h_morphine", "Korostynski_8h_heroin", "Piechota_8h_morphine", "Piechota_8h_heroin")
studies_t24 <- c("Liu", "Imperio_SS", "Imperio_LS")

i <- rep(studies_t0, seq(from = (length(studies_t0)-1), to = 0, by = -1)); j <- c()
for(n in 2:length(studies_t0)) {j <- c(j, studies_t0[n:length(studies_t0)])}
i <- c(i, rep(studies_t1, seq(from = (length(studies_t1)-1), to = 0, by = -1)))
for(n in 2:length(studies_t1)) {j <- c(j, studies_t1[n:length(studies_t1)])}
i <- c(i, rep(studies_t2, seq(from = (length(studies_t2)-1), to = 0, by = -1)))
for(n in 2:length(studies_t2)) {j <- c(j, studies_t2[n:length(studies_t2)])}
i <- c(i, rep(studies_t4, seq(from = (length(studies_t4)-1), to = 0, by = -1)))
for(n in 2:length(studies_t4)) {j <- c(j, studies_t4[n:length(studies_t4)])}
i <- c(i, rep(studies_t8, seq(from = (length(studies_t8)-1), to = 0, by = -1)))
for(n in 2:length(studies_t8)) {j <- c(j, studies_t8[n:length(studies_t8)])}
i <- c(i, rep(studies_t24, seq(from = (length(studies_t24)-1), to = 0, by = -1)))
for(n in 2:length(studies_t24)) {j <- c(j, studies_t24[n:length(studies_t24)])}
JI_intra <- c()
for(n in 1:length(i)) {JI_intra <- c(JI_intra, (jaccard_indexes[i[n], j[n]])*100)}

i <- rep(row.names(jaccard_indexes), seq(from = (length(row.names(jaccard_indexes))-1), to = 0, by = -1)); j <- c()
for(n in 2:length(row.names(jaccard_indexes))) {j <- c(j, row.names(jaccard_indexes)[n:length(row.names(jaccard_indexes))])}
JI_all <- c()
for(n in 1:length(i)) {JI_all <- c(JI_all, (jaccard_indexes[i[n], j[n]])*100)}
JI_inter <- JI_all
for (n in 1:length(JI_intra)) {JI_inter <- JI_inter[-which.max(JI_inter == JI_intra[n])]}

data.frame(jaccard_indexes = c(JI_intra, JI_inter), type = c(rep("intratimepoints", length(JI_intra)), rep("intertimepoints", length(JI_inter)))) %>%
  ggplot(aes(x = type, y = jaccard_indexes)) + geom_boxplot()
t.test(x = JI_intra, y = JI_inter) # significant: p = 0.0009851
```
