## Supplementary material for "Functional genomic mechanisms of opioid action and opioid use disorder: a systematic review of animal models and human studies": Code and data: Fig7.R

```
# Code to reproduce Figure 7A-E and Supplementary Figures 2.A-B and 3.A-B
# Alba Caparros-Roissard

library(dplyr)
library(readxl)
library(biomaRt)
library(RRHO2)
library(ComplexHeatmap)
library(UpSetR)
library(stringr)
library(ggplot2)

#####################################################################################
# Supplementary Figure 2A - Seney both sexes vs Townsend female rats (Venn Diagram) #
#####################################################################################

Seney_human <- read_excel("DEG_Nac.xlsx", sheet = "Seney_human")
Seney_human <- Seney_human %>% filter(!is.na(coefficient) & !is.na(corrected.p))
Seney_human <- Seney_human %>% filter(!(coefficient == 0))
Seney_human <- Seney_human[!duplicated(Seney_human$gene_symbol),]
Townsend_rat <- Townsend_rat <- read_excel("DEG_Nac.xlsx", sheet = "Townsend_rat_females", skip = 1)
Townsend_rat <- Townsend_rat %>% filter(!is.na(log2FoldChange) & !is.na(pvalue))
Townsend_rat <- Townsend_rat %>% filter(!(log2FoldChange == 0))
Townsend_rat <- Townsend_rat[!duplicated(Townsend_rat$Gene.name),]

# Townsend is a study done on rats, we chose to find orthologs of rat > human genes for the Townsend et al study
mart <- useMart("ENSEMBL_MART_ENSEMBL")
mart <- useDataset("rnorvegicus_gene_ensembl", mart)
Townsend_conv <- getBM(attributes = c("hsapiens_homolog_associated_gene_name", "external_gene_name"), 
                       filters = c("with_hsapiens_homolog", "external_gene_name"),
                       values = list(TRUE, Townsend_rat$Gene.name), mart = mart)
Townsend_conv <- Townsend_conv[!duplicated(Townsend_conv$hsapiens_homolog_associated_gene_name),]
Townsend_conv <- Townsend_conv[!duplicated(Townsend_conv$external_gene_name),]
Townsend_rat <- Townsend_rat[Townsend_rat$Gene.name %in% Townsend_conv$external_gene_name,] %>% arrange(Gene.name)
Townsend_conv <- Townsend_conv[Townsend_conv$external_gene_name %in% Townsend_rat$Gene.name,]
Townsend_rat$Gene.name <- (Townsend_conv %>% arrange(external_gene_name))$hsapiens_homolog_associated_gene_name

Seney_human <- Seney_human[Seney_human$gene_symbol %in% Townsend_rat$Gene.name,]
Townsend_rat <- Townsend_rat[Townsend_rat$Gene.name %in% Seney_human$gene_symbol,]

Seney_genes <- (Seney_human %>% filter(corrected.p < 0.01 & abs(coefficient) > 0.26))$gene_symbol
Townsend_genes <- Townsend_rat[Townsend_rat$pvalue < 0.05,]$Gene.name
upset(fromList(list(Seney_human = Seney_genes, Townsend_rat = Townsend_genes)), nsets = 2, order.by = "freq", nintersects = NA, keep.order = TRUE)
fisher.test(matrix(data = c(sum(Seney_genes %in% Townsend_genes), sum(!(Seney_genes %in% Townsend_genes)), sum(!(Townsend_genes %in% Seney_genes)), 
                            dim(Townsend_rat)[1]-(sum(Seney_genes %in% Townsend_genes)+sum(!(Seney_genes %in% Townsend_genes))+sum(!(Townsend_genes %in% Seney_genes)))), 
                   ncol = 2, byrow = TRUE)) # p = 0.2489; JI = 7.45%

##################################################################
# Figure 7A - Seney women vs Townsend female rats (Venn Diagram) #
##################################################################

# this file contains Seney et al data that was re-analyzed to have differential expression analysis for both sexes separately, and includes a column for mice gene name
Seney_human <- read.table("Human_Mouse_data_DESeq2_Seney21_FC1_FDR10.csv", sep = ",", header = TRUE, row.names = 1)
Seney_human <- Seney_human %>% filter(!is.na(Log2.FC..OUD_vs_CONT_F_NAc.) & !is.na(P.value..OUD_vs_CONT_F_NAc.))
Seney_human <- Seney_human %>% filter(!(Log2.FC..OUD_vs_CONT_F_NAc. == 0))
Seney_human <- Seney_human[!duplicated(Seney_human$MGI_symbol),]
Townsend_rat <- Townsend_rat <- read_excel("DEG_Nac.xlsx", sheet = "Townsend_rat_females", skip = 1)
Townsend_rat <- Townsend_rat %>% filter(!is.na(log2FoldChange) & !is.na(pvalue))
Townsend_rat <- Townsend_rat %>% filter(!(log2FoldChange == 0))
Townsend_rat <- Townsend_rat[!duplicated(Townsend_rat$Gene.name),]

# since our Seney et al reanalysed data file contained mice gene names, we chose to find orthologs of rat > mice genes for the Townsend et al study
Townsend_conv <- getBM(attributes = c("mmusculus_homolog_associated_gene_name", "external_gene_name"), 
                       filters = c("with_mmusculus_homolog", "external_gene_name"),
                       values = list(TRUE, Townsend_rat$Gene.name), mart = mart)
Townsend_conv <- Townsend_conv[!duplicated(Townsend_conv$mmusculus_homolog_associated_gene_name),]
Townsend_conv <- Townsend_conv[!duplicated(Townsend_conv$external_gene_name),]
Townsend_rat <- Townsend_rat[Townsend_rat$Gene.name %in% Townsend_conv$external_gene_name,] %>% arrange(Gene.name)
Townsend_conv <- Townsend_conv[Townsend_conv$external_gene_name %in% Townsend_rat$Gene.name,]
Townsend_rat$Gene.name <- (Townsend_conv %>% arrange(external_gene_name))$mmusculus_homolog_associated_gene_name

Seney_human <- Seney_human[Seney_human$MGI_symbol %in% Townsend_rat$Gene.name,]
Townsend_rat <- Townsend_rat[Townsend_rat$Gene.name %in% Seney_human$MGI_symbol,]

Seney_genes <- Seney_human[Seney_human$P.value..OUD_vs_CONT_F_NAc. < 0.05,]$MGI_symbol
Townsend_genes <- Townsend_rat[Townsend_rat$pvalue < 0.05,]$Gene.name
upset(fromList(list(Seney_human = Seney_genes, Townsend_rat = Townsend_genes)), nsets = 2, order.by = "freq", nintersects = NA, keep.order = TRUE)
fisher.test(matrix(data = c(sum(Seney_genes %in% Townsend_genes), sum(!(Seney_genes %in% Townsend_genes)), sum(!(Townsend_genes %in% Seney_genes)), 
                            dim(Townsend_rat)[1]-(sum(Seney_genes %in% Townsend_genes)+sum(!(Seney_genes %in% Townsend_genes))+sum(!(Townsend_genes %in% Seney_genes)))), 
                   ncol = 2, byrow = TRUE)) # p = 0.2056; JI = 9.5%

Seney_genes_up <- Seney_human[Seney_human$P.value..OUD_vs_CONT_F_NAc. < 0.05 & Seney_human$Log2.FC..OUD_vs_CONT_F_NAc. > 0,]$MGI_symbol
Townsend_genes_up <- Townsend_rat[Townsend_rat$pvalue < 0.05 & Townsend_rat$log2FoldChange > 0,]$Gene.name
upset(fromList(list(Seney_human = Seney_genes_up, Townsend_rat = Townsend_genes_up)), nsets = 2, order.by = "freq", nintersects = NA, keep.order = TRUE)

Seney_genes_dn <- Seney_human[Seney_human$P.value..OUD_vs_CONT_F_NAc. < 0.05 & Seney_human$Log2.FC..OUD_vs_CONT_F_NAc. < 0,]$MGI_symbol
Townsend_genes_dn <- Townsend_rat[Townsend_rat$pvalue < 0.05 & Townsend_rat$log2FoldChange < 0,]$Gene.name
upset(fromList(list(Seney_human = Seney_genes_dn, Townsend_rat = Townsend_genes_dn)), nsets = 2, order.by = "freq", nintersects = NA, keep.order = TRUE)

###########################################################
# Figure 7C - Seney women vs Townsend female rats (RRHO2) #
###########################################################

# 10 435 total genes for RRHO2
list1_DDE <- c(-log10(Seney_human[Seney_human$Log2.FC..OUD_vs_CONT_F_NAc. < 0,]$P.value..OUD_vs_CONT_F_NAc.) * (-1),
               -log10(Seney_human[Seney_human$Log2.FC..OUD_vs_CONT_F_NAc. > 0,]$P.value..OUD_vs_CONT_F_NAc.))
gene_list1 <- data.frame(Genes=c(Seney_human[Seney_human$Log2.FC..OUD_vs_CONT_F_NAc. < 0,]$MGI_symbol, 
                                 Seney_human[Seney_human$Log2.FC..OUD_vs_CONT_F_NAc. > 0,]$MGI_symbol),DDE = list1_DDE, stringsAsFactors = FALSE)
gene_list1 <- gene_list1 %>% arrange(DDE)

list2_DDE <- c(-log10(Townsend_rat[Townsend_rat$log2FoldChange< 0,]$pvalue) * (-1),
               -log10(Townsend_rat[Townsend_rat$log2FoldChange > 0,]$pvalue))
gene_list2 <- data.frame(Genes=c(Townsend_rat[Townsend_rat$log2FoldChange < 0,]$Gene.name, 
                                 Townsend_rat[Townsend_rat$log2FoldChange > 0,]$Gene.name),DDE = list2_DDE, stringsAsFactors = FALSE)
gene_list2 <- gene_list2 %>% arrange(DDE)

RRHO_obj <- RRHO2_initialize(gene_list2, gene_list1, labels = c("Townsend (2021) Female Rats - Morphine Effect", "Seney (2021) Women - OUD Effect"), method = "hyper")
RRHO2_heatmap(RRHO_obj)

###################################################################################
# Supplementary Figure 2B - Seney both sexes vs Townsend male rats (Venn Diagram) #
###################################################################################

Seney_human <- read_excel("DEG_Nac.xlsx", sheet = "Seney_human")
Seney_human <- Seney_human %>% filter(!is.na(coefficient) & !is.na(corrected.p))
Seney_human <- Seney_human %>% filter(!(coefficient == 0))
Seney_human <- Seney_human[!duplicated(Seney_human$gene_symbol),]
Townsend_rat <- Townsend_rat <- read_excel("DEG_Nac.xlsx", sheet = "Townsend_rat_males", skip = 1)
Townsend_rat <- Townsend_rat %>% filter(!is.na(log2FoldChange) & !is.na(pvalue))
Townsend_rat <- Townsend_rat %>% filter(!(log2FoldChange == 0))
Townsend_rat <- Townsend_rat[!duplicated(Townsend_rat$Gene.name),]

# Townsend is a study done on rats, we chose to find orthologs of rat > human genes for the Townsend et al study
Townsend_conv <- getBM(attributes = c("hsapiens_homolog_associated_gene_name", "external_gene_name"), 
                       filters = c("with_hsapiens_homolog", "external_gene_name"),
                       values = list(TRUE, Townsend_rat$Gene.name), mart = mart)
Townsend_conv <- Townsend_conv[!duplicated(Townsend_conv$hsapiens_homolog_associated_gene_name),]
Townsend_conv <- Townsend_conv[!duplicated(Townsend_conv$external_gene_name),]
Townsend_rat <- Townsend_rat[Townsend_rat$Gene.name %in% Townsend_conv$external_gene_name,] %>% arrange(Gene.name)
Townsend_conv <- Townsend_conv[Townsend_conv$external_gene_name %in% Townsend_rat$Gene.name,]
Townsend_rat$Gene.name <- (Townsend_conv %>% arrange(external_gene_name))$hsapiens_homolog_associated_gene_name

Seney_human <- Seney_human[Seney_human$gene_symbol %in% Townsend_rat$Gene.name,]
Townsend_rat <- Townsend_rat[Townsend_rat$Gene.name %in% Seney_human$gene_symbol,]

Seney_genes <- (Seney_human %>% filter(corrected.p < 0.01 & abs(coefficient) > 0.26))$gene_symbol
Townsend_genes <- Townsend_rat[Townsend_rat$pvalue < 0.05,]$Gene.name
upset(fromList(list(Seney_human = Seney_genes, Townsend_rat = Townsend_genes)), nsets = 2, order.by = "freq", nintersects = NA, keep.order = TRUE)
fisher.test(matrix(data = c(sum(Seney_genes %in% Townsend_genes), sum(!(Seney_genes %in% Townsend_genes)), sum(!(Townsend_genes %in% Seney_genes)), 
                            dim(Townsend_rat)[1]-(sum(Seney_genes %in% Townsend_genes)+sum(!(Seney_genes %in% Townsend_genes))+sum(!(Townsend_genes %in% Seney_genes)))), 
                   ncol = 2, byrow = TRUE)) # p = 0.3492; JI = 4.11%

##############################################################
# Figure 7B - Seney men vs Townsend male rats (Venn Diagram) #
##############################################################

Seney_human <- read.table("Human_Mouse_data_DESeq2_Seney21_FC1_FDR10.csv", sep = ",", header = TRUE, row.names = 1)
Seney_human <- Seney_human %>% filter(!is.na(Log2.FC..OUD_vs_CONT_M_NAc.) & !is.na(P.value..OUD_vs_CONT_M_NAc.))
Seney_human <- Seney_human %>% filter(!(Log2.FC..OUD_vs_CONT_M_NAc. == 0))
Seney_human <- Seney_human[!duplicated(Seney_human$MGI_symbol),]
Townsend_rat <- Townsend_rat <- read_excel("DEG_Nac.xlsx", sheet = "Townsend_rat_males", skip = 1)
Townsend_rat <- Townsend_rat %>% filter(!is.na(log2FoldChange) & !is.na(pvalue))
Townsend_rat <- Townsend_rat %>% filter(!(log2FoldChange == 0))
Townsend_rat <- Townsend_rat[!duplicated(Townsend_rat$Gene.name),]

Townsend_conv <- getBM(attributes = c("mmusculus_homolog_associated_gene_name", "external_gene_name"), 
                       filters = c("with_mmusculus_homolog", "external_gene_name"),
                       values = list(TRUE, Townsend_rat$Gene.name), mart = mart)
Townsend_conv <- Townsend_conv[!duplicated(Townsend_conv$mmusculus_homolog_associated_gene_name),]
Townsend_conv <- Townsend_conv[!duplicated(Townsend_conv$external_gene_name),]
Townsend_rat <- Townsend_rat[Townsend_rat$Gene.name %in% Townsend_conv$external_gene_name,] %>% arrange(Gene.name)
Townsend_conv <- Townsend_conv[Townsend_conv$external_gene_name %in% Townsend_rat$Gene.name,]
Townsend_rat$Gene.name <- (Townsend_conv %>% arrange(external_gene_name))$mmusculus_homolog_associated_gene_name

Seney_human <- Seney_human[Seney_human$MGI_symbol %in% Townsend_rat$Gene.name,]
Townsend_rat <- Townsend_rat[Townsend_rat$Gene.name %in% Seney_human$MGI_symbol,]

Seney_genes <- Seney_human[Seney_human$P.value..OUD_vs_CONT_M_NAc. < 0.05,]$MGI_symbol
Townsend_genes <- Townsend_rat[Townsend_rat$pvalue < 0.05,]$Gene.name
upset(fromList(list(Seney_human = Seney_human[Seney_human$P.value..OUD_vs_CONT_M_NAc. < 0.05,]$MGI_symbol, 
                    Townsend_rat = Townsend_rat[Townsend_rat$pvalue < 0.05,]$Gene.name)), nsets = 2, order.by = "freq", nintersects = NA, keep.order = TRUE)
fisher.test(matrix(data = c(sum(Seney_genes %in% Townsend_genes), sum(!(Seney_genes %in% Townsend_genes)), sum(!(Townsend_genes %in% Seney_genes)), 
                            dim(Townsend_rat)[1]-(sum(Seney_genes %in% Townsend_genes)+sum(!(Seney_genes %in% Townsend_genes))+sum(!(Townsend_genes %in% Seney_genes)))), 
                   ncol = 2, byrow = TRUE)) # p = 0.3606; JI = 4.5%

Seney_genes_up <- Seney_human[Seney_human$P.value..OUD_vs_CONT_M_NAc. < 0.05 & Seney_human$Log2.FC..OUD_vs_CONT_M_NAc. > 0,]$MGI_symbol
Townsend_genes_up <- Townsend_rat[Townsend_rat$pvalue < 0.05 & Townsend_rat$log2FoldChange > 0,]$Gene.name
upset(fromList(list(Seney_human = Seney_genes_up, Townsend_rat = Townsend_genes_up)), nsets = 2, order.by = "freq", nintersects = NA, keep.order = TRUE)

Seney_genes_dn <- Seney_human[Seney_human$P.value..OUD_vs_CONT_M_NAc. < 0.05 & Seney_human$Log2.FC..OUD_vs_CONT_M_NAc. < 0,]$MGI_symbol
Townsend_genes_dn <- Townsend_rat[Townsend_rat$pvalue < 0.05 & Townsend_rat$log2FoldChange < 0,]$Gene.name
upset(fromList(list(Seney_human = Seney_genes_dn, Townsend_rat = Townsend_genes_dn)), nsets = 2, order.by = "freq", nintersects = NA, keep.order = TRUE)

#######################################################
# Figure 7D - Seney men vs Townsend male rats (RRHO2) #
#######################################################

# 10 162 total genes for RRHO2 - males
list1_DDE <- c(-log10(Seney_human[Seney_human$Log2.FC..OUD_vs_CONT_M_NAc. < 0,]$P.value..OUD_vs_CONT_M_NAc.) * (-1),
               -log10(Seney_human[Seney_human$Log2.FC..OUD_vs_CONT_M_NAc. > 0,]$P.value..OUD_vs_CONT_M_NAc.))
gene_list1 <- data.frame(Genes=c(Seney_human[Seney_human$Log2.FC..OUD_vs_CONT_M_NAc. < 0,]$MGI_symbol, 
                                 Seney_human[Seney_human$Log2.FC..OUD_vs_CONT_M_NAc. > 0,]$MGI_symbol),DDE = list1_DDE, stringsAsFactors = FALSE)
gene_list1 <- gene_list1 %>% arrange(DDE)

list2_DDE <- c(-log10(Townsend_rat[Townsend_rat$log2FoldChange< 0,]$pvalue) * (-1),
               -log10(Townsend_rat[Townsend_rat$log2FoldChange > 0,]$pvalue))
gene_list2 <- data.frame(Genes=c(Townsend_rat[Townsend_rat$log2FoldChange < 0,]$Gene.name, 
                                 Townsend_rat[Townsend_rat$log2FoldChange > 0,]$Gene.name),DDE = list2_DDE, stringsAsFactors = FALSE)
gene_list2 <- gene_list2 %>% arrange(DDE)

RRHO_obj <- RRHO2_initialize(gene_list2, gene_list1, labels = c("Townsend (2021) Male Rats - Morphine Effect", "Seney (2021) Men - OUD Effect"))
RRHO2_heatmap(RRHO_obj)

##################################################################
# Figure 7E - Single cell studies comparison of DEG by cell type #
##################################################################

Avey <- read_excel("single_cell_studies.xlsx", sheet = "Avey_sc_mice_DEG")
DE_Avey_cluster <- list()
for (i in seq(from = 1, to = 78, by = 3)) {
  DE_Avey_cluster[[colnames(Avey)[i]]] <- unname(unlist(Avey[,i]))[2:length(unname(unlist(Avey[,i])))][!is.na(unname(unlist(Avey[,i]))[2:length(unname(unlist(Avey[,i])))])]
}
Reiner <- read_excel("single_cell_studies.xlsx", sheet = "Reiner_sc_rat_DEG", skip = 1)
# since Avey et al was based on mice, we find orthologs rats > mice for genes in the Reiner et al study
DE_Reiner_cluster_tr_mice <- list()
library(biomaRt)
mart <- useMart("ENSEMBL_MART_ENSEMBL")
mart <- useDataset("rnorvegicus_gene_ensembl", mart)
for (cluster in unique(Reiner$`Cluster ID`)) {
  cluster_genes <- (Reiner %>% filter(`Cluster ID` == cluster))$Gene
  cluster_tr_mice <- unique(getBM(attributes = c("mmusculus_homolog_associated_gene_name", "external_gene_name"),
                                  filters = c("with_mmusculus_homolog", "external_gene_name"),
                                  values = list(TRUE, cluster_genes), mart = mart)$mmusculus_homolog_associated_gene_name)
  DE_Reiner_cluster_tr_mice[[cluster]] <- cluster_tr_mice
}

jaccard_indexes_DEG <- matrix(rep(NA, length(DE_Avey_cluster)*length(DE_Reiner_cluster_tr_mice)), ncol = length(DE_Reiner_cluster_tr_mice), 
                          dimnames = list(names(DE_Avey_cluster), names(DE_Reiner_cluster_tr_mice)))
for (i in rownames(jaccard_indexes_DEG)) {
  for (j in colnames(jaccard_indexes_DEG)) {
    common <- sum(DE_Avey_cluster[[i]] %in% DE_Reiner_cluster_tr_mice[[j]])
    jaccard_indexes_DEG[i,j] <- common/(length(unique(c(DE_Avey_cluster[[i]], DE_Reiner_cluster_tr_mice[[j]]))))
  }
}

Heatmap(jaccard_indexes_DEG)

####################################################################################################################
# Supplementary Figure 3A - Single cell studies comparison of markers (genes used to define clusters) by cell type #
####################################################################################################################

Avey <- read_excel("single_cell_studies.xlsx", sheet = "Avey_sc_mice_markers")
marker_Avey_cluster <- list()
for (cluster in unique(Avey$`Cluster Name`)) {
  marker_Avey_cluster[[cluster]] <- (Avey %>% filter(`Cluster Name` == cluster))$GENE
}
Reiner <- read_excel("single_cell_studies.xlsx", sheet = "Reiner_sc_rat_markers", skip = 1)
marker_Reiner_cluster_tr_mice <- list()
for (cluster in Reiner$Cluster) {
  marker <- unlist(unname((Reiner %>% filter(Cluster == cluster))))[2:dim(Reiner)[2]]
  marker_Reiner_cluster_tr_mice[[cluster]] <- unique(getBM(attributes = c("mmusculus_homolog_associated_gene_name", "external_gene_name"),
                                                           filters = c("with_mmusculus_homolog", "external_gene_name"),
                                                           values = list(TRUE, str_replace(marker[marker != ""], " ", "")), mart = mart)$mmusculus_homolog_associated_gene_name)
}

jaccard_indexes_marker <- matrix(rep(NA, length(marker_Avey_cluster)*length(marker_Reiner_cluster_tr_mice)), ncol = length(marker_Reiner_cluster_tr_mice), 
                          dimnames = list(names(marker_Avey_cluster), names(marker_Reiner_cluster_tr_mice)))
for (i in rownames(jaccard_indexes_marker)) {
  for (j in colnames(jaccard_indexes_marker)) {
    common <- sum(marker_Avey_cluster[[i]] %in% marker_Reiner_cluster_tr_mice[[j]])
    jaccard_indexes_marker[i,j] <- common/(length(unique(c(marker_Avey_cluster[[i]], marker_Reiner_cluster_tr_mice[[j]]))))
  }
}
Heatmap(jaccard_indexes_marker)

################################################################
# Supplementary Figure 3B - Correlation of JI, markers vs DEGs #
################################################################

jaccard_indexes_DEG <- jaccard_indexes_DEG[rownames(jaccard_indexes_DEG) %in% rownames(jaccard_indexes_marker), colnames(jaccard_indexes_DEG) %in% 
                                             colnames(jaccard_indexes_marker)]
jaccard_indexes_marker <- jaccard_indexes_marker[rownames(jaccard_indexes_marker) %in% rownames(jaccard_indexes_DEG), colnames(jaccard_indexes_marker) %in% 
                                                   colnames(jaccard_indexes_DEG)]
jaccard_indexes_marker <- jaccard_indexes_marker[rownames(jaccard_indexes_DEG), colnames(jaccard_indexes_DEG)]

data.frame(JI_DEG = as.vector(jaccard_indexes_DEG), JI_marker = as.vector(jaccard_indexes_marker)) %>% ggplot(aes(x = JI_DEG, y = JI_marker)) +
  geom_point() + geom_smooth(method = "lm")
cor.test(x = as.vector(jaccard_indexes_DEG), y = as.vector(jaccard_indexes_marker), method = "pearson") # cor = 0.02678277, p = 0.8233
```
