## Supplementary material for "Functional genomic mechanisms of opioid action and opioid use disorder: a systematic review of animal models and human studies": Code and data: FigS2.R

```
# Code to reproduce Figure S2
# Alba Caparros-Roissard

library(stringr)
library(dplyr)
library(ggplot2)

# this file contains information regarding whether a DEG down-regulated or up-regulated in a given study, this process was not automated
up_or_down_deg <- read.csv("DEG up or down, comparison.csv", sep = ";")[1:5]
order_genes <- c("Polr3e", "Itgad", "Arrdc2", "Rasd1", "Plekhf1", "Hif3a", "Ddit4", "Cldn5", "Tnfrsf25", "Homer1", "Cyp2d22", "Cables1", "Slc6a11", "Rprml", "Pdzd2", "Npy", 
           "Irs1", "Irf2bpl", "Enho", "Dbp", "Cryab", "Camk1g", "Txnip", "Tfcp2l1", "Rasl11a", "Mcam", "Mat2a", "Angptl4", "Pnpla2", "Phyhd1", "Oscp1", "Opalin", "Net1", 
           "Heph", "Fn1", "Dusp6", "Cx3cr1", "Kcng1", "Hmgcs1", "Gjb6", "Fam163a", "Fam107a", "Etnppl", "Emx2", "Avil", "Xdh", "Spsb1", "Sgk1", "Pdk4", "Map3k6", "Errfi1", 
           "Dusp1", "Tsc22d3", "Pla2g3", "Arid5b", "Wscd1", "Plin4", "Fkbp5", "Ccdc117", "Arrdc3", "Sult1a1", "Slc2a1", "Cdkn1a")
prop_agreement <- c(rep("other", 8), rep("50%", 4), rep("60%", 10), rep("66%", 6), rep("75%", 9), rep("80%", 8), rep("100%", 18))
names(prop_agreement) <- order_genes
df <- data.frame(gene = factor(rep(up_or_down_deg$Gene, each = 24), levels = order_genes), 
                 prop_agreement = factor(rep(prop_agreement[up_or_down_deg$Gene], each = 24), 
                                         levels = c("100%", "80%", "75%", "66%", "60%", "50%", "other")), 
                 species = rep(c("Mice","Rat","Rat","Rat","Human","Human","Human","Mice","Rat","Rat","Mice","Human","Rat","Mice","Mice","Mice","Mice","Mice","Mice",
                                 "Mice","Mice","Mice","Mice","Mice"), 63),
                 regions = rep(c(rep("Dorsal Striatum", 2), rep("Frontal Cortex", 4), rep("Nac", 8), rep("Striatum", 10)), 63), 
                 study = rep(c("Lefevre DS","Valderrama", "Kuntz-Melcavage", "Liu", "Seney FC", "Mendez", "Albertson", "Grice", "Imperio SS", "Imperio LS", "Lefevre Nac",
                               "Seney Nac", "Townsend", "Ye", "Coffey", "Korostynski acute", "Korostynski chronic", "Korostynski morphine", "Korostynski heroin", 
                               "Piechota chronic", "Piechota morphine", "Piechota heroin", "Skupio chronic", "Skupio acute"), 63), 
                 direction = rep(NA, 63*24))
dir <- c()
for (gene in unique(df$gene)) {
  for (study in unique(df$study)) {
    position <- str_detect((up_or_down_deg %>% filter(Gene == gene))[3:5], study)
    if (sum(position) == 0) {
      dir <- c(dir, NA)
    } else {
      if (position[1] == TRUE) {
        dir <- c(dir, "up")
      } else {
        if (position[2] == TRUE) {
          dir <- c(dir, "down")
        } else {
          if (position[3] == TRUE) {
            dir <- c(dir, "none")
          }  
        }
      }
    }
  }
}
df$direction <- dir

# by species
ggplot(df, aes(study, gene, fill = factor(direction))) + geom_tile(colour = "black") + 
  scale_fill_manual(values = c("dodgerblue3", "white", "firebrick2"), name = "Direction of DEG", labels = c("Down", "No direction provided", "Up", "Not DEG")) + 
  facet_grid(cols = vars(species), rows = vars(prop_agreement), scales = "free", space = "free") + theme(axis.text.x = element_text(angle = 45, hjust = 1)) +
  labs(y = "Genes", x = "Studies")

# by region (which we chose for Fig S2)
ggplot(df, aes(study, gene, fill = factor(direction))) + geom_tile(colour = "black") + 
  scale_fill_manual(values = c("dodgerblue3", "white", "firebrick2"), name = "Direction of DEG", labels = c("Down", "No direction provided", "Up", "Not DEG")) + 
  facet_grid(cols = vars(regions), rows = vars(prop_agreement), scales = "free", space = "free") + theme(axis.text.x = element_text(angle = 45, hjust = 1)) +
  labs(y = "Genes", x = "Studies")
```
