## Supplementary material for "Functional genomic mechanisms of opioid action and opioid use disorder: a systematic review of animal models and human studies": Code and data: README.txt

We split the code used to generate figures in several files:
 - "Fig4.R": code to reproduce figure 4.B-F
 - "Fig5.R": code to reproduce figure 5.A-D and supplementary figure 1.A-B
 - "Fig5E_timepoint.R": code to reproduce the "timepoint of last exposure" comparison in figure 5.E
 - "Fig6.R": code to reproduce figure 6
 - "Fig7.R": code to reproduce figure 7.A-E and supplementary figure 2.A-B and 3.A-B
 
All files necessary to reproduce the analysis are included in this folder

We used the following tables to constitute lists of differentially expressed genes:
 - Albertson et al, 2006: Supplemental Table 1
 - Seney et al, 2021: Data file S3 (Nucleus accumbens) Data file S2 (Frontal cortex)
 - Lefevre et al, 2020: Table S3, sheet "Control vs. MOR-SAL
 - Grice et al, 2007: Table S2, "B.Genes showing main effect of treatment"
 - Ye et al, 2014: Table S2
 - Imperio et al, 2016: Supplemental Table 2, "Large Suppressor vs Saline" and "Small Suppressor vs Saline"
 - Townsend et al, 2021: Supplementary Data 2, "NAc Fentanyl vs. Saline"
 - Valderrama-Carjaval et al, 2018: Supplementary Table S2, "Morphine vs Control"
 - Piechota et al, 2010: Additional file 2, sheet "drug_vs_saline", we selected columns regarding heroin and morphine
 - Korostynski et al, 2007: Additional file 2, sheet "C57BL6"
 - Korostynski et al, 2013: Genes lists were extracted from their website "http://genes2mind.org/gwt/searchforsignature.htm"
 - Skupio et al, 2017: Supplementary Table S2
 - Coffey et al, 2020: Input data was downloaded from https://github.com/DrCoffey/Morphine-Microglia-Manuscript/blob/main/Figure%204%20WGCNA/Data%20Sheets/DESeq2_5_Morphine%20Vehicle_Saline%20Vehicle_Input.xlsx
 - Piechota et al, 2012: Table 1, we only extracted genes which had highest fold change for "HER treatment"
 - Mendez et al, 2021: Supplementary Table 3
 - Ammon et al, 2003: Table 2
 - Ammon-Treiber et al, 2004: Table II
 - Kuntz-Melcavage et al, 2009: Additional file 1
 - Liu et al, 2021: Supplement 4
