## Supplementary material for "Functional genomic mechanisms of opioid action and opioid use disorder: a systematic review of animal models and human studies": Code and data: Seney_GEA.R

```
##################################################################################################################################
# Author: Charles DECRAENE
# input #1: a .csv file corresponding to the raw counts of the study. rows=genes (Ensembl gene ID) and columns=samples
# input #2: a .csv file corresponding to a description file (descr) with all information about the samples
##################################################################################################################################

descr <- read.csv(file="descr_GSE174409.csv", h=T, sep=",")
rownames(descr) <- descr$Sample_ID

input_rawCount <- read.csv(file="GSE174409_raw_counts.csv", h=T, sep=",")

# input_rawCount rownames (=ensembl ID) ---------------------------------------------------------------------------------------------
rownames(input_rawCount) <- input_rawCount$X
input_rawCount <- input_rawCount[, -1]

# (QC) Number and name of libraries in the initial project ----------------------------------------------------------------------------
nlib <- dim(descr)[1] # Number of samples
# Number of samples as well as their names are the same between initial project and raw counts provided > we move on

# (QC) suppress genes not expressed in all samples (input_rawCount becomes rawCount) -----------------------------------------------------
rawCount <- input_rawCount[apply(input_rawCount,1,sum)!=0,]; rm(input_rawCount)

# Convert data.frame (rawCount) to matrix (mat_rawCount)
mat_rawCount <- as.matrix(rawCount); rm(rawCount)

# --------------------------------------------------------------------------------------------------------------------------------
# Data annotations
# --------------------------------------------------------------------------------------------------------------------------------

library(biomaRt)
ensembl <- useMart("ensembl", dataset="hsapiens_gene_ensembl")
annotation <- getBM(attributes=c("ensembl_gene_id", "description", "ensembl_transcript_id","exon_chrom_start","exon_chrom_end"), 
                    filters="ensembl_gene_id", values=rownames(mat_rawCount), mart=ensembl)
# exon length
ensinfos.exlen <- data.frame(annotation$ensembl_gene_id, annotation$ensembl_transcript_id, abs(annotation$exon_chrom_start - annotation$exon_chrom_end)+1)
colnames(ensinfos.exlen) <- c("ensembl_gene_id", "ensembl_transcript_id", "exon_length")
# transcript length
tlen <- tapply(ensinfos.exlen$exon_length, ensinfos.exlen$ensembl_transcript_id, sum)
tlen.gene <- merge(tlen, unique(ensinfos.exlen[,1:2]), by.x="row.names", by.y="ensembl_transcript_id"); rm(ensinfos.exlen, tlen)
colnames(tlen.gene) <- c("ensembl_transcript_id", "transcript_length","ensembl_gene_id")
# gene length = median of the size of all transcripts corresponding to this gene
glen <- tapply(tlen.gene$transcript_length, tlen.gene$ensembl_gene_id, median); rm(tlen.gene)
# data with gene length
datalen <- merge(mat_rawCount, glen, by="row.names"); rm(glen)
colnames(datalen) <- c("ensembl_gene_id",colnames(mat_rawCount), "Gene_length")
# data with annotations and gene length
annotationgene <- unique(annotation[,1:2]); rm(annotation)
dataannot <- merge(datalen, annotationgene, by.x="ensembl_gene_id", by.y="ensembl_gene_id"); rm(datalen, annotationgene)

# to keep only the first part of the gene description (before [)
tmpdesc <- strsplit(as.character(dataannot$description),"[", fixed=T)
f <- function(l){
  if (length(l)>=1){
    return(l[[1]])
  }
  else{
    return("")
  }
}
tmpdescok <- unlist(lapply(tmpdesc, f))
dataannot$description <- tmpdescok; rm(tmpdesc, tmpdescok, f)

# data + annotations (+ rpkn + rpkn_max if necessary)
nbcol <- ncol(dataannot)
dataall <- data.frame(dataannot[,-c(nbcol,nbcol-1)], dataannot[,c(nbcol-1,nbcol)]); rm(dataannot, nbcol)
# dataall <- dataannot

#adding entrez_gen_id et hgnc_symbol
entrez_hgnc <- getBM(attributes=c("ensembl_gene_id", "entrezgene_id", "hgnc_symbol"), filters="ensembl_gene_id", values=rownames(mat_rawCount), mart=ensembl)
dataall.entrez_hgnc <- merge(entrez_hgnc, dataall, by.x="ensembl_gene_id", by.y="ensembl_gene_id"); rm(entrez_hgnc, dataall, ensembl, mat_rawCount)

# removing duplicates
doublons <- which(duplicated(dataall.entrez_hgnc$ensembl_gene_id))
final <- dataall.entrez_hgnc[-doublons,]; rm(dataall.entrez_hgnc, doublons)

finalrawCount_DESeq2 <- final[, c(1:(nlib+3))]
annot_study <- final[ , -c(4:(nlib*3+3))]; rm(final, nlib)

################################################################################
################   GENE EXPRESSION ANALYSIS (GEA)   ############################
################################################################################

# Creating coldata from descr
coldata <- descr; rm(descr)

# creating the input data.frame for GEA (input_GEA)
input_GEA <- finalrawCount_DESeq2; rm(finalrawCount_DESeq2)
rownames(input_GEA) <- input_GEA$ensembl_gene_id
input_GEA <- input_GEA[, -c(1:3)] # only rownames and raw counts

# Creating a DESeqDataSet
rawcount <- input_GEA; rm(input_GEA)
condition_ALL <- coldata; rm(coldata)
condition_ALL$diagnosis <- as.factor(condition_ALL$diagnosis)
condition_ALL$sex <- as.factor(condition_ALL$sex)
condition_ALL$source_name <- as.factor(condition_ALL$source_name)
condition_ALL$tobaccoatod <- as.factor(condition_ALL$tobaccoatod)
condition_ALL$Sample_ID <- as.factor(condition_ALL$Sample_ID)
condition_ALL$group <- as.factor(condition_ALL$group)
condition_ALL$group1 <- as.factor(condition_ALL$group1)
condition <- condition_ALL; rm(condition_ALL)

# Use the DESeqDataSetFromMatrix to create a DESeqDataSet object
library(DESeq2)
dds0 <- DESeqDataSetFromMatrix(countData = rawcount, colData = condition, design = ~ diagnosis); rm(rawcount)
dds0 <- DESeq(object=dds0, betaPrior = F)

# Here we perform a minimal pre-filtering 
# to keep only rows that have at least 10 reads total
dds.ALL <- dds0; rm(dds0)
keep <- rowSums(counts(dds.ALL)) >= 10
dds.filtered <- dds.ALL[keep,]; rm(dds.ALL, keep)

# --------------------------------------------------------------------------------------------------------------------------------
# unnormalized and normalized data
# --------------------------------------------------------------------------------------------------------------------------------

normlzd_dds <- counts(dds.filtered, normalized=T)
un_normlzd_dds <- counts(dds.filtered, normalized=F)
df_normlzd_dds <- as.data.frame(normlzd_dds); rm(normlzd_dds)

# RAW COUNTS for GEA
input_GEA <- un_normlzd_dds; rm(un_normlzd_dds)

# Checking the normalization
library(affy)
while (!is.null(dev.list())) dev.off()
epsilon <- 1 # pseudo-count to avoid problems with log(0)
par(mfrow=c(2,2),cex.lab=0.7)
boxplot(log2(counts(dds.filtered, normalized=F)+ epsilon),  #col=coldata$diagnosis,
        cex.axis=0.7,
        las=1, xlab="log2(counts)", horizontal=TRUE, main="Raw counts")

boxplot(log2(counts(dds.filtered, normalized=T) + epsilon),  #col=coldata$diagnosis,
        cex.axis=0.7,
        las=1, xlab="log2(normalized counts)", horizontal=TRUE, main="Normalized counts")

plotDensity(log2(counts(dds.filtered, normalized=F) + epsilon),  #col=coldata$diagnosis,
            xlab="log2(counts)", cex.lab=0.7, panel.first=grid())

plotDensity(log2(counts(dds.filtered, normalized=T) + epsilon), #col=coldata$diagnosis,
            xlab="log2(normalized counts)", cex.lab=0.7, panel.first=grid()); rm(epsilon)

data_annot <- annot_study; rm(annot_study)
rownames(data_annot) <- data_annot$ensembl_gene_id
data_annot <- data_annot[rownames(assay(dds.filtered)),]; rm(dds.filtered)

df_normlzd_dds$Gene <- data_annot$hgnc_symbol
df_normlzd_dds$Entrez <- data_annot$entrezgene_id
df_normlzd_dds$Ensembl <- data_annot$ensembl_gene_id

# DESeq2 ANALYSIS

# Comparisons of interest were performed using the Wald test
# for differential expression proposed by Love et al. (2015) and
# implemented in the Bioconductor package DESeq2

# Conditions to compare
A = "OUD"
B = "CONT"

# Comparisons
compM_N <- paste(A, "_vs_", B,"_M_NAc", sep = "")
compF_N <- paste(A, "_vs_", B,"_F_NAc", sep = ""); rm(A, B)

# --------------------------------------------------------------------------------------------------------------------------------
# CONTRAST ANALYSIS
# --------------------------------------------------------------------------------------------------------------------------------

## MALES ##
condition_M_NAC <- subset(condition, sex == "male" & source_name == "NAC")
list_M_NAC <- rownames(condition_M_NAC)
input_M_NAC <- input_GEA[,list_M_NAC]; rm(list_M_NAC)

dds_M_NAC <- DESeqDataSetFromMatrix(countData=input_M_NAC, colData=condition_M_NAC, design=~group1); rm(condition_M_NAC, input_M_NAC)
dds_M_NAC <- DESeq(object=dds_M_NAC, betaPrior=F)

dds <- dds_M_NAC; rm(dds_M_NAC)
res <- results(dds, contrast=c("group1", "OUD_male_NAC", "CONT_male_NAC"))
res$Gene <- df_normlzd_dds$Gene
res$Entrez <- df_normlzd_dds$Entrez
res$Ensembl <- df_normlzd_dds$Ensembl
res_OUDvsCONT_NAC_M <- res

## FEMALES ##
condition_F_NAC <- subset(condition, sex == "female" & source_name == "NAC"); rm(condition)
list_F_NAC <- rownames(condition_F_NAC)
input_F_NAC <- input_GEA[,list_F_NAC]; rm(input_GEA, list_F_NAC)

dds_F_NAC <- DESeqDataSetFromMatrix(countData=input_F_NAC, colData=condition_F_NAC, design=~group1); rm(condition_F_NAC, input_F_NAC)
dds_F_NAC <- DESeq(object=dds_F_NAC, betaPrior=F)

dds <- dds_F_NAC; rm(dds_F_NAC)
res <- results(dds, contrast=c("group1", "OUD_female_NAC", "CONT_female_NAC")); rm(dds)
res$Gene <- df_normlzd_dds$Gene
res$Entrez <- df_normlzd_dds$Entrez
res$Ensembl <- df_normlzd_dds$Ensembl
res_OUDvsCONT_NAC_F <- res

##################################################################################################################################
# Exporting data as a final table
##################################################################################################################################

library(tidyr)
library(dplyr)

comp <- compM_N; rm(compM_N)
res <- res_OUDvsCONT_NAC_M; rm(res_OUDvsCONT_NAC_M)
res <- as.data.frame(res[ , c('log2FoldChange', 'pvalue', 'padj','stat'), drop=FALSE])
colnames(res) <- c(paste("Log2 FC (", comp, ")", sep = ""),
                   paste("P-value (", comp, ")", sep = ""),
                   paste("Adjusted p-value (", comp, ")", sep = ""),
                   paste("Stat (", comp, ")", sep = ""))
dfres <- res

comp <- compF_N; rm(compF_N)
res <- res_OUDvsCONT_NAC_F; rm(res_OUDvsCONT_NAC_F)
res <- as.data.frame(res[ , c('log2FoldChange', 'pvalue', 'padj','stat'), drop=FALSE])
colnames(res) <- c(paste("Log2 FC (", comp, ")", sep = ""),
                   paste("P-value (", comp, ")", sep = ""),
                   paste("Adjusted p-value (", comp, ")", sep = ""),
                   paste("Stat (", comp, ")", sep = "")); rm(comp)
dfres <- cbind(dfres, res); rm(res)

count <- df_normlzd_dds; rm(df_normlzd_dds)
count.res <- merge(count, dfres, by.x="Ensembl", by.y="row.names"); rm(count, dfres)
count.res <- count.res %>% relocate(Ensembl, .before = HN1)
colnames(count.res)

count.res <- count.res %>% relocate(Gene, .after = Ensembl)
count.res <- count.res %>% relocate(Entrez, .after = Gene)

# Human to mouse gene names
human <- useMart(host="https://dec2021.archive.ensembl.org","ensembl", dataset = "hsapiens_gene_ensembl")
mouse <- useMart(host="https://dec2021.archive.ensembl.org","ensembl", dataset = "mmusculus_gene_ensembl")

genesV2 <- getLDS(attributes = c("hgnc_symbol"), filters = "hgnc_symbol", values = data_annot$hgnc_symbol , mart = human, attributesL = c("mgi_symbol"), 
                  martL = mouse, uniqueRows=T); rm(data_annot, human, mouse)

Human_Mouse <- merge(count.res,genesV2,by.x= "Gene", by.y = "HGNC.symbol"); rm(count.res, genesV2)
Human_Mouse$MGI_symbol <- Human_Mouse$MGI.symbol

# Final output .csv file
write.csv(x=Human_Mouse, file=paste("../Human_Mouse_data_DESeq2_Seney21_FC1_FDR10.csv", sep=""))
```
